# Public exams decrease anxiety and facilitate deeper conceptual thinking

**DOI:** 10.1101/2022.04.15.488479

**Authors:** Benjamin Wiggins, Leah Lily, Carly Busch, Meta Landys, J. Gwen Shlichta, Tianhong Shi, Tandi Ngwenyama

**Affiliations:** Shoreline Community College; University of Washington, Western Washington University; Arizona State University; Oregon State University; Edmonds Community College

## Abstract

Assessment methods across post-secondary STEM education are traditionally constrained by logistics and likely contribute to the widespread inequities in STEM education outcomes. As part of attempts to improve and diversify the methods used in assessment, the authors have developed a flexible and low-tech style known as ‘public exams’ based in educational best practices. Public exams attempt to authentically involve students into the process of assessment through the use of pre-released portions of the exam. Using mixed-methods research techniques at a closely matched pair of institutions (a research-intensive university and a community college classroom), we observed signals of positive impact from the public exam on student learning experiences. Public exams appear to promote deeper thought, to direct students more efficiently to the core concepts in the discipline, and to decrease student anxiety around exams. The public exam experience does not show evidence for exacerbating gaps in exam outcomes for students from underrepresented backgrounds. This suggests that public exams are worth deeper investigation as an evidence-based and effective assessment style.

## Introduction

High-stakes examination-based assessments (hereafter, exams) are a common and widespread feature of postsecondary education (Stobart & Eggen, 2012). Whether used to give formative feedback to students, to summatively assess students’ knowledge, to create selection barriers for capacity-constrained programs or careers, or simply to assign grades for external use, these exams are complex structural elements that students must grapple with (Wideen et al., 1997). Problematically, the educational practices used widely in college and universities today are often based in traditional routines and logistical concerns instead of evidence-based, student-centered practices (Ambrose, 2010; Chase, 2020). Improving the practices of giving and taking exams has the potential to improve educational experiences for a more diverse, deeper, and more talented pool of future students (Intemann, 2009; Ralph et al., 2022).

In our own classrooms, we struggled to develop traditional exams that met student needs well. The frictions and problematic points in our exams reappeared consistently in student feedback and seemed to align with problems discussed in education literature. Within the highly unequal power relationship between students and their STEM faculty, students have little to no agency about their own assessment (Metevier et al., 2022). For students who are new to college practices, they are figuring out the ‘rules of the game’; those rules can be unpredictable and highly consequential (Canning et al., 2020). Multilingual learners in monolingual classrooms face high-stakes challenges during an exam (Sibanda, 2021). Anxiety around education can be exacerbated by exams and this anxiety tends to affect groups of students unjustly (Cooper et al., 2018; Jenkins et al., 2021). Whether in stressed conversations or anonymous evaluations, students repeatedly expressed problematic experiences with the exams we constructed. The ‘public exam’ style that follows is the result of our efforts to design a better system for delivering assessment challenges that still logistically fit within the constraints and pressures of a full teaching load (Rossing & Lavitt, 2016).

Our solution is an interrelated set of evidence-based practices that we collectively describe as the ‘public exam’ style. The traditional style for postsecondary STEM education is to reveal assessment tasks to students during the exam itself (Roberts et al., 2021; Zhuang et al., 2019). Ongoing work to redesign traditional exams includes efforts such as competency-based grading and other modes of producing formative and summative assessments in college STEM (DeCoito & Estaiteyeh, 2022; Diegelman-Parente, 2011; Gao et al., 2020; Kitchen et al., 2006; Lynd-Balta, 2006; Reynders et al., 2020; Stanger-Hall, 2012). In this work, we use the lens of educative assessment to re-design our exams to be more educative and transparent. Educative assessment can be summarized by saying that assessments have many purposes but the primary goal among them should be as a tool for facilitating student learning (Buxton et al., 2013; Fink, 2003; Jönsson, 2008; G. Wiggins, 1998, 2011). We chose this educative lens to ensure that our exams were helping to meet our larger learning goals in a system where other goals like summative grades are often strongly prioritized institutionally. Specifically, educative assessment suggests that educators can create challenging exams for students that are useful practice for their careers and lives such that teaching directly to these exams will be worthwhile (Erduran & Wooding, 2021). This increased focus on transparency in assessment is in contrast with traditional methods and likely to improve student outcomes (Felten & Finley, 2019; Harsma et al., 2021).

### What is a public exam?

The public exam style is the result of our attempts to re-design exams to engender trust and authentic engagement between students and instructors (Black & Wiliam, 1998; Driessen et al., 2022). Throughout the manuscript, we use the term “authentic involvement” to refer to stylistic elements of public exams that address this overarching theme relating to trust (Brown, 2017; D’Ambrosio, 2021; Felten & Finley, 2019). The three evidence-based elements of public exams described below are frequently addressed throughout K-12 education and are useful in convincing students that the assessment process can work for them (Darling-Hammond & Bransford, 2005; R. Keith. Sawyer, 2005; Zeichner et al., 2000). These three stylistic elements of public exams are:

- Partial exam content is pre-released to students prior to the exam to deepen the cognitive work that students perform during the actual assessment. This allows students to read information about the nature of their tasks beforehand as well as to engage with conceptually complex content that might take more time to comprehend than is typically available for an exam (Margot & Kettler, 2019). Traditionally, exam content is often encountered all at once in the context of the exam, and this rapid transmission of large amounts of information constrains the asking of interesting and higher-order cognitive exam questions due to the high cognitive load (J. Momsen et al., 2013; J. L. Momsen et al., 2010; Sweller, 2010). Throughout the manuscript, we use the term “Deepening Thought” to refer to this first element of public exams.
- The pre-release of exam content allows for communal improvement of the language used on exams. By allowing students an opportunity to give feedback on exam formats and wording through an optional editing activity, we leverage a larger group of motivated student-editors to find and fix grammatically problematic pieces of the exam. Students readily report these problems prior to the summative assessment, which is helpful to faculty. These developing student-experts can also contribute to the writing of the exam itself when faculty use public exams by being assigned editing or content creation. This is especially important because language barriers around exam content are hard to disassociate from actual struggles with content (Charity Hudley & Mallinson, 2017; Heath, 2000; B. Wiggins et al., 2020). Traditional exams are revealed late in the learning process and cannot be co-created, so the onus of grammatical perfection falls on faculty who are rarely as linguistically diverse as their student population (Brame, 2019; National Academies of Sciences, 2018). Throughout the manuscript, we use the term “Language Barriers” to refer to this second element of public exams.
- Pre-released exam content allows instructors to amplify the importance of core concepts in the course (Council, 2007). Instead of indirectly indicating core concepts through study guides or practice exams or review sessions, the concepts and skills that are core to the discipline are outlined in the context of the exam. We use the term ‘core concepts’ here to broadly describe the content that instructors believe is more central to the practice of their discipline as in (McFarland & Michael, 2020). Traditional exams do this after the fact, at which point the opportunity to direct optimal study practice is generally lost. Throughout the manuscript, we use the term “Directing to Core Concepts” to refer to this third element of public exams.

Because students and classrooms differ so greatly, the use of this public exam style is not intended to be narrowly prescriptive. Instead, we offer this stylistic definition of public exams to help guide instructors incrementally closer to more engaging assessments and b) provide a basis for exploratory research to identify impacts on and benefits for postsecondary students. A timeline comparison of a public exam and a traditional exam is shown in Figure 1.

**Fig 1.** Comparative timeline of traditional and public exams. Tasks to be completed are separated into those that are transparent to students and those that must necessarily be kept secret from students at the risk of giving away exam answers. For readers unfamiliar with traditional exams, the top timeline is offered as an approximation. The bottom timeline approximates a public exam structure. The purpose of this figure is to illustrate the differences in increased transparency and opportunities to study from exam material in public exams.

As a simplified example, imagine an exam question in which the student is directed “For ten points, explain in three sentences or less how detoxification of human blood is performed by the cells in the liver.” By pre-releasing the exam question for students but withholding only the word ‘liver’, the possible variants of the exam question are increased to include at least several organs. While providing the meta-information for the task as well as the framing of the topic area itself, this question maintains enough surprise to deeply examine student understanding. A further variant of a pre-released exam question might be: “For ten points, explain in three sentences or less how [withheld] of human blood is performed by the cells in the [withheld].” By withholding just a single additional word, students are now given direct information about both the method/scope of written assessment as well as tangible evidence that their understanding of processes affecting human blood will be crucial for demonstrating deep understanding of the topic. Other examples of public exam questions are presented in Figure 2.

**Fig 2.** Examples of public-style exam questions. For each of 4 exam questions, the pre-released version provided to students well before the exam is shown. In dashed insets are the changes made to the question for the actual version that students complete for course points. The purpose of this figure is to give examples of a few of the types of exam questions that can be used in public exams.

#### Research Framework and Questions

Our goal in this work is to demonstrate if and how public exams may affect the experiences of college students. Our methodology follows a design-based tradition in which education interventions are implemented and researched dynamically and iteratively in parallel (Collins et al., 2004). Design research is an appropriate choice because we are studying student-centered systems in context (Sandoval, 2004). To explore our research questions, we apply mixed quantitative and qualitative methods and attend to signals in the data that triangulate similarly across multiple types of investigation (Denzin, 2012).

Our research questions are the following:

- In what ways do public exams affect the learning experience?
  o Are the effects of public exams negative or positive for student learning?
  o Are these experiences affected by Language Barriers, Directing to Core Concepts, Deepening Thought, and/or Authentic Involvement?
- Do public exams exacerbate grade inequities across demographic groups?
- Do students dislike public exams?
- Can public exams be applied across postsecondary education contexts?

## Methods

### Research environments

Research was conducted at both a research university (R1) and a community college (CC) in the Pacific Northwest of the United States. Students were enrolled in lower-division courses in Biology departments during Quarter 2 of 2021. The R1 course was taught for 300 students and the CC course was taught for 48 students of which 292 and 32 participants, respectively, were included through IRB-approved consent processes (under protocol #s STUDY00012237, ECIRB-20210512 and IRB-2020-0813). These courses were chosen for consistency of general topic and level, for the large population in the R1 course which allowed quantitative analysis of subgroups, and for institutional access to research. Students in the R1/CC courses were 63%/59% non-white, 77%/66% registrar-identified female, 24%/20% first-generation attending college, 12%/24% international and (at the R1) 31% identified as being from historically underserved populations by the R1 university. Students in both courses typically have interest in a wide range of career goals around healthcare, science, research and business. Public exam techniques were used in both courses. Both the CC and R1 courses used public exams for the first time in those environments. In the large R1 course, students were graded primarily on the basis of 5 total exams given every 2 weeks throughout the 10-week quarter. In the smaller CC course, students completed a total of two exams written in the public exam style.

### Research flow

We conducted research using a design-based research methodology, which allows for preliminary research findings to be used to guide the collection and analysis of subsequent data in an iterative fashion (Collins et al., 2004). Examining human experiences in this methodology is intended to be more rigorous than simple, self-reported data, while allowing a greater breadth of possible findings than quantitative experiments alone could observe. This methodology is a good fit for education systems where iterative redesign and incremental improvement of human experiences are the primary goals of research and implementation work (Sandoval, 2004).

Here, we used *qualitative interviews* to broadly assess the experiences of students taking public exams around our three main research questions. Interviews were also used to assess differences in the learning experience between institutions. These interview findings refined our analysis of a larger data set by *coding open-ended survey items*. In parallel to this qualitative and mixed-method work, students in the R1 course took exams that used both public and traditional questions to experimentally observe signals of inequity in exam outcomes. This *within-exam experimentation* controls for student identity, instructor ability, classroom environment, and content material in comparing data from public and traditional assessment questions. This *quantitative data collection and analysis* is intended to cast a wide net for possible negative impacts of public exams on learning experiences. Any positive impacts of the public exam system that are observed are likely to be conservative because of issues with first-time implementation fidelity (in both R1 and CC courses) and incomplete application of the public exam system (in the R1 course). *Student self-reported preferences* for exam style were collected to triangulate with other types of data; these self-reported data may help to illuminate the presence of unknown negative impacts of the intervention, but is not in itself convincing of positive impacts of the intervention. The overall process of data collection is described below in Figure 3. The purpose of collecting a wide range of types of data was to broadly investigate the possible outcomes from this intervention and better understand the possible avenues for future, deeper research investigations. Here we present the results of this initial design-based research study.

**Fig 3.** Data collection scheme. The purpose of this figure is to identify when and in which class environment the data were being collected.

### Qualitative interviews

We conducted group and individual interviews to collect student learning experiences using grounded ethnographic principles (Glaser & Strauss, 1968; Rubin, 2012) and subject-centered and -driven methodology from dialectical behavioral therapy (Linehan, 2018). Eleven interviews totaled 488 minutes of recorded discourse with 20 participants. Student participants were recruited via random course list emails. Data around the interviews at both sites as well as all transcripts are available in <u>Supplement</u> 1.

During qualitative interviews, we used open-ended experiential questions to elicit a broad spectrum of conversations around students’ experiences (Cameron, 2005). Rather than providing specific questions or prompts, the facilitator followed up with probing questions about student-raised topics pertaining to our research questions (Rubin, 2012). Thematic representation in anonymized transcripts saturated (Saunders et al., 2018) the R1 site after six interviews.

We coded the transcripts of interviews on four prevalent themes drawn from our original research questions: 1) Language Barriers, 2) Directing to Core Concepts, 3) Deepening Thought, and 4) Authentic Involvement. We decided to investigate these four themes due to anecdotal discussions with many prior students. The new codes 5) Anxiety or Confidence and 6) Collaboration emerged during iterative qualitative analysis of interviews with students at the CC and R1 institutions, where students strongly expressed the importance of these themes. Lastly, the code 7) “Not Related to the Public Exam System” was designed to capture student experiences that were not part of the public exam system. The descriptive language found in the coding tables was iteratively improved for clarity and to better match student language. Transcripts were subsequently re-coded using this improved set of seven codes. The final consensus coding table of interviews with exemplary quotes is available in <u>Supplement 2</u>.

### Coding of open-ended survey items

We used open-ended survey items as a quantifiable source of qualitative data at scale. In a participation-only study, all students at both sites were asked to answer the question: *“Did the style of exams in [this course] work for you? Why or Why not?”*. Cognitive testing for validity of this question was performed with a separate group of students that were of the same age and progression as students at the CC and R1 sites. Responses to the final version of this open-ended survey question were collected and anonymized from 242 participants at the R1 site and 32 participants at the CC site.

Within the open-ended survey responses, the seven final codes were iteratively coded, discussed and then re-coded for presence. Each code was also sub-coded as positive or negative with regards to literature-based learning outcomes for students. This was not opinion-based coding on the part of students, but rather researcher-based assessment of whether the practices or experiences presented were positive or negative based on nationally recognized educational best-practices texts including How People Learn II (National Academies of Sciences, 2018) and the biology-focused AAAS document Vision and Change (AAAS, 2011a). In other words, these results were not coded for what students enjoyed or appreciated (see examples in-text below) but rather for conditions in which learning is likely to be supported. Two members of our research team independently coded 15% of the blinded responses and achieved an acceptable interrater reliability score of kappa = 0.88 (McHugh, 2012). One researcher coded the remaining blinded responses.

For an examples of the positive or negative coding, a student who indicated “*The public exam made it harder to know what I needed to know*” would be coded into the category of ‘Directing to Core Concepts’ and as a ‘Negative’ impact, since confusion about core concepts is a problematic distractor for learning across fields (Meyer, 2004; National Academies of Sciences, 2018; R. Keith. Sawyer, 2005). In another example, if a student indicated that they “*hate public exams because they force me to think more deeply*,” then this would be coded as a ‘Positive’ impact within the theme of ‘Deepening Thought’, even though the student may not have enjoyed that aspect of the learning challenge. Additional examples and final codes are available in Supplement 3.

To determine if the prevalence of any codes was significantly important for learning experiences, we calculated the percent of students who provided feedback on each qualitative theme of the public exam system in the open-ended survey items and whether that feedback was positive or negative. To determine if there was a relationship between the type of feedback students provided (i.e., about the public exam system or not) and the nature of that feedback (i.e., positive or negative), we conducted a series of Pearson’s chi-square tests of independence for each of the six codes of the public exam system as well as an aggregate of all six codes. A control for statistical tests was needed that is more conservative than a simple 1:1 null ratio. We used Code 7 (‘Not related to the Public Exam System’) as a control group, which is more conservative than a simple control ratio like 1:1 and thus controls for the general tendency for participants to report positive experiences more often than negative experiences. When a given count in the contingency table was too small (i.e., less than five) to conduct a chi-square test, we used a Fisher’s exact test (Bower, 2003; McCrum-Gardner, 2008).

### Within-exam experimentation

Within the large RI course, students completed five summative exams that contained a mix of traditional ‘surprise’ and ‘public’ style assessment questions. For this course, all exam questions were written in multiple choice format. The relative amounts of traditional or public exam questions changed throughout the course. Students began the quarter with two exams that used the same distribution of multiple-choice exam questions: 15 public-style exam questions and 10 traditional, surprise-style exam questions. Subsequent exams (in response to student survey responses, see discussion) included 20 public-style exam questions and five traditional, surprise-style exam questions. The purpose of this within-exam experimentation was to collect well-controlled data that might lead to observation of inequity with this style of assessment, rather than to demonstrate value of an assessment style. Because the variation between assessment styles happens within each exam, this experimental design controls for scientific topic areas, for student identities, and for the instructor, among other variables that are otherwise difficult to control.

### Quantitative data collection and analysis

Within the large R1 course, the following discrete data were collected for each participant: College GPA, course grade, exam results for each question on each exam, scores for participation-based assignments, completion or not of an exam editing activity, and (via the university registrar) race/ethnicity, gender, international student status, first-generation in college status, and inclusion in the university-assigned Education Opportunity Program (EOP). This last categorization is particularly important to this work: The R1 institution defines “under-advantaged” students as students identified as part of the EOP and these students hail from educationally or economically disadvantaged backgrounds. Because this EOP categorization is based on family income and other variables not typically represented in simpler demographic statistics, we followed Freeman, et al (2017) and Wright, et al., (2016) and selected this as the single variable to pre-build models for analysis. All data collected using these methods are available in anonymized form in <u>Supplement 4</u>.

In order to determine if students performed differently on public or traditional exams, we used a two-sample t-test to compare the total percentage of points students earned on all public exam questions and all traditional exam questions throughout the term.

In order to determine whether there were differences in exam performance on each type of exam question based on students’ demographic characteristics, we used linear regression models and included gender (male/female), EOP group of interest (yes/no), and overall GPA (from the registrar on a 4-point scale) as predictors (Example model: percent score on public exam questions ∼ gender + interest group + GPA). Gender has been shown to affect student exam performance (Odom et al., 2021) and students in our EOP group of interest have been found to do worse than their peers on exams at this institution (Cooper et al., 2020). We acknowledge that registrar data for gender that includes only male/female do not best represent all individuals’ gender identity and that not every person identifies in the gender binary (Cooper et al., 2020), but we did not ask students to self-report their gender.

To examine potential demographic differences in students’ self-reported preferences for the proportion of each question type on an exam, after the second and third exams, we asked students if they would prefer to have more public questions, fewer public questions, or keep the same ratio of public to traditional questions for future exams. After the fourth exam, we asked students if they would prefer more or fewer public questions with no neutral option. We calculated the percentage of students who selected each option and assessed potential demographic differences of students’ preferences after the second and third exams using multinomial regressions and using logistic regression for preferences after the fourth exam. We again included gender (male/female), EOP group of interest (yes/no), and overall GPA (based on registrar data on a 4-point scale) in our models. [Model for post-exam two and three preferences: exam preference (more public/fewer public/same) ∼ gender + interest group + GPA; model for post-exam four preferences: exam preference (more public/fewer public) ∼ gender + interest group + GPA.)]

Preceding each exam, students were given the opportunity to provide edits on the public portion of the exam. This was an optional part of a required online assignment which students were able to bypass and still receive full participation points. To investigate the extent to which the experience of providing edits on the exams might correlate with their overall course grade, we used a linear regression with the total number of exams for which the student provided edits, EOP group of interest (yes/no), and overall GPA as the predictors in our model. [Model: course grade ∼ total edits + interest group + GPA.]

## Results

### Qualitative interviews

Interview-based methods were used to guide the overall flow of research, and also to better understand whether public exams might be applicable to community college courses, which are generally smaller and less available to quantitative research, therefore, we undertook qualitative interviews in a closely-matched community college course. This CC course closely matched the R1 course in terms of topic, location, timeline, and the first-time use of the public exam style for the course. Our primary codes were evident in similar proportions, and student comments to interviewers brought up similar challenges and gains. No thematic signals appeared in one environment and not the other.

### Coding of open-ended responses

Students in the large R1 course answered a survey item: “*Did the style of exams in [the R1 course] work for you? Why or Why not?”*. All coding data for open-ended survey items is available in <u>Supplement 5</u>. When compared with a conservative control group using Code 7 (‘Not part of the public exam system’), we observed a strongly significant statistical signal for the overall positive impacts of public exams (Table 2, Row 1). No significance (positive or negative) was observed for student mentions of language barriers, authentic involvement in the process of assessment, or collaboration. Learning experiences with ‘Directing to Core Concepts’ were significantly positive (p value = 0.0002). Learning experiences with ‘Deepening Thought’ and ‘Anxiety or Confidence’ were also significantly positive (p values = 0.004 and 0.0101, respectively). Positive or negative experiential effects showed no statistical difference for students in the EOP group. These data suggest that students’ unprompted experiences with public exams are predominantly positive, which correlates well with preference data described below. These data also triangulate well with interview results noting that deeper cognitive work, decreased anxiety, and more efficient directing to core concepts are likely outcomes of public exams. These results of the quantitative analysis of open-ended coding are presented in Table 2.

### Within-exam experimentation

Comparisons of exam outcomes on mixed traditional/public exams were used to quantitatively assess possible issues of equity. Student exam outcomes on public and traditional exam questions were analyzed for two groups of students: A university-identified diverse group of students in the Educational Opportunity Project (EOP), and the rest of the student population. As shown in Figure 4, in our model, we observed that all students performed better on public exam questions compared to traditional exam questions (blue lines). Because of the differences in learning processes between public and traditional exam questions, this difference in performance is not evidence of learning differences between content assessed in a given method. We also observed the expected decrease in high-stakes exam scores across question types for students from EOP underrepresented groups (red lines). The combination of these trends was consistent for students in both EOP and non-EOP groups, giving no indication that public exam questions resulted in increasing inequity.

**Fig 4.** Exam outcomes for traditional- and public-style exams. Color plots are separated by underserved Education Opportunity Project (EOP) group in yellow and non-EOP (majority) group in purple. The EOP group is a university-defined amalgamation of underserved identities that was chosen as a single, objective variable to collectively indicate students for whom traditional postsecondary education outcomes has served poorly. Significant differences were found in the higher scores for students on public style exam questions as compared to traditional exam questions (indicated with blue asterisks), although the difficulty or achievement on these exam questions cannot be directly compared as the learning structures were different. Significant differences were found in exam scores between groups of students, which is consistent with pernicious gaps in outcomes in postsecondary education (indicated with red asterisks). No differences in the patterns of outcomes for traditional/public exam questions were found in either group of students, which is consistent with public exams being similarly equitable compared to traditional exams. The purpose of this figure is to display the outcomes of this experiment intended to observe any differences in equitable treatment of students if they exist.

### Student self-reported preferences

In the large R1 course, students were asked about their preferences for public or traditional exam questions. After completing two mixed exams with 15 public and 10 traditional questions each, 41% of students preferred to keep the same distribution for future exams, 3% of students wanted more traditional questions, and 56% of students wanted future exams to have a greater proportion of public-style questions. After listening to this student voice and increasing the proportion of public questions for the following exam, students were surveyed using the same questions. After this exam made of 20 public and five traditional questions, 67% of students wanted to keep the increased 20:5 distribution while 6% wanted more traditional questions and 24% wanted more than 20 of the 25 questions to be public. Course instructors kept the 20:5 ratio for the next exam, and students after this exam were given only two options in order to better understand the preferences of the majority of students. On the final survey, prior to the final exam, 15% of students wanted to decrease the number of public questions and 85% wanted to increase it. Throughout these exams and the overall self-reported desire for more public exam questions than traditional questions, there was no significant signal for a demographic basis on which these preferences were made, nor was preference correlated with course grade outcomes.

#### Does editing of the exam affect students?

As part of the public exam, students were given the opportunity to suggest edits or contributions to the public exam document itself. Edits suggested by students included highlighting grammatical errors in the exam, suggesting improvements to the content text in the exam, and suggesting creative additions to partial questions. Students who undertook these optional, non-credit opportunities, after controlling for course grades and demographic backgrounds, were significantly more likely to improve their overall course grade (p value = 0.000402).

## Discussion

Here we discuss the results in light of our research questions, as well as future research questions and limitations of this work.

### In what ways do public exams affect the learning experience?

Analysis of students’ open-ended survey responses showed an overall significant and positive impact of public exams on learning experiences in a large STEM course. The positive impact of public exams on learning experiences was significant even when controlled against other student responses in the same environment. They also triangulate well with themes from interviews, self-reported preference surveys, and with anecdotal narratives from public exam practitioners more widely.

‘Directing students to core concepts’ was significant and positive, which speaks directly to a consistent challenge for novice learners. While accepting the deluge of information present in any fast-paced course, novice learners struggle to develop mental models to organize incoming information (Meyer, 2004). Modern courses typically offer an array of learning materials to assist students in developing understanding of which pieces of information are core to the discipline and which pieces of information are facts or ideas that simply reinforce the concepts that an instructor feels are core to mastering the material in their course. Within the public exam structure, students have early access to exam materials that are directly connected to course points. Instead of deducing core concepts from lectures, assignments, study guides and other indirect sources, students in a public exam course have the contextual clues to infer value by placement (or not) on the actual assessment itself. In contrast, traditional exams typically hide these valuable assessments until the moment of the exam itself. The significant, positive impact of ‘Directing to Core Concepts’ on public exams may be a reflection of these benefits to learning. As one R1 participant said:

> *“…they provide me with some direction on what to study a lot for. I think that there’s a lot of material that’s covered in this course throughout the lectures, and it would be hard to remember every single detail from the textbook, so I think the guidance of the public questions really helps you to look back at that specific part in your notes and/or the lecture to refresh your memory on what you learned.”*

Many instructors are frequently confronted with, “What do I need to know for the exam?” from students. Perhaps similar to some types of practice exams given before an exam (Crowther et al., 2020), public exams help students to focus on key content instead of feeling overwhelmed.

‘Deepening Thought’ was also found to be statistically significant and positively aligned with student learning experiences. The logistical constraints of exams can push instructors to a ‘flattening’ of thought; this problematic trend contributes to science being aligned with low-level accumulation of simple facts (Crowe et al., 2008). It is possible that benefits from public exams come from the increase in higher-order exam question (Anderson & Krathwohl, 2001; Barnett & Francis, 2012; Lemons & Lemons, 2013), which was the intent of the designers but not assessed in this study. The significant, positive benefits from the public exam style may be due to shifting exam-provoked thought from a one-time performance into a longer and more collegial set of learning cycles (Schwartz et al., 1999). Because students are less limited by the time needed to read and comprehend a complex exam scenario, more interesting scenarios can be approached by the instructor. Assessment materials transmit the values of the instructor into real terms (G. Wiggins, 1998, 2011). Students reported being challenged by the public exam format to more in-depth learning of a concept. In interviews, students realized that with the extra time to think about and discuss exam questions, there was an expectation of exam responses that demonstrated deeper thought and synthesis. For example, a CC student said:

> *“Personally, I liked this type of exam a lot more. I didn’t feel like I had to memorize anything. More like I understood the concept and could be asked questions about [it] from multiple angles. It helped learning with others as well because when explaining to other people a certain topic, and they begin to understand tells me that I understand the concept exceptionally well.”*

As more disciplines make calls for deeper critical thinking skills (AAAS, 2011b; Halpern, 2001; McConnell et al., 2019; Engage to Excel: Producing One Million Additional College Graduates with Degrees in Science, Technology, Engineering, and Mathematics, 2012), it is possible the pre-release of exam material (as in Crowther et al., 2020) is a motivating factor in pushing students to do, share, and enjoy this deeper thought.

‘Anxiety and Confidence’ around education (and more specifically exams) is a constant and increasingly-pressing concern (Disability, 2017; Health, 2020). While this is well-studied in STEM courses (Cooper et al., 2018; Downing et al., 2020; Schussler et al., 2021), it may be more relevant instead to courses for which high-stakes exams are a primary feature (Brady et al., 2018; Culler & Holahan, 1980; Harris et al., 2019); STEM courses generally meet this description (J. L. Momsen et al., 2010). Learning is maximized at moderate levels of stress (Rudland et al., 2020), but greater stress hampers learning and motivation and disproportionately affects students from groups traditionally underrepresented among those with college degrees (Lee et al., 2021; Medina, 2011; Misra & McKean, 2000; Vaidya & Mulgaonkar, 2007). There is some indication that this current most-diverse and most-economically challenged generation of students in college are also understandably the most over-stressed that have ever enrolled (Lederer & Hoban, 2020). With less anxiety associated with the surprise of the exam, they were able to feel more confident and prepared. An R1 student noted:

> *“… with the availability of the public exam I am able to study the possible directions the questions might take. It reduces the amount of stress and anxiety I usually get when I take exams, I feel more prepared.”*

Public exams may help students to alleviate some of their stress through some familiarity with non-content information. Strategic points like where to focus effort and time can be usefully discussed and digested at home. Shifting non-content mental effort away from the exam performance time may explain why coding analysis shows better outcomes in public exams, which aligns with prior research (Hacker et al., 2008; Pate et al., 2019). It is also possible that the steps made towards exam transparency have a role to play, as signals of equitable behavior on the part of powerful authorities may suggest to students that they need not worry about being caught in a negative power-dynamic over other disputed elements within assessment (Bang & Medin, 2010; Bell et al., 2012; Fredricks et al., 2004).

Public exams include opportunities for students to authentically engage in the creation of the assessment by editing and making suggestions. Students who took advantage of these opportunities also performed better in the class. Those edits are sparse among many exam questions, and the changes suggested rarely alter content, so this trend is unlikely to be explainable by gains on the specific question edited by the student. The statistical model used controlled for demographics and for student course grade, so it is less likely that this is a self-selection of which students choose to take on this extra task. If the correlation observed (p value = 0.000402) indicates a causative relationship, then it may be explainable in one of three ways. It might be that students who interact with the exam in this editorial mode are finding a new way to engage with the material. By seeing the content from a different angle, one more closely aligned with the perspective of the faculty instructor, they may find their own perspective on the content to be broadened in useful ways. This is in line with learning theory about critical thinking skills (Halpern, 2001). A second possibility is that engaging with assessment as a partner, even in a temporary way, may help students to feel authentically involved in the process of assessment. Affective experiences can improve learning (Dweck, 1986), so this specific observation would be in line with learning theory. Lastly, it is possible that this result conflates students who did not provide edits with students who never accessed the public exam materials (even after frequent instructor guidance), which might contribute to their lower course grade. In the first two models, the benefit to student learning would be valuable and further research will be required to better understand how, for which students, and under what conditions this benefit is generated.

### Do public exams exacerbate grade inequities across demographic groups?

Prerequisite to understanding more about the specific impacts of public exams, and as part of feminist and anti-racist drives within education research, we want to ensure that public exams do not contribute to the extant inequities in student outcomes within postsecondary education (Museus et al., 2015). Those concerns are most pressing for assessments, which often represent a gateway for student success at which inequities are both created and revealed. The primary goal of our quantitative within-exam experimental design in a large R1 course was to help understand if public exams are creating or exacerbating inequities for students from groups historically marginalized in postsecondary education. Analysis of question-by-question exam outcomes in a large course is our most likely opportunity to observe a signal of inequitable outcomes. Close analysis of question-by-question outcomes make clear that these pernicious gaps in outcomes exist beyond our research environment: Students from underrepresented groups are associated with lower scores on both public and traditional exam questions. Clearly, improving outcomes for all students will take much more than the use of public exams. Of particular importance for our study is that outcome gaps are not exacerbated by public exams. In other words, the gaps between public and traditional question outcomes are not different between groups of students or exacerbated by this new exam style.

### Do students dislike public exams?

Beyond these three emergent aspects of the learning experience, student self-reported preferences for exam style were strongly in favor of questions in the public exam style. These positive preferences are not explainable by demographics or class success. It is not clear why students prefer public exams from this study; it may be a preference for deeper thought, for more directed practice, for the novelty of a new challenge, for their involvement in the assessment process, or for the perception that pre-released exam information will make for an easier course assessment (although the similar scoring averages between exam question types help students to understand quickly that this is not the case). Further research is needed to better understand the parts of the public exam system that are preferred and how to maximize these elements usefully. If these preference survey results can be taken at face value, then student preferences for public exam questions are strong, are in accordance with findings from open-ended coding and qualitative interview analysis and are unlikely to indicate any widespread dislike of the public exam style that might forewarn practitioners from attempting to implement it. From the lens of a classroom instructor, it is also encouraging to see a strong signal of preference for a style of exam that requires more complex thinking.

### Are public exams likely to be applicable across postsecondary contexts?

The results of qualitative research across two disparate types of institutions give early evidence that public exams can fit in an array of environments. Analysis through iterative coding of interview transcripts brought us to the conclusion that students in the two courses had similar experiences with public exams. This is an initial attempt to explore the possible broad application of public exams, and clearly more research will be required on a greater scale to make similar conclusions. In the meantime, the outcomes of these analyses are consistent with public exams being similarly applicable across these two institution types.

For example, a CC student noted that:

> *“We were able to sit down and start bouncing information off of each other and asking different questions about the questions…just kinda sharing information right before the exam and that just gave me so much confidence as to how much I know going into the exam…”*

This student suggests a deeper questioning style beyond memorization and notes the effect of deeper thought as well. A second CC participant mentioned:

> *“It helps more with like understanding but sometimes when you’re panicking about an exam you’re like ‘I don’t want understanding; I just wanna know’ but at the same time you do have to understand things…if we hadn’t had the public exam I would have studied all five of the chapters and had like less knowledge on each of the things and I don’t feel like I would have remembered the exact definition of phenotypic plasticity as well as like when I saw the question and was like, I really do need to know this for the exam.”*

These three coded themes of Anxiety and Confidence, Directing to Core Concepts and Deepening Thought are evident here and were strongly present in both environments. Weaker themes of Collaboration, Language Barriers, and Authentic Involvement were evident in both environments but to a consistently lesser degree. While we did identify emergent themes in this work, no thematic signals appeared to us in one institutional environment and not the other.

#### Redesigning assessments through a lens of educative assessment

From the beginning, the development of the public exam style has prioritized educative improvement over other outcomes and purposes of summative assessment (G. Wiggins, 1998). The public exam style is educative in that it both informs and improves outcomes. Our initial research here indicates that student behavior is informed by the different challenges to be more focused on the core concepts of the material. Outcomes are improved, as best we can observe through independent coding of learning experiences, to facilitate deeper thought around conceptual science material as well as doing so with less perceived anxiety around the assessment process. Especially in light of continuing challenges in STEM fields to retain students and to push those students beyond simple memorization, the educative capacity of our choices around assessment style may become increasingly important.

#### Limitations of this study

As an initial foray into research on public exams, this study has many limitations. The design-based research model used in this study is likely to unearth important features of the student educational experience. However, this model is not intended to prove that a particular feature is more or less important than another, or to compare overall impacts of the learning experience on learning or career success. Future research, using longitudinal analysis and topical challenges, will be important for assessing the overall impacts of the public exam intervention beyond these initial analyses. Constructs like anxiety are treated as emergent themes; future research should apply established theoretical frameworks around anxiety to make use of established survey instruments that may be a good fit to better understand the ways and extent to which public exams affect student anxiety. The core features of public exams are examined as a unit, and more work will be required to understand if benefits can be achieved modularly. Largely a single-course study, this analysis may be conflated by the specific instructors or the environment of early 2021 (in itself, a unique time to be working in postsecondary education during a pandemic). Education impacts tend to be relatively small in comparison to the effect sizes in other fields, so it is possible that other important features have gone unexamined for lack of analytic power in a single course of 300 students. This is especially true for particular groups of students of historic importance, for whom numbers are smaller and backgrounds unique to this particular study environment. The lack of evidence of specific equity gaps created by this exam style is not proof of its absence, as this research may not have had the experimental power to observe subtler demographic shifts. Furthermore, the novelty of the public exam style in post-secondary classrooms means that specific instruments for investigating assessments on more traditional models were not available. Future research should involve validation of specific instruments for assessing these learning cycles, such as those seen in (Arikan et al., 2022; Chang et al., 2021; Hicks et al., 2017; Johnson et al., 2022; Reynders et al., 2020). Perhaps most importantly, this study did not directly assess student learning but rather the student learning experience. We hope that the benefits demonstrated, combined with positive anecdotal reports on the strengthened student/instructor relationships in similar courses, motivate future research to better understand how varied assessment styles can better serve the next generation of students and improve on this work.

One salient criticism of public exams is that the process can be summarily characterized as ‘teaching to the test’. This pejorative has a long and well-deserved history in K-12 education, especially in situations where externally-created assessments are linked to a motivation to maximize scores for the purposes of accumulating outcome-linked resources (Jensen et al., 2014; Ravitch, 2020). We propose that many college and university exams are fundamentally different in that the instructors have wide purview to create exactly the kinds of assessments that reflect the values, skills and content needed in modern pursuits. In other words, professors can create the kinds of exams for which ‘teaching to the exam’ is inherently valuable for students (Jensen et al., 2014; G. Wiggins, 1998). We hope that public exams are a useful way to do this.

#### Considerations for interested practitioners

Transitioning from traditional exams to a public exam style is a low-tech strategy to employ several best practices likely to improve student learning. Instructors found that they could make simple changes to the exams or exam blueprints that they were already using by withholding some of the information. In many cases these adjustments shorten the exam by augmenting the higher cognitive exam questions and allowing students to discuss core concepts in more detail because students had more time to reflect on the question. Additionally, instructors were receiving meaningful feedback from students during the editing process of their new public exam that improved the exam questions. Importantly, instructors do not need to adjust the entire exam to the public method. Instructors can slowly transition to a greater percentage of the exam being publicly available over a quarter, semester or academic year. Anecdotally, students were excited to be part of the public exam process and participate in a new assessment strategy.

Postsecondary instructors have numerous choices when designing exams (Gezer-Templeton et al., 2017; Hodges, 2004; Knierim et al., 2015; Wieman et al., 2014). For those who want to take up public exams as a style of assessment, we suggest adjusting a small number of questions on an upcoming exam into a public-like format as depicted in Figure 2. This helps create a positive feedback loop for instructor design and feedback from students, and it also helps avoid taking on an unsustainable overhaul of all assessment in one course. In our experience, instructors who take up a few challenging pre-released exam questions a) quickly develop the communication needed for students to understand how and why to access the materials, and b) invariably lead to greater use of these methods in future assessments. An earlier, deeper, non-peer-reviewed logistical discussion of public exams within the field of molecular biology may be of interest to practitioners (B. L. Wiggins, 2019).

We have proposed that public exams may be a strategy to address some of the anxiety associated with taking exams. It is important to note that the adjustment period as instructors implement a new exam style may be longer for some students compared with others. Some strategies that could facilitate a smoother transition are to start off with lower stakes quizzes or exams, practice assignments or quizzes, or set up student groups where students can support each other. Although we did not find support for “Collaboration” in the quantitative coding analysis, at least some students recognized the advantages in collaborating when preparing for the exam. A R1 student described this by saying:

> *“I have noticed that it only works for me when I work with other people in study sessions. I try to study on my own. I have a more difficult time understanding the material, which is something quite new to me since I am used to studying on my own. But overall, I like it.”*

Students may not have recognized that collaboration was not only acceptable but highly encouraged, often not utilizing that strategy until later exams. As a CC participant explained:

> *“I loved the second exam because I was able to meet up with others outside of the classroom to go over a couple different concepts before the exam.“*

Anecdotally, instructors using public exams routinely describe increased collaborative studying similar to what these students described. Emphasizing and encouraging collaboration as a strategy for student success on the exam may be another way the instructor can facilitate the transition from a more traditional exam model.

## Conclusion

We analyzed the impacts of public exams in STEM courses. Our mixed-methods design research shows that students find significant positive impacts on their experiences. Those impacts are largely focused on improving the direction of students to core concepts, the deepening of thought in the assessment process, and helping students to manage anxiety. The public exam method is likely to be similarly equitable to traditional methods and potentially applicable across institutional contexts without exacerbating issues of educational equity. We present this work in the spirit of improving assessment for all students as a core feature of critical, high-quality education.

## Supporting information

Public Exams Supplement 1 Interview Transcripts

Public Exams Supplement 2 Interview Coding

Public Exams Supplement 3 Survey Item Coding

Public Exams Supplement 4 Data Set

Public Exams Supplement 5 Coding for each

## Acknowledgments

This work was funded by the generosity of a private, independent research gift from CourseHero, without which none of this research would have been possible. Participants and their data are all protected by Institutional Review Boards under #s STUDY00012237 (University of Washington), ECIRB-20210512 (Edmonds Community College) and IRB-2020-0813 (Oregon State University). We appreciate the efforts and contributions of Greg Crowther, Deb Donovan, Kelly Hennessey, Lori Kayes, Teddy Maley, Devon Quick, Christine Savolainen, Mandy Schivell, Katie Simons, Shelley Stromholt, Jeannette Takashima, Liz Warfield, and Seth Wiggins. C.A.B was supported by an NSF Graduate Research Fellowship Program Grant No. 026257-001 (any opinions, findings, and conclusions or recommendations expressed in this material are those of the authors(s) and do not necessarily reflect the views of the National Science Foundation). Thank you to the volunteers and staff at Seattle Public Libraries for maintaining access to research-quality facilities even throughout a global pandemic.

