## Supplementary material for "Public exams decrease anxiety and facilitate deeper conceptual thinking": Public Exams Supplement 1 Interview Transcripts

| **Interview #:** | **Site:** | **# of student participants:** | **Duration (minutes):** | **Starts on Page #:** |
|---|---|---|---|---|
| 1 | R1 | 2 | 47 | 2 |
| 2 | R1 | 1 | 26 | 11 |
| 3 | R1 | 1 | 41 | 17 |
| 4 | R1 | 2 | 39 | 25 |
| 5 | R1 | 2 | 58 | 33 |
| 6 | R1 | 1 | 55 | 43 |
| 7 | CC | 2 | 55 | 53 |
| 8 | CC | 4 | 49 | 61 |
| 9 | CC | 3 | 50 | 73 |
| 10 | CC | 1 | 33 | 83 |
| 11 | CC | 1 | 35 | 92 |

### ***<u>Interview #1 April 13 2021 @R1 site (Leah Lily interviewer = bold, with students J and K)</u>***

L: Blah blah You read the consent form so you know a little bit … What I'm interested in looking at here is what strategies we can use to improve college teaching so that students like you have a better experience and so we know more about what we as instructors can do to make things accessible for all students and effective and responsible—er responsive to your time of course because students are so busy and have so much going on. And so what I do is basically I talk to a whole bunch of different students, we try to reach out to as many people as reach back to us because we try to spread a wide net and talk to a lot of different people. And then I go through these transcripts and try to look for patterns about what students say about … What they're interested in maybe having more of or less of and I try to look for big themes that we can use to inform um either more research into a specific intervention or change in courses or some conclusion about, we sometimes find things about learning itself, which is interesting for me. Any questions before we jump into it?

J: Is this geared towards online learning or remote learning or is it more like just classes structures in general?

L: Good question. Just because we're stuck doing online learning as we do this research I think it's dominating for online learning but we're interested in um some of the sort of the fundamental structures of the courses and how we assess and how we introduce that content. Does that make sense?

J: Ok, Yeah I'm good with that, are you looking for like me to give like a background of what classes I've taken at UW or STEM classes

L: What I would like to start with is what's important to you, so, because that matters, what's sticking out for students is relevant to what we should be looking at. And so If you want to give a background that informs your experience in Bio 118 then that's certainly relevant and I'd love to hear it

J: And so I'm talking about what's important to me with how the education is delivered or how, just like, how the resources in the class are able to aid me in learning?

L: Um well boy that's a really good question. Both is the quick answer. For this particular study we're interested in assessment so we're trying some different things with the exam styles, and so thinking about the exams themselves would be really helpful for me. But definitely if something came up in the course that is important, or something about the content delivery comes up for, or something that informs how the exams work for you, like Something in the teaching that made the exam work or not work for you then I'm interested in that.

J: Ok, Well this is the first STEM course I've taken at UW that actually has a more predominately multiple choice style test, all the other ones—so basically I've taken the chem intro series and I also took the Bio series at UW and all the tests that I've taken up until this point were either hybrid of multiple choice and free response or just all Free response. And so you know, The nature of the UW specifically, I've noticed that the classes they go at a really fast rate and they expect you to become really familiar with some material that could be considered dense in a relatively short amount of time and I'm not dissing that because I feel like that's most universities, especially those with a pretty prestigious research background like UW but It's always kindof bugged me a little bit how they really they ask for not just knowing the content, but kindof just like mastery of content that like you're applying, you're always applying it to like case studies, which is—I like those questions, they're interesting, but at the same time some of them can be really hard and it's like, man I don't know if I've learned this material well enough, I've been exposed to it for a long enough time for me to do my best on the exam, And then you kind of get this feedback where you'll take an exam, you'll get a poor score, you're like, is this on me, do I need to totally shift up this, was it on the test, like I'ts probably mostly on me, And so you kindof get this sense of inadequacy and then that really fuels like I dunno, if fuels self doubt and it fuels all those negative emotions that I don't feel are all that healthy, I mean it's a very common part of the college experience, no doubt, I feel like you could ask engineering students computer students, Any STEM major and then I interact mostly with STEM majors but I'm sure you could ask someone else too, and these negative feelings don't contribute to them having a really great college experience. But if I'm talking more specifically about the recent exam I've taken in BIO 118, I kindof liked the structure, it was a nice switch up to –what I appreciated about it was that it was a mix of questions that were kidof at the intermediate level and then some that were at the applied level so I felt like I could at least build up my confidence and feel secure in what I did know, So I had that foundation and then I was like ok, now there's these challenging ones, if I get a—if I don't get these right you know my whole grade isn't gone down the tank because I at least had that foundation to not just get a 50 back, you know? So I think that the test from Dr. Wiggins and Dr. Hennessey is something that I'm totally interested in their style and they seem really um invested in trying to get as many students on board with what they're trying to do and just trying to get students maybe not so focused on tests but focused on the learning aspect, and I really, I like that a lot so far. Is there something I haven't really touched on that you want me to touch on? Like that I've been talking about so far?

L: No thank you that was great, really good intro. There's some stuff that I want to go back and ask you about but I do want to hear from K. So yeah, how's Bio 180—118 treating you?

K: This is the first psychology class I've taken, I've taken other STEM courses at UW, kindof the same thing that J said, a lot of the STEM course are like hybrid, they're like free response, or there's like 10 multiple choice and then free response Where you type in the answer and then show your work, so this is definitely new for me as well. I think I do like how he structures it, how some things he shows and then some he doesn't because it really gets you thinking about I do agree with how you kindof have to master that subject in order to answer the question, whereas—but I understand why he does it because if there were just a simple like "what is a cell" then the average rate of the exam would be very much higher and I understand why he does it but It's kinda difficult to grasp in such a short amount of time but it does make sense because it's a really rigorous subject and UW is also known for that and other universities as well. But it also can be discouraging at times when you study for two weeks at a time and you think you understand a topic or certain subject and then it just like, it kinda brings you down when you think you know it and then they ask you a question that is related to the subject but you hadn't really thought about it that way, which I understand because when you're in the health care system There's gonna be questions like that, but I do like the structure of the exam and how he does it because he does explain very well that exams aren't a big reflection of who you are so he does a good job of explaining that to us and making sure that you don't feel discouraged but sometimes it's hard not to feel discouraged when you get back your exam score and it's like oh, you studied so hard and it's not a reflection of how hard you worked but did I apply that right, did I not apply that right. Did I miss anything?

L: That was great. Um, I guess so what I'm hearing from both of you is that it seems like no matter how well you end up doing on an exam, just having to struggle with those higher level questions is demoralizing whether or not you get them right? Is that about right?

K: Yeah I would say so, I would--if I do get it right it's like "oh yes thank you" because sometimes it's really just guessing for me there's like one or two that I really want to do because I can narrow it down to those two but then it's like a tiny detail that I never thought about and I'm sure he mentioned it in lecture but it's closed notes so I'm like trying to think back like what did he say in the lecture, so it's like those tiny little things That I take note of when I'm watching lecture and then the recorded things like that but it's just kindof for me personally hard to reflect back when there's two options and they're very very similar and that's why some scores are so low which I understand because they want you to understand it on a very kindof detail so you can apply it in the field. But that's just how I feel

J: I can build off what K is saying, like when you have it narrowed down to two answer choices then it really does require a pretty nuanced understanding of the topic, kindof like, it's really just critical thinking at the end of the day, like especially with physiology you know how it's all this happens then that changes this then that changes this, like if you don't know that middle step than you might miss these steps down here. I guess maybe in the past when I've taken an exam and it's mostly free response, like they--I guess that gives you the opportunity to get partial credit and I guess that is somewhat of the advantage of taking a free response exam as opposed to just multiple choice where it's A or C and you picked A and the answer was C um so In that sense some of the harder critical thinking questions might be better in kind of a free response form, it does put off a lot of students especially when they see a big table they've gotta fill in and it's just like ugh, I don't know if I know this and you get like this pre-question dread like even before you go in and start doing it But it's, if it ends up giving you partial credit where you would have either gotten a zero or a five I think that is kindof nice.

L: K, I saw you nodding with that "pre-question dread" so can you tell me a little bit on that?

K: Yes I can speak on that, I feel like a lot of college students feel that as well, especially when you're taking an online exam, It's a really different feeling than when you're taking an in person, like when I take my STEM courses in person it's really different, like it's like oh you, we studied all week or whatever like and you go in and you just take it and you're in that mindset of like oh everyone around me is taking the exact--not the exact same test but the same midterm or final or whatever, so you're kinda like ok I can do this. But when you're online it's kinda hard, you don't see everyone around you, I feel that like pretest like Not anxiety but like I'm just dreading it and some of them are like, not long, but I think one of the case studies was like you have to read a little bit so you have to hurry up, hurry up. Cuz he does have the 2hr time frame which is really nice so you get to think about it a little bit longer and um a lot of my other STEM courses that require free response I like how they're partial because like J said if it's C and you choose A it doesn't really reflect your understanding but I understand if it is free response it can be intimidating to oh like write as much as they know, and some people will write as much as they know but it might drag on and that kinda might show that they don't know what they're talking about they're just kinda writing to see if they can get all the points So that's also where it's kindof hard to decipher whether to go free response or multiple choice during an online exam.

L: Mmm.  Sounds like the communal aspect of taking an exam in person is resonating with you

J: I mean surely seeing all your peers in the same room as you struggling with the same exam even if it's at the same time, like online yes it's at the same time but you don't get that same physical feeling aspect that K was talking about because like I'm in it with everyone else At this place, at this time and yeah I like that aspect of taking classes in person, um, even though that is a whole different kind of stress at the same time, but you know, K also mentioned the time window that Ben has added and I also really appreciate that.  You know for me I'm kindof a slow test taker and especially on these kinda critical thinking questions I want to have the time to have to parse it out in my head and some of the other kinds of classes I've taken it's always just such a rush to submit the test and Canvas pops up and it's Counting down from 10, it's like noooo, I've ran out of time again, I hate that, but with this last test anyway I felt like I had time and I think I submitted like 15 minutes ahead or like 20 minutes ahead or whatever, I think I had class.  It definitely—even if you don't utilize all the time just knowing that you have it gives you kindof a sense of security and it calms down some of the anxiety that you might have going into it.

L: Good stuff, Yeah, ok.  Um yeah, so the emotional aspect of taking an exam is something we've kindof been looking at and we've been thinking about because of course its great to be able to teach the content in a better way or to prepare students for the actual questions in a better way but we also know that students get emotional around exams, and exams are an emotional thing, you know it's not a student problem, exams are stressful.  Is there something that maybe you've been thinking about like, I wish we could you know, see the questions in advance, I wish we could like have a study period something like that that would help with the emotional side of things?  It sounds like the extra time is definitely one thing that's good, so noted that.

K: Um I dunno I'm trying to think We do, for the first one that we had there was, on the discord that we have there was a lot of study groups so I think study group wasn't the issue, but there also wasn't like any scheduled times, so if you can just make this one come to this one it was also hard to come to some. But htye definitely help because it's nice to talk it out with your peers on what they thing and just kinda look through the public exam and go question by question like, what could the fill in the blank be and it is like nice seeing the whole public exam but then obviously there is some parts that were withheld that makes it kindof like oh, it could be this could be this and sometimes me personally I know we're not supposed to look at the entirety of the public exam, like that's the part of it so we can critically think about each option and what the blank could be but sometimes it's like a little bit too like, I don't want to say too blank because obviously he can't just give us the exam but sometimes I'm like could be this could be this could be this And then going to all—I think I went to 3 or 4 study sessions before the exam and everyone would say different things and I was like oh I don't know what to think anymore.  And so just like a little bit confusing sometimes and when it was like actually on the exam I was like wow, I didn't even think about that.  So just a little bit yeah.

L: With the public exam, you said you weren't supposed ot look at the whole thing, what do you mean by that?

K: Sorry I just meant like the entirety of IF he gave us the whole question, and then all the [something]

L: Ok so just like the public exam can't be the exact test

K: Yeah yeah

L: Ok Gotcha.  Did that public exam version, so um it sounds like you studied using the exam in a certain way, like you were trying to basically fill in the blanks, does that sound right?

K: Yeah I was essentially  like—because I've never done this before, a professor has never partially given me an exam so it was a new thing for me so I was trying to figure out what kind of study habits I need to do in order to do well on the test So I kindof I would look at the public exam by myself, go through each question like what it could be and then I'd go back to my notes or rewatch a couple lectures or two, and kindof not find the answer but try to figure out what he could be talking about or what kind of topic he is trying to figure out in kindof a way So it's kindof an interesting like study kinda session I had over the last two weeks, it's interesting to say the least.

L: Interesting, ok, J how about for you?

J: Yeah I never had a professor who used a public exam either. To be honest I don't think I spent as much time with the public exam as K or maybe as I could have, I really just, I opened it and I looked at it and I was just like assessing whether the general subject matter of each question was something that I was familiar with and then I did kinda my own review.  For me personally I think like my best kindof review comes from practice problems and explaining what I think is going on and if something is wrong with that explanation then someone corrects it for me.  And if someone can correct it for me—actually the sessions we have every morning at 9:30 have actually been pretty helpful.  Like I think I understand the concept and then we get put into like a little breakout room and I'll explain the concept to my peers if they are struggling with it and then the peer facilitator is right there to correct it or Affirm it, and for me I'm just like that's a really good way of cementing my schemas of how it's working, um, the public exam, like, K kinda touched on this but it's just like if you really spend a lot of time on it you could get—you could look into it almost too much, you get almost confused or you like, you look at this and it's just like these certain questions could be so many possibilities and then he only gives you one answer choice and it's like ok I could spend 15 or 20 minutes trying to decipher this but I don't know, when I looked at it I just thought I could be spending my time elsewhere.  That's just my take on it, I know a lot of people really did benefit from the public exam just personally I didn't spend too much time on it And I didn't feel like it impacted my test score that much either.

L: So you said you like the practice problems, would that be like if there were just like a full practice exam?

J: A practice exam has been really helpful for me in the past uh so I do like a practice exam and like a key release, where the professor is like here, here's a practice exam from maybe another quarter or another year and then here's a key, just do it when you want and get an understanding of it.  I've also had professors who release not necessarily a practice exam but like a thing called a unit packet which is just a packet of Essentially applied questions and then the key it's more the professor explains how they were thinking through it and then also the peer facilitators or TAs contribute to that as well.  And then you can kinda see—you can do it more so. This is more helpfulf for like the critical thinking questions where You think through it and you think ok, this is how I would respond, and then you get that instant feedback or instant enough feedback when you're going over the key and you're like this was little off in how I was thinking about it, this is more the correct pathway, and I think Pathway-specific learning is really important specifically in physiology

L: Yeah yeah because so much is dependent on taking every step correctly.

J: Pretty much yeah.

L: K I saw you nodding, what do you have to say about [something]

K: I just agree with the parts where, I also think I spent too much time looking at the public exam because you kinda thinking about like it's partially correct, like All the information that we will see on Friday is like right in front of us so you're like I have to look at this deeply and study it as much as I can because if not, if you don't look at it at all the exam on Friday could be like a complete surprise And there's some questions that you won't see because you don't look at the public exam, but I do understand what J is saying when some of us could spend too much time and I think that's what I did, I was studying off the public exam instead of off my notes, even though I did come back to that when I was confused on like a certain topic, so I understand what he was saying that Cuz I think I did spend a little bit too much time just trying to figure out the little, like I said earlier, the little puzzle pieces trying to figure out both what the professors want us to think here, like fill in the blank and the correct answers would be withheld so I was trying to like fill it in kindof

J: Sometimes there just wasn't enough information to figure stuff out.  Other questions you could kinda figure out what they were getting at but I just felt like half that exam was like There's just no way I could really figure out what this question could be on because some of the questions are applicable to multiple body systems, for example.

L: Did it help you decide what to study at all, as if it were a study guide maybe?  I guess cuz it's like, it could be this system or this system, I guess I should study both these systems.  Just as an example

K: Um I think For me personally, I understand what J is saying when it's not enough information to try to figure out what topic they're having us try to study so it was just A little bit confusing just in that way because there were some questions where you could be like oh.  I think It was specifically like the membrane one and it had a picture of the membrane and you could tell what molecule they want us to try to figure out so that was nice, For me it was like a break because I knew that when we saw it on the public exam that's what he was trying to go for and I wish there were more questions like that because it's a lot less critical thinking—but I know that's very important in the healthcare field but it was just like a nice break to see that I know this is exactly what he and Dr Hennessey were going for So it was nice to see that.  And um, what was I gonna say, and the um.  The ones where it's like was it this system or this system or this system we can like-- it's hard to study so much based on just one question because sometimes it's forgetful and it's closed notes so its easier to just refer back to what we did study based on the system, because the system can have so many answers but then it's just one, like the large intestine or the small intestine, it's a little hard to remember those small details But I know it is important for us to use that thinking.

L: So, ok, in terms of—you mentioned, ok, critical thinking is a crucial skill for healthcare.  Was there some way that you would like hope that this course would teach critical thinking about these systems like this is how you're gonna do it in a healthcare scenario?  Cuz I guess once you're in the real world everything is an exam, right, you don't actually know the answer and no one is going to confirm it for you.  So is there a way that the exam could be more like an actual health care environment?

J: It would have to be more case study based, in my opinion, it would have to be like say maybe a 20 question exam is like 4 or 5 case studies and then you have, you know if you wanted to go that route you'd have to have maybe more reading, which could be beneficial for the public exam, you know you release the case studies and get people thinking about what--and introduce the patient and signs symptoms onset type things and from there there's different things wrong with the systems that they want to test on and so in that sense that would be pretty cool I think because it would be more like you're on the job, you're putting together the mystery, You're solving the questions from the concepts that you learned, I think that would be cool if the test could be written like that, I do understand that that's kindof time—you know that would take a fair amount of time to get a test that uses case studies and also incorporates a couple questions that are not so applied, so there's definitely a balance there. At the end of the day it's just, I hear Ben say all the time, or at least I heard it all the time leading up to the test, like testing is not that great of a way to assess thinking, but like what else do we turn to, you know our education system relies on testing to indicate whether kids understand concepts, and it's just like, It seems so—from the highest learning systems in college down to first grade when you're tested on your reading to move on it's all about can you test and like can you test well, and I understand some kids are not that great at testing and that sucks, because the system is against them you could say. But we gotta find a way to hybridize it or encourage—get rid of the testing anxiety, I really appreciate what you're doing because I feel like there's gotta be ways that we can at least improve it

L: Yeah we sure hope so, and I appreciate what you're doing because your input is how we find out what's working and what's not. And I'm actually gonna circle back because you like the case study idea is where you get to go a little deeper, and one of the things I have you saying earlier was that like some of the exam questions, you get that testing anxiety that time anxiety when you have a question that's really long, So I wonder if it would be like, well, if we had that public exam with really long questions beforehand, and it's not like you're supposed to work the answers before but you can just get int—you can find a bigger scenario, would that be stressful, would that be helpful? Both to taking the exam and prepping for the exam.

J: I would have spent more time on the public exam if they were scenarios because I feel like it would have been more directional with my studying, for example the one really long reading question that K was alluding to Was about pretty much like MDMA being cleared out in the liver instead of alcohol being cleared out in the liber, but just that reading was like, yeah I had to spend a whole minute or two reading and digesting those big words, but then I knew, well, they're going to ask me a question about how the liver cleans out stuff and what might happen if something's wrong with that and so that gave me a direction, so I appreciated that because that did lead to me studying about the liver clearing out toxins and leading to me getting the right answer on the test, so that was nice. What I was saying about the balance aspect, you get anxiety when they're all critical thinking questions, right when you just know it's going to be objectively a pretty hard question set you know question 6 thu 10 is going to be hard, and that's when that anxiety comes in, because you know there's a big reading associated with it, but if it's luke question 6 thru 10 is associated with this case, you've read the case before on the public exam, and then maybe question 6 is something that's more lower-level learning,

L: Like a warmup question?

J: Yeah, kindof like a warmup question so, then you're getting into it and you feel like you didn't just lose everything on that case study, like you built up to it and heck maybe question 10 is kindof like, shoot, what the word, dangit—like a diagnosis type thing.

L: Ok, cool yeah thanks. K, thoughts from you?

K: Yeah I actually really like the case study idea aspect because I remember that when I was also looking at the public exam I was like, oh this question is kindof a lot of words and if I was reading it during exam time I would stress myself out. So I was reading it before hand like during the time he gave it out to us so I was like oh this is not too bad, I know it's about the liver, so I would touch back with the liver, and then I got it right, so it's nice to like kindof know, cuz with the other questions it was like is it this body system, this body system, does he want us to go in this direction, Kindof like just a guessing game, and with the game you go with different things, and it could be these set amount of answers. And then

L: Like a set of answers, or a set of questions?

K: Like one of those scenarios, I think it was like the first set of 1 to 3 and it had like the picture For me the picture didn't really help me at all, it was just something to look at, I don't know if anyone else felt that way and the questions were just like—it was just confusing for me because I wasn't sure if they were talking about this body system or if they were going in this direction, and if there were three different things that it could be about and I would just go to those three different things and try to figure out what it could be, and if I couldn't I would just refer back to my study group and get their input but then As for the diagnosis questions out of the scenarios that J was talking about, and having like a warmup. Personally I think it's a good idea, or I like it, because at first it'll get you to grasp the concept while also—and if you know the topic you'll get the correct answer, which is nice because I find myself getting like A whole scenario wrong and I was like, I do understand this system though which is kinda discouraging. And I remember one question was about ketosis, or about ketones, and I remember studying that specific topic rigorously because I remember he mentioned it and I was like I don't really know what it is but I hear people talking about the keto diets all the time, so I feel like he's going to have a whole scenario on ketosis and just having one of it, like what is ketosis, I was like oh I know what it is and having that like one warmup question was Just like a nice start, even though it didn't go any further than that it was just about ketones and ketosis, so I really like that idea of just doing like a warmup question, and then having either another warmup question or having two diagnosis questions that are more critical thinking does lead you to thinking, oh I do know this topic why am I doubting myself and it gives you just like a little bit confidence as I am a college student taking an online exam. But I do understand that if everything was definitional and everything was a warmup question, the averages would be much higher. That's why finding a balance for these two would be really nice and beneficial for this class because for me personally Since a lot of it was critical thinking it was just, it gave me a lot of anxiety like I'm not gonna get this right because they kinda bounce off of each other.

L: Do you think that you could have practiced that critical thinking aspect more during the classes, or do you feel like you actually did practice that some?

K: I think I tried [something] balance but looking back at how the exams are structured, the first one, I think I could have done a little more critical thinking and that could have led me to more correct answers, and so I think my studying, like how I studied did affect how I did on the test, so depending on how much the exams change or if he does anything different I am probably going to change my study habits in Being critical in how I think and not just oh I have to figure out what he's talking about

L: Oh gotcha ok. So if the second exam, or the next exam is the same type of structure, same type of thing, how do you think you would change your study habits based on what you know now?

K: I think I would—as much as I want to look at the public exam I would try to shy away from it because I know it'll make me want to just do that puzzle piece thing and figure everything out. I do really like meeting with study groups because it helps me reflect on if I'm thinking the right way or if there's another way I can think about it. And I think reading into the text book a little bit more because I did read but I guess I didn't go into too much Depth, so maybe doing that more but also that's what I'm struggling with because I don't really know how to study for this types of exams like if I'm supposed to use or Supposed to utilize the public exam as like my study guide, because he doesn't give a study guide, so like the 10 other questions that are like the surprise ones, it's hard to see what the surprise ones will be. So I think having the public exam and then a study guide cuz when we have study guides in my other online classes I kinda use that as a Concept check. Like large intestine, ok, what are the functions, what does it do, I kinda do it in my head. It'd be nice if it was like—cuz there was only like one ketones question, and I was thinking there was going to be a whole scenario about ketosis and ketones so I just spent too much time on that. I am also trying to develop how I am supposed to study for this class and It is a little bit confusing for me right now but hopefully I will get into it by exam two.

L: Great. How bout you, J? Anything you would change about how you would study for the exam if it's the same way?

J: I was also thinking about what K said in relation to the textbook. I have kinda this love-hate relation with the textbook because yeah it can really provide some insight especially if there's like a subject that wasn't really covered that well in lecture but then it's like what subject wasn't covered that well in lecture, it's kinda hard to know that so I Personally didn't really use the textbook too much for the first exam, um. I was an, I'm kindof an advocate for just like, if it's in the lecture that's what 's going to be tested on so like I started to, build --in the build up to exam 1 I was kinda on a time crunch I was doing some extracurriculars that kinda took over that week so I didn't get a full chance to really study that well but in the past and what I would do probably for exam 2, Every day you have the lecture that you watch and answer the questions for but I have heard of, basically you watch all the videos again on times 2 speed so you're just whipping through the lecture and just you're just kinda listening for those quick points, like Ben or Dr. Hennessey they're touching on that and I know that, like I got that, like nothing of what they're saying is like huh? Or what was that? So as long as I feel like I'm with it, then that's kindof a good strategy for me. I guess also a study guide would be cool, but I would also be ok with like a practice of 20 questions too, like I wouldn't mind that. So probably just one of those two things. I know personally for exam 2 I'll probably just watch the lectures again and that'll be it. I'll definitely look at the public exam but probably cap it at like 15-20 minutes.

L: Wow, thank you good stuff. Anything to add, K?

K: I think it was a previous question you mentioned a while ago, and I feel like I just forgot to answer it, it was like how can you make the exam more as if it was a healthcare field. And One of my ideas is like the case study that J was talking about and then another part I was thinking about was like, watching all the inside look videos was really cool and then seeing how a lot of them are like team based, and I know it's not ideal for people to take exams together as a team because someone could be dominating the entire thing and they could all get that one grade, but I was in another class at UW and in a breakout room and someone mentioned that they did take physiology like a couple quarters ago or last quarter, not with Ben and Dr. Hennessey, and they were able to take exams in A group of three, and I thought that was really interesting and didn't really think about it before because it's an online exam, and when the person in the breakout room mentioned that I was just like huh interesting. Cuz I mean we can't do that so I was just like—I never like think about how different classes are structured at UW even though it's the same subject, so when you mentioned how it could be similar to a healthcare field, Or like in like the clinical care, I know that you are in a group setting and if you don't know something you can refer back, and also, I know I said this before but it was just really cool watching the inside look videos and everyone was like-- I think it was the physician who said I was never by myself, and even if I am doubting a certain procedure There's always someone to refer back to, and I am working with a team, and I—also it's not ideal for a test because there can be that one dominant person and they all get the same grade but I do think it's something to maybe look into, I know it's not ideal for a physiology exam but I just heard it from someone else that did take it at UW that they did take it as a group and I was like oh I never thought about that. But I understand where differences lie.

J: I think it's really funny that you mentioned that because like the whole intro bio department is like team group test, like Bio 180, maybe not 200 I don't remember, and then Bio220 those are all group tests so

K: Wow I did not know that I did the series it was all by yourself

J: I thought that, I took a couple, I mean a bunch of group tests and I…they went ok. I did like the aspect of having another person to resonate thoughts on, it was really interesting, The first group test we took we had like five or six members in our test taking group and then the second group test we realized that we had to split it up into twos, because there's just almost too many ideas, so like for example if everyone studied and everyone's more or less at the same level, Everyone's gonna be batting around ideas and everyone wants verification of their wording, so then it kinda gets like, it really becomes a time crunch actually because you're trying to manage group time and you're trying to manage, like, do you actually have stuff on your paper for your submission. And I actually thought the grading ended up being pretty fair, I didn't feel like I got taken advantage of and I didn't feel like I took advantage of anyone else. At the end of the day it does become an integrity thing and Whether your group members want to carry you on a test or whether you want to carry a group member on a test, but it's a good idea. And surely I do like the inside look videos I look forward to watching those, because they're cool, just seeing what real physicians or other people in the Health field are doing and the fact that they mostly say that they're in a team kindof environment, it's encouraging and it make me feel happy that I want to go down that path and I'm going to have a support team, like structure around me.

L: Yeah totally. I love those videos too, I got to watch some of them and I was like oooh I wish we'd had something like that when I was in school. We did not have anything like that.

J: I liked how the massage therapist was like the most in depth about her diagnosis, like I was not expecting that. But we had all these other physicians and then the massage therapist comes in and hits us with this very specific pathway I'm just like what, like why do YOU know that, but hey that's sweet.

L: Cool.  Well I'm conscious of time and I don't want to keep you too long.  This has been a very very productive discussion, lots of great ideas and thoughts from both of you.  So I do want to ask before I wrap it up, anything I should have asked or should ask students, or anything that comes up that you want to add?

K: I can't think of anything.  I thought all the questions were straightforward and to the point of the exam so I liked everything you asked, so I can't think of any new ones.

L: All right

J: Yeah I feel similarly, it felt like it was an on topic interview

L: Good I'm glad, and I hope that was a little bit of a de-stressor, too, after your stressful exam.  So you have my email, if you think of anything you want to change or add or a burning thought, just email me and blah…

***Interview #2 April 13 2021 @R1 site (Leah Lily interviewer = bold)***

**Better Mastery of Content and are better prepared basically I talk to a whole bunch of different students…and then o back over all those things later and try to find patterns across what different students are finding useful or not useful or different ideas students come up with on ways we can improve. Make sense?**

Yeah

**Alright, great, so my first question is how's Bio 118 treating you**

It's really good I actually really like the structure of the class and I'm definitely I'm usually not a very sciencey person I guess, but I feel like the way the class is organized makes it pretty easy for me and…yeah. I appreciate the way that both of the professors have it set up

**What's good about it for you?**

I mean ok the main thing for like just—it really only applies to online learning but just having everything in one place like I feel like the bar is set kinda low just because so many teachers do so many different things about how they organize stuff, like I have one teacher who just puts a bunch of random word documents in the files tab and you have to go through and sort through to figure out what your homework is, so not a good system. Yeah so. But I feel like a way that 118 is set up is honestly the best that I've seen in all my classes this year, is that under the modules tab there's a block for each week and within that all the lectures are just like numbered 1-4 and given the number of the week and then all of the materials are in there too. So I feel like that's honestly the biggest thing about the class is that, is just how organized it is also that it's all recorded ahead of time, I think that's really helpful, I think that's way more helpful than live Zoom classes for a science class because then you can pause and replay stuff immediately instead of having to go back and watch the whole lecture again.

**Gotcha, so like you're making sure that you actually get it before you move on in lecture**

Yeah

**That's cool, that's very impressive, a lot of students go through the lecture just blinders on, getting through it so you know what you're doing. So I have another nosy question, you mentioned you're not a science person, what led you take this course in particular**

Well I guess in high school I never thought that I was a sciencey person, I liked bio just find but I kinda hate chem and I hate phsyics, yeah, I don't know why just could never work that well in those subjects but um yeah this year I took—last quarter I took nutrition, an Intro Nutrition class just cuz I needed a science credit and I ended up really liking it so I might now major in Nutrition and Food Science, who knows

**Cool**

So bio 118 is one of the classes that you need for that and I'm then taking the next level in Nutrition as well just to figure out if that is the major I want to do.

**Those must really complement each other well, taking physiology and nutritition together**

Yeah definitely

**Yeah cool, and your interest is maintained, I hope?**

Yes it is

**Wonderful, yeah, that's great. Not to do that annoying high pressure thing but do you think that these course are actually doing you a good job of preparing you for what you think you might want to do with a nutrition degree? If you end up sticking with nutrition of course haha**

I don't know, I've been kindof all over the place, like I started out as an English major but I've taken an English class each quarter and I feel like my English class this quarter kinda solidified that it's not what I want to do, and I just had the opposite reaction to Nutrition and STEM classes which kinda surprised me, but yeah I guess that's kinda hard to answer because within nutrition I have no idea what I would want to do but it's the subject I found most interesting so far but career wise I have no idea, so I don't really know if it's preparing me that well but I do feel like—yeah I've gotten like a better understanding from 118 than I have from previous science classes, I feel like I've been understanding stuff more quickly and more thoroughly than before, so I think yes

**Excellent. What interests you about nutrition? What gets it for you?**

Yeah well I guess as far as science classes go, both chem and physics always felt super disconnected from real life, like it's cool that these things are happening but it doesn't really affect me in any way, um, and especially physics I felt like I don't really need to know like if I kick a ball the speed and the accelera- -like I don't really care, it doesn't really impact me in any way. Bio has felt like the closest to that, like actually applying to real life and being able to use it in a helpful way, um and then nutrition just feels like a subsection of bio I guess in which it's even more applicable. Yeah I think that's what feels the most interesting about it

**That's really cool, that's actually, I was a bio major as well and that was one of the things that kept me coming there because it just feels so important, like it's me life**

Yeah

**Ok cool so it sounds like this course is really working for you and that's sort of in contrast to other STEM courses—do you think that's because of the way it's being taught or mostly just because of the nature of the subject.**

Um well I took two Bio classes in High School, just the regular Bio then I took Honors Bio. I really liked the honors class but that was basically all lab based, it was very little memorization and lots of just how to do a lab in a successful way so that's kinda hard to compare but I absolutely hated my Freshman year of highchool bio class, that class was terrible, but I think that was all because of the teacher, like he was just really bad, so comparing it to that like the teacher for—I think for science especially I the teacher makes a really big difference, and so I've liked the way everything gets explained in 118, and I feel like it's actually staying in my brain more than other science classes have

**Well that's great to hear, Let's see I'm gonna push you on that—what do you think is actually, so you're saying the instructors are good, what are they doing that's making it effective?**

Yeah um well, just on kindof a basic level I think that the format of like having a powerpoint presentation with a lot of the information there and then also talking—like I know that's really basic and that's what most teachers do but I think that is the most like successful part of it and then honestly I think the questions that pop up in the middle of the lectures, I think those are pretty helpful um cuz usually, I dunno, I feel like I've never, I don't really fit into the categories of like visual or auditory learner, I feel like I need a combination of them, and also I find it super helpful the way the way they give examples I guess, like a lot of times during the lecture they'll stop and be—I dunno just like apply it to an actual real life situation, which I feel like helps to kinda like solidify the idea, cuz if you're talking about…like it's really easy to talk about science in a really abstract way, it's like you're talking about something on such a small scale that it's hard to conceptualize it and like apply it to what's actually going on, and so I feel like the way that that kinda stuff is explained and given Real world examples and applications of that stuff is super helpful

**Great ok, one thing we're looking at for this study in particular is actually the exam styles, so I understand you've had one exam so far, how'd that go?**

Honestly, not as well as I was expecting. Yeah, I felt pretty solid on the material ahead of time and then during the test I didn't feel great and then I didn't get an excellent grade but then I think that's part of that, just how STEM courses go, that grades are usually lower than in humanities, but one thing I will say about the exam is-- I don't know how like fixable this is but I really don't like the style of exam where you can only see one question at a time, and once you submit it it goes to the next one, and I feel like that goes against all the test taking strategies that like in High School everyone explained, it was just like if you don't know a question skip it and come back to it later, and you can't do that and I find that really frustrating

**Yeah**

And I guess some of that might be like if the second question answers part of the first one then you don't want to be able to see everything together but I feel like the questions in this class don't do that so I don't really understand why it has to be in that format, it seems like it could be in a like regular testing format where you could come back to them

**Mm hmm**

Um yeah. I don't know I feel like that is the main thing that kindof threw me off I guess.

**Yeah that's totally valid, that's uh that would really annoy me haha. Ok cool. How did you prepare for it?**

Um well I take pretty detailed notes, I dunno that's just kinda always been the only way I remember stuff is if I'm actually writing it down so like. Actually I know people always say don't just go reread your notes but I do find that helpful a lot of the time. So I did that. That was the main way I was studying and then also like trying to go through the test that was-- I forget what it's called, the like Public exam or whatever that was like given out, like trying to answer all of those. I looked at Piazza a lot too to see what other people were asking and like see if I could answer any of the questions up there

**Excellent, that sounds like excellent studying, as I say before, you know what you're doing! That's great. Yeah. All right so that was the first exam, is there anything you're going to try differently for the second exam?**

Um…I'm not sure, I've --like I would want to study in person with someone if possible, I've found like on study groups over Zoom I just feel like I just don't gain anything from them or yeah, I dunno, just something about—I feel like it needs to be in person for it to be—I feel like that makes it a lot more helpful so I might ask on Piazza to see if anyone's in my dorm building and hope that someone in the class is

**Hopefully we can actually do that**

Yeah cuz I took another bio class last quarter and it just so happened that the girl who lives two doors down from me was in that class and so we would study together and that was really helpful. Yeah I think what I was doing for the last exam was pretty good over all, I just need to do more of it I guess.

**Yeah that's fair, and there's always a time crunch, students are busy, it's a thing!**

Very

**Ok, ok, um yeah that makes sense . I'm gonna ask a little more about the public exam. Have you encountered any exam prep tools like that before in your school career?**

No, this was the first time.

**Did that seem, was that I guess significantly different than anything else**

Yeah um. I mean I guess to a certain extent it's pretty similar to getting practice questions. This was definitely more confusing, I dunno, there were a couple of the questions on there that as I was reading through it I was like, I don't even know what this is asking because the parts where they don't even show the whole question or all the answers

**Sure**

Yeah, I feel like there were a couple that could've been clarified more um. I feel like there was one, I don't remember exactly but there was one that was like how would this graph change if blank, and so that part wasn't filled in, that part was withheld or whatever they called it, and so I feel like that should be—like instead of saying withheld it would be helpful if they gave a couple different things that could happen to it, like a couple different options instead of just like redacted that part of the question. So I didn't find all of the questions like super helpful as a study tool just because some of them I spent like too long just trying to figure out what it was asking, or what it could possibly turn into as a question.

**So would you have preferred something like a practice exam where the questions are like**

Yeah

**Different but complete?**

I think so, I think, maybe just because I'm more used to that though, I think maybe now that I've seen what this looks like it'll be easier on the next exam but in general I would say yeah I probably would prefer the practice question style of study.

**What about a study guide?**

Um…I feel like since everything's online and it's all so neatly split up into topics anyway a study guide, like you basically already have a study guide. Yeah. I dunno I feel like if you just look at the titles in the slides, I mean at the beginning and end I think of each lecture it has a list of all the things that are in that lecture so I don't know that a study guide would have been all that helpful honestly

Yeah, cuz it sounds like you already had what you were using as a study guide which was the very **organizied layout of the course that you mentioned.**

Yeah

**Sso then, Ok, so you have like—when you were thinking about How to study for the exam, did you go sorta to the study guide quote unquote first, or were you looking at the released exam um to kinda see what you were in for first?**

Um I think I went through my notes first, and so I have my notes essentially organized the way the lectures are split up. So that was essentially like looking through a study guide with some extra information on there. Yeah so I think I did that first then looked at the public exam.

**Ok. And so it sounds like looking at the public exam did not change what you were studying or how you were studying. Is that right?**

Yeah not really, I think as I was going through the public exam I looked at Piazza with that to see if anybody had questions about the same questions on the exam, if anyone else was confused about the same ones

**Were people?**

Um a couple of them yeah. But I also looked at the exam on the earlier end so people hadn't asked a lot of questions yet. But I went back a couple days before the actual exam to look at Piazza again, so that was helpful.

**Yeah all right. And then taking the exam itself. You said you hated the one at a time questions. Anything else that came up for you during the exam?**

Um… I don't think so. I mean like the questions weren't exactly what I was expecting I guess

**How so?**

I don't really know why, um I guess I just noticed while I was—I felt pretty confident about the material and then the questions were at a higher level of application I guess than I was expecting, and I feel like it was at a like higher level than the like public exam to a certain extent. Yeah I don't think it was necessarily unreasonable, I think I just--it was just about like—cuz every teacher writes tests differently, um and so now that I've taken one I feel like I'll have a better understanding of what level I'll need to grasp something yeah for the next test.

**It sounds like the level that you needed to grasp it at is like pretty advanced, is that right?**

Yeah um I dunno I feel like, I ok, at the beginning of the course I remember them saying..there were a lot of questions about like what level do you think you need to understand like this specific material, and you don’t need to understand it at a super detailed level, but I feel like they zoom out, like they go the other direction more than I was expecting if that makes sense, like less of “which elements and which molecules are at play here” but rather like “in a person if these are the symptoms like what do you think is happening” like that kind of thing, and so I was expecting something a little bit more in the middle I guess, yeah yeah. But I feel like now that I’ve taken one I understand it more.

**Ok. Um. All right, so this is interesting to me because you’re saying that you need sortof a different understanding of the material than you expected you would need, and you also said you probably weren’t going to study any differently, and so I’m curious about you know—and it sounds like you have really good study habits, so I don’t think that you need to—now that you have these expectations that are different what do you think is going to change for the way you process the material or the way you approach the exam or the material.**

Um yeah I guess. I don’t know that I really have an answer

**Oh yeah that—these are not easy questions because that’s what makes my job fun because I get to stump you**

Oh is that the goal

**Well it’s not the goal but**

Yeah, I don’t really know, I feel like this exam—in a lot of ways I think a lot of college tests haven’t been that different from um like what I was used to in High School, I feel like I was worried about that a little bit when I came to college, like maybe I’m just good at doing high school not good at, and maybe it’s just a different thing but This one does definitely feel just like the application of knowledge is a lot like more heavily emphasized than the knowledge itself, I guess. Um so I think that I had an understanding of the material like to a certain extent, and I need to like go a little bit past that, like I feel like I could regurgitate I guess what they said but not necessarily entirely explain it on my own. Yeah which is why I think having someone else um would be helpful honestly-- my roommate’s here, I should just try and explain everything to her, I feel like that has been helpful in the past, just like trying to explain a concept to somebody who doesn’t know anything about it so maybe that will get added into the studying next time.

**Yeah, that’s an excellent solution. Actually some research recommends you try to teach someone else and you end up deepening your own understanding, so let’s pick on your roommate**

Yeah sounds good.

**Well I don’t want to keep you for too long I know you’ve ahd a very long day already. is there anything in the course that isn’t working for you, or something you want to complain about, any other thoughts?**

Um no, I don’t think so. I think organization of the class, in online learning I think that’s the biggest thing, honestly, regardless of how you’re teaching the material because I feel like everyone, teachers have their own styles of that but the thing that’s varying the most is just organization and figuring out what you’re supposed to do an I feel like this class has been the best at that. Yeah so I don’t have any complaints.

**All right so that's good to hear, so good luck with the rest of the course and whatever direction your studies take you**

Thank you

**Nutrition is super important and a fascinating field, you know if you end up sticking with that, great, that's a—a nutritionist can just be the best, I have Type 1 diabetes so I have seen a nutritionist and they're just like so helpful, they know so much stuff.  Anyway, there's my rave.  Yeah, one of the reasons I really like my job is I get to talk to the next generation of professionals, and I think about like—it makes me very hopeful for the future blah blah**

### ***Interview #3 April 14 2021 @R1 site (Leah Lily interviewer = bold)***

*[transcript recording starts…]* **make the educational experience for college students better. Make sense?**

Yeah

**Aesome. Ok to start with I'd just like to know how Bio 118 is treating you**

Mmm I would say it's getting a little difficult, or I think I think that it is difficult after we had exams

**Ah**

Though it was difficult I thought I could manage it but after we had our exams I was like oh this is challenging

**Yeah I've been hearing a lot that that exam was pretty hard it sounds like**

Yeah personally for me um I think it's mostly because of for Bio 118 at least for um like for us the lecture is like, there's questions, embedded questions in the lecture and the questions was…ok, so like I can do it. But when I went to do the exams to questions that are kindof different, like, I think it's more like application based which I feel like is ok because I have like a bit of STEM background but like um it's been really different because it's not what I expected it to be since the questions that we got um that we got for practice are different than what is out for the exams so after the exams I was like ok I have to change all the different ways I study for this class I think which is why I think it's started to get challenging because if not I'd think it was ok

**Ok, what are you going to change do you think, for the next exam**

I have to prac—its more prac—I have to do more practice if that makes sense for like When I was started out I thought it's more like memorization based or like you know, it's not really like how applying like I don't like, I don't relate bio 118 to like having to apply a lof of t hings like Having to apply what it is, It's an introductory course to like our physiology, I was like ok so I have to know where the liver is, what is the function and stuff but no, it turns out like ok if this happens what will happen to this, it had to like, it was a whole different way, I have to think in a whole different way, I have to make sure that the foundation that I have for the function of this like particular organs and like the enzymes, I have to like really know what they're doing in order to like apply it to the questions that are like the exams. Already exams are like ok the graph, it's about the graph I'm not really sure but it's about a graph, it's like ok, if this happens what will happen to the graph? So even though we got questions like this like in our lecture, it's like different from this because the questions are structured differently, like I remember the questions are structured in ways that are like oh this is testing my English

**oh no, Did the instructor give you any help for preparing for the exam**

We have like a public exam that are like I think a documents that are with yeah, but for I me I personally think that we don't get the questions, like answers to it that we can use, the instructors say like these are the questions that are going to be on the exam and stuff like that so I think that it's more kindof challenging that we don't get to see the answers because I wouldn't know what I did right and what I did wrong, like I can't be sure like is this the right way that I should study or if it's not the right way, so I was just like poking around, like putting best, like it turns out that it's not the way so

**That's frustrating.**

Mm hmm

**Is there something that you think the instructor could have done that would have helped? In particular it sounds like some of the wording of the questions was difficult, was there something they could have done to make that easier for you?**

I personally think that it will be best if like all the words could be more specific, because like for instance like for example for me even though I have a little bit of STEM background I didn't use it for a long time but for STEM I feel like if you don't practice it kindof like, it'll get away, so like I have to get it back so like I slowly transition into like doing STEM I feel like questions that are really broad like ok, so I remember I think it's the first one, the enzyme is this and then something like if this is wrong what will happen to the protein or something and I realize that it would be best if I can know like what actually is the function of the enzyme or what it actually does, like make it more specific so I can

**SNEEZE excuse me**

It's ok, so if it's more specific it can go through my brain more easily rather than like I have to think it again, ok like what is it and then after you know to go through all that thought process come back to this question, and after I come back to this question I probably just forget what it is again So I'm like really hopeless about this So if the questions could be worded more specifically like tell me like what does this enzyme like specifically do so I would know like if something happens so what is the result So yeah.

**So that's the exam questions, not the public exam?**

Yeah

**Ok huh. Ok, so more specific wording. That makes a lot of sense, definitely makes the questions clearer and easier to focus on. I guess my pushback I guess my answer would be if we're trying to prepare you for the real world of science or health care or whatever, that kind of thinking of thinking big then going piece by piece by piece, that's something we want tin scientists. Did that help you work on that kind of thinking at all**

I guess, I mean like yeah, that's the, if that is the kind of—yeah. I do like have to think small and then big again. So I think it's like thinking-wise it has for the exam-wise, no

**Yeah sort of unfair to throw that at you when you're trying to learn the course content?**

Yeah yeah because I feel like if it's for prac—when I do it for practice it's ok, but for the exam it directly affects my grade, I kindof like, oh, this is bad but it's…

**Ok, well that makes sense, because yeah since you're dealing with grades and it's not the real world yet you still have to worry about your grades so that makes sense. Did that kind of thinking, did that help you with the content at all**

Mmm so far mmm kindof like on the fence because I'm kindof like at personally probably best when it—not sure if this helps a lot but after the test you kindof like you know if you expect to do better but you didn't I'm kindof like that says like, oh man, it's kinda [something]. But I think that it does help me to understand the content, but not a lot because I understand it to exams. I realize it's like oh, this is like how I have to do this questions or like oh this is what I actually have to understand, like for this specific question or like this specific topic about the enzymes. So like ok so this is how I have to think about the questions that are going to come on the exams, or this is how I'm going to think about you know like this, like how I have to understand the basics, not just reading and memorizing and just know like what it is, but I have to like really think about it so that

**So it's, is it like you have to study towards the exam? Like they're going to ask this stuff so I'm going to learn it in this way**

Yeah yea

**That's frustrating because you can't just focus on learning it.**

T least that's how I think right now, it'll proablay change a few quarters in, I'll probably like you know if I get more confident I'll probably be able to learn more but for now I think my focus is kindof like ok I got to like—it's kind of like study for the exam way, I got to study this so I can do well on the exam because if not I wouldn't be able to do well on the exam?

**So what are you going to do differently now that you know how the exams go?**

I think rather than reading textbooks and all of that I would most likely, you know how the lecture would have the questions for su, so I would kindof like look at it—I actually would try to like rephrase the questions as well as like this is how the professor is going to ask this so what if I kindof like change this so like if this is positive effect, think about ok what would the negative effect be. A lot of this won't work but think about it in the opposite way of how the professor is talking about, ok so like the function of this-- I think we're learning about um like it's a hepatic system, so when I think about this one, so like this is the hepatic system, what about the other one, and make sure that they're related to each other but not you know, how do I say this, like basically just like kindof like do it like kindof like when the professors asking something I will also do like relate it to every other thing so that I don't, so that you know when they really change the questions and see the question again I get to ok,I know this so I will just like you know.

**You try to think about, it's sounds like you're saying you try to think about all the different ways they could ask a question**

Yeah yeah yeah Like if you get it correct how do you like say it

**Ok**

I tried to like guess, I'm not sure how but I'm trying

**Yeah, that's a really good idea, that's excellent studying strategy, so you know I feel like the course is difficult, it's very hard, and you're doing a good job of adapting to the way the material is working for you and I think that's really great. Um So where are you in your education career? Are you a freshman, sophomore, junior**

I'm a freshman—well it's my first year but I have sophomore credits

**--oh you're cutting out**

Hello?

Hello Sorry I missed the last little bit there

I was saying I have transfer credits so I have sophomore standing but this is my first year at UW

**Well welcome and congratulations! Ok so you have to kindof jump in to learning all this new style stuff**

Yeah, yeah I think that's kindof like kindof like exhausting, yeah I thought I was good in high school but The way I learn in high school was so different from like specifically Bio, it was like really different

**Yeah? So it sounds like um yeah um can you tell me about the difference a little bit? What's different between high school and now UW**

Ok so. I think so in high school first of all you have a teacher that you can go to whenever I can and like so, this is why I from the program ascpect. In my high school I had a super good teacher, sh'es like my bio teacher and she's like the best teacher in the whole world and you know I get to find her whenever I need help, so like I have—it's kindof like 24/7 help, it's like yeah, she's really nice to all her students, I get to see her and then not only that but like for Bio my teacher has like the best notes, like she lists out the things we need to study, like it's kindof like study for the exams but I kindof like know it because I'm in high school and back then we did--when we do A levels, I do A levels so like every day every word that you have to do like for at least for me, actually, all the teachers and A levels thing the things are for, like for A levels it's not really about your understanding, actually understanding is important but it's about key words they have to see these particular keywords. So if you don't have the key words even though the understanding is right you'll still getting wrong because you don't have the words that you want. So in high school we're kindof like going, we're studying for hte exams, like we have to learn these words for the exams but the understanding, you don't really need it, I mean that's wrong, that's wrong I said it wrong, It's important in your life, it's important in your knowledge but it's not important in the exams. So like if you go to the exams you know the keywords you know what the questions was, you get to give the examiner the answer that they want, that is basically like keywords, and you can ace your exams but that's how a lot of us did it, because We have like this structured way of learning so we get this, we give you this, we get this. We give you the answer, we get the um the grade. But when I come here, um the first few courses I thought like it's more like understanding, it's more important than on like the keywords and stuff, so like you have to understand how this works and then um like I have to like really know like ok what this is for me to like you know get to answer the question in the exams for like in Bio 118 now it's the university course so you know back in high school I literally could study for the exam it's like no understanding, no. It's keywords. Memorizing answers. I got it. But then now I have to not just memorize but answer and understand so it's kindo like oh no that's like a little bit hard. At first I thought I can just go by, like memorizing and then I did the lecture questions and I'm like oh it's actually pretty straightforward so I probably will be able to do it, so I studied and then like do the lecture questions again because I mean that's what worked for me in like high school because we literally just do the same thing and we kindof like know this is what they want, and so we get to be able to ok ok I'll be able to do this, Familiarize myself with the questions and the lecture questions and then I'll be able to do it in the exams because I'll see questions like this and I'll say ok this is the answer, but then the exam the questions are like so different than so it's more like application questions that'll be like, that's really hard, so  Different like you know, not just the lecture question but also different from how in my high school, because I think there's a whole different like US education system with other education systems that are like really really wide difference so I was like Good thing it's 2 hours or I would not be done, even If the exam is two hours I find it difficult to have to grasp it but even though I don't have to like even though I finished it within like an hour or something my brain the way I have to process the way the questions differently I have to like think back and then come back to the main questions again after I get every understanding, like this out of my brain, this is what this is what this is what, then I have to come back to the questions again, so I think it's like challenging but it's agood [something]

**Say that again?**

Like it's challenge

**Yes, oh yes.  Ok.  So so it sounds like the questions in lecture are more like what you were used to, the memorization, and then the exam questions were about applications.  My question is with the public exam did you have a sense of oh these questions are going to be different than what I'm used to, or was it still like a surprise on the exam?**

I mean for the public exam kindof like the professors will allow us to like ok you know what are some improvements that can be made and I think that's ok because I think maybe one of the main reasons I'm not confident even with the public exam is I don't get to see the answers, like I don't get to see what is right what is wrong so I don't know where, even thought he public exams questions are like questions that are like in our exams too, but some are in our exams.  So I thought like, ok, when I read it I was like ok it's a little bit difference but I get to find an answer but the only thing is I don't know if this answer is the right one, so I thought ok it's probably this answer is the right one [something] Ok it's probably the right one but then like some of it is not right, so it's wrong, but we get to know it, after I finished yes. Because it's like this, you know it's in the exam so we only know that we get it wrong in the exam, not before the exam, so I don't get to correct anything, so I learned, and yes.

**So it sounds like when you were going through the public exam you were, like oh, I probably know the answer to this question, I think I'm getting it right, I think I understand it but when you took the real exam it was like, oohhh maybe not.**

Yeah, and so I think there's like some of the questions are being corrected which is better too, but the thing is we don't know whether it's right or wrong when we did it ourselves, the public exam, because I think it's we are supposed to do it ourselves so we can get the hang of doing the exams, so that's what I did so I did it and you only know that it's wrong when the results are up so like these are the questions and I got it wrong, and some of them are public exam questions so I'm like, ok, it's wrong.

**Ok. Oh.  So for you it sounds like what would have been more helpful than the public exam would be like a practice exam that you can take and get the answers later?**

Yeah yeah that's what I think.  So if we're going to have a public exams, you know, maybe we could have the answers.  I know you can change the questions on the exam, it's fine, but at least for me it'll be better if I get to know how the exam and the answer that you wa—what is that answer that you know, that's for changing questions, if that makes sense, so like I could have a practice exam and even if you change the questions like you don't need to use the same question on the public exam on the actual exam, like it doesn't matter but at least we get to know this is how it's like and we get to have the answer like this is how we are supposed to answer this kind of question. Rather than I don't know how to answer that kind of question, like I thought I knew but mine is actually not he right one.

**Um ok that makes sense. Like having to rethink how yo'ure being tested**

Yeah

**So No please**

Yeah I just think that we don't really need to have the same exact question as on our exam, I think the public exam questions aren't questions on the exam, so we don't necessarily need the exam to be the same but it would be more helpful if you could give us the answer to the public exam and you could make probably changes to the actual exams, it doesn't matter, but they could be similar and we could just know that at least we get to have a practice of how it's going to be like rather than for us to sit in the dark with no idea, only like to get only to know it on the actual exam. I think that is like a little bit sad.

**Yeah, yeah, because you had to adjust to an entire new paradigm of what an exam is asking from you**

Yeah So like

**Ok**

Yeah, yeah, so I said like the questions on the exams are not straightforward or antying, even in the public exam, if we're not given answers, it's like Ok this is probably the correct one, because you know we are the ones, I'm the one who did it so I probably think this is probably the right one because I trust my own knowledge, but my understanding of my own knowledge might not be the understanding that is needed in the exams. Like for me at least that's the way

**So if I'm hearing this right, when you went through the public exam it was like, ok, I found this question about the liver, I know about the liver I know this this this this so I know this question, Ok. That's kinda what happened?**

Yeah

**And then on the exam, what was different, like**

So the exam. Then whoever—we'll use the liver question. Ok so for this is the liver and then something happens to the liver and they'll give you a graph. So then graph this, now this happens to the liver, what would happen to this graph if the liver is like this and then the other parts of something else so like maybe time, so this is time, and this happened, and then what will happen to you know the juice produced by the liver so what happened. And then so you just have to think, what what ok, so it is related, I mean after the exam I know that it's related because I know that they're doing it but before the--when I was taking the exam I was like, what is this? So I have to like you know and it's not like open book or anything too, so you I have to like really think back to what the like what should I do? Like I was like it's not open book it's not open note I just have to go back to the only knowledge I have about liver and stuff and be like ok, so I took a paper and pen and must sofrta graph it out so ok, this is what happens to the liver if x increase what happens to y, And then like it was for me I think it was really difficult to have to think x increase,y increase and have to think back to all my knowledge of liver and stuff so I think it's kindof difficult because like yeah

**Yeah, definitely difficult. So Thank you for describing that, that was really helpful for me because it's making me think about what different kinds of questions on an exam, what they're asking from a student and what they're asking is really complicated stuff because you have to take, you know the liver, you know the pancreas and you've memorized them separately**

Yeah

**And you have to bring them together in new ways**

Yeah, yeah, And not only that but but you have to add a graph in it oooooooh ok, there's too much things or me to like do! Or at least for me the'res like too many different ways I have to connect them together which I never thought, you know I never connected them before the exam because I didn't know that we needed to. When the exam you sort of realize that the questions on the exam every single thing is going to be interconnected. So this question can be asking about he brain, but they also connect it to like the blood so what will happen to the blood if it goes there, like what is the speed of the blood if this thing happened to the brain what will?!?!??

**Too much!**

Yesh, yes, so that is what happened. So it's not just like, if this is going to happen to the brain what will be the exact thing that we will get you know this illness or something, then what will effect the blood that is going to you know something like that. So it's like if this is the liver, what will happen to the pancreas blah blah blah Basically things like that, it's just like and then aaaagh. Now I'm doing the questions, I have to like take a paper and even though I have paper it's really though just this increase so what…and then I kindof like took a long time for me to have to know their connections because I'm like stuck at "so what" So like ok, so this increase, that increase, that's what the question is saying but what is it that you want me to answer, where should my thought go? So I have to go a bit round you know, thinking what is the function of the liver, what is the function of the juice stuff, what is the function of the thing they're saying, is like this organ is going bad or something so I say ok and I have to connect them all together again, and find one sentence answer it

**Right, yeah, it's like there are so many things you can think about and you still have to pick one answer kinda thing?**

Yeah, and I feel like it's probably gonna be easy if we have alike a lot of STEM background because that tends to make things easier, but like the STEM background is not really good and it's kinda difficult to do the application.

**I'll tell you—this isn't just you, this is difficult for everyone. Um all the students I've talked to actually were saying that the exam was a lot harder than they expected and that it was a lot of application and not a lot of memorization and that was different for a lot of people. So this is not your problem, like this is--you're doing something really hard you're taking this very complicated class at a new level and new environment a different language than you're used to and that's including all the STEM vocabulary that you have to remember from your previous studies and I appreciate you not just oidng it but going through it again for me. So, ok. Oh hello?**

Yes, I think I have to charge my computer

**Oh, ok**

I'm so sorry it's going to take like aminute

**Oh you're fine, just let me know when you're ready**

I'm so sorry

**You're absolutely fine, don't even worry about it. And I'm not going to keep you for too long because we're been talking a while so …if you do need to hang up now that's also all right**

Ok, ok I'm back,

**Hi, oh and your background changed, very nice. So what I'm wondering now, is because as I say a lot of students have the experience you had where they're used to doing a sort of memorize the thing then tell the thing for the exam, and they get surprised-excuse me—by the application that they then have to think about. And I know that for the instructors one of the things they though the public exam was going to help with was getting students used to that kind of question, but it sounds like that didn't really help. So I wonder—you're saying you would want to see the kind of answer you're expected to give, like would a worked example, like one full question being like "we're going to ask something like this, and this is the kind of answer we expect you to think through. Would that be something that's helpful?**

Mmm yeah. I think personally for me it would be yes, like what you said you get this like it doesn't have to be a lot, it can be like 2-3 questions and answers that are like, ok this is the application questions and then this is how the answers that we are going to—this is the answer, this is the question and this is the answer that we need. But I think it will be really helpful but it probably will take a lot of work so it will be easier if like the practice questions, so if the practice questions could guide us through how to think you know, like you know connect the thing, connect the dots, for example like you said this is like we'll just go back to the liver and the juice questions. For example, there's the questions and that's the answer, and there can probably be like a short guide telling like, how do they get to this answer, so like this is the function of the liver and the function of the juice and something and this is how they connect it to each other and it's kindof like a guide to guide us through one or two questions so we kindof know how to think that way

**Like that connection**

Yeah

**So first like, so so it might be useful for a professor to say, here's an example of a big complicated question, and here's an example of how to think about it. First we have the liver, and what do we know about he liver, and then we have the**

The whatever juice that is connected to it

**Yeah then we have that and what do we know about that, and what do we know about how they're connected, and with all of those things, how do we answer the question. Kindof like that?**

Yeah, eayh, So like you know like guide us through how to do your questions so we know how to do it, like guide us through it because at least for me it takes a while for me to know how to think about it

**Yeah it's new**

[something]

**Thank you that's really useful.  Ok well we're about out of time and I don't want to keep you too long, Thank you so much for your help so far.  is there anything else that you want to add? Um no? All right, and any questions you hav about what we're doing or anything else**

I do have like Will I, I mean I saw in the consent form that the researcher is actually my professor so my professor know this?

**Um so your professor is not gping to know that you participated.  By the time we use your data, I will give you a fake name in your transcript, so your data will not be associated with your name at all.  I am going to talk to the professor about some of the stuff we talked about because we're going to try to improve what we're doing but I won't bring up your name, he'll never know that you participated at all, this will not affect your grade, and by the time he actually looks at your data, like the hard data, instead of just me being like "hey a lot of the students are saying that the test was really really hard for them" like that's the kind of thing I'll say, and by the time he looks at the transcript the quarter will be over.**

When the transcript is done will the recording be destroyed?

**Yes, I destroy the video version so no one will …. Yes destroyed after 7 years ….**

Is the study for Bio 118 only?

**We're doing it in Bio 118 at the UW, we're doing at some different colleges as well.  What we're looking for we're lookin at the type of exam style We're trying to see what kind of preparation professors can do so that students basically won't be surprised like you were, which is why your input is so valuable here and really really helpful, I think we can have some good ideas of what will be more helpful for students like you, and that's exactly what we're looking for**

Thank you.

**Yeah, all right, and I will email you tow links to Starbuck gift cards, I made a mistake and I accidently bought $10 gift cards, so I will send you two of them for your gift card and you'll get those in your email shortly.**

Thank you

**And anything else that you think of, if those cards don't work please let me know but alsow if you think of anything else that you want the researchers to now about …. that you thought of you can just email me and I'll add that to your dataset.**

All right, thank you.  So If I have some other thoughts after the exam during the quarter I can just email you?

**Yup definitely …  blah blah**

### ***Interview #4 May 4 2021 @R1 site (Leah Lily interviewer = L, with Students B and C)***

L: …I interview you and get you talking about the course..looking for patterns…exam…what students say help them…data anonymous, no one affiliated…any last questions before I give you the first official question?

B: Don't think so

L: Well my first official question is How has bio 118 been treating you?

C: Um I can / B: Go ahead

C: Ok, I was just gonna say, um, I have taken a lot of other upper level classes so far, cuz this is my senior year, so Compared to like o chem and those other, it's been really good, the teacher really, I feel like he gets back to you quick and through Piazza he responds to you well, so pretty good overall so far.

B: I, ok I'm a sophomore so this I the first like bio class really or even like STEM class I have ever taken, so much different experience, I think it's been really challenging for me, um. I definitely love that the professor is so engaged, I feel like he's been one of the most engaging professors I've ever had whichhas been great and makes me excited and want to come to class and other things, However especially with the exams being so frequent I feel like especially on the quarter system that's been really hard for me, I feel like I study for so long just all the time which, I get you have to do so I think it's definitely been a little bit of a shock from some other courses I've taken

L: Yeah

C: I think since we're in different aspects, er areas because I've been through Biochem and they did it very similar where we'd have it every two weeks, have an exam, but um, and It's 25 questions multiple choice and I think just like being in person from that and now doing this I feel like it's easier but as like a freshman or sophomore it's definably like super intense, like 100% that's how I felt with biochem and everything

L: Yeah yeah I'm glad you brought up exams because that's what we're focusing on for this particular study, though any comments on the course we take those into account as well, but we are, I'm curious about the frequency and having to study more because um while I get that's like really stressful and kind of a pain I wonder if it's also useful to just have something that's just like all right, we're in it, I gotta be studying already…yes no like maybe hehe

B: Um I think it's nice to kinda have this like strict schedule like ok you know every two weeks on Friday we're taking an exam, I think that is nice and helpful to like plan and here's something to study for around there. And like maybe just because I've never really had classes like this before I just felt like I'm always studying and that makes it hard and there's the next day new content then you gotta study that, so that's been kindof a lot but I feel like C in your case I feel like you're definitely more used to it so

C: I agree 100% also, the only reason I'm not like super stressed out is because my other classes they're not doing exams but if I normally was in my usual Ochem, Bio and usual series it would be a lot because honestly 5 exams a quarter is a lot I feel like already having three with most normal biology courses is a heavy load because I personally, like you were saying, B, like how you were always studying, a week before my exams I like to do deep dives back into my studying so if I'm doing that already for 5 exams that's 5 weeks I'm studying heavy on top of my other classes so

B: Mmm hmmm I totally hear that I think obviously you can't just take one class and graduate in 4 years, a quarter, but I think if you could only take one class it would be a lot easier to take those deep dives like you were saying like once a week and actually go through everything, like The amount of content that we're going through um but I'm also taking 20 credits this quarter so that's a lot so anyways.

L: Wow, yeah that is a lot

B: Mmmhmmm

L: Yeah the stress of the exams is definitely a ubiquitous problem, like it's definitely not unique to you, um, but yeah we, so we do wonder about ways to mitigate that, and one thing we've been trying is the public exam system of releasing those questions early, and so I'm wondering if knowing what you're going to be asked, or having some information that way has been helping for you

C: It defiantly has helped me again, with my biochem where it was set up similarly they didn't do that at all and it honestly caused a lot more stress, that could have also been due to it was in person, and you're in a giant Room full of almost 500 people but I mean now that it's online and I feel like I could be alone and like really concentrate and also study the beforehand material that helps a lot I think.

B: I know for sure, I really like the public exam, I feel like it's just a massive study guide which is really helpful, because then for like the questions and then it has some parts of it withheld and You can really like come up with different scenarios and come up with all that and that really helps me study so definitely very thankful for the public exam, I think without it I would just be so lost because there's so much content that I wouldn't even know where to start

L: Great, good, that's really good to hear. I'm gonna I guess press a little bit on how do you use the public exam, like you said, B, it's sort of like a study guide, so what's your process getting that use out of that?

B: I, So what I've been doing is going on the Discord or just any other group um and then we all kinda problem solve and work through it together which has been really helpful so there's just a ton of different ideas to flush out one question and the possible answers and then just goes through that. So I think a group study for the public exam is best, I think by myself I'd be pretty overwhelmed and there's just again, there's just so much so I feel like having other pepole working on the public exam with you really really helps me

L: And just for clarification, so you basically, you sort of brainstorm as a group to see what the full question is going to be actually on the exam

B: Yes

L: and then do you try to answer that as well?

B: Yes and then come up with all the possible answers.

L: Cool / B: Yeah. / L: yeah, and how about you C

C: I kinda do that similar but I kinda, I'll go through on my own first and um with each part kinda think of different answers cuz a lot of times the question might be withheld and think like oh if it was this scenario maybe it could be this and this and do that for like each question, like I did that this morning and I'm gonna rewatch the lectures probably tomorrow and the day after then go back over it and see what I can fill in the gaps with

L: And it sounds like you don't really do that as a group

C: I yeah I don't do too much group work, I think it's because I'm also in anatomy right now and I'm doing a lot of group work in that. That's what's nice so you don't have to go to lecture so you can do it on your own time. I really enjoy that. With my other classes I just feel like I'm being trampled with three hour labs and things like that so it's hard to fit in other things so I've been doing it a little more individually but it's, but I've been doing decntly well so I think it's been working for me pretty well.

L: Yeah that's great, it's different, different things work for different people but it sounds like you both have found systems that are really effective for you, I'm glad, I really—I like my job because I get to talke to students who are just like troubleshooting and problem solving and just getting through it, because, college is hard, I don't know if you've heard, so um it's very inspiring to me to see people who are coping with all that and you're gonna be like the next generation of healthcare providers and whatever, it's exciting for me, I use healthcare providers a lot. So uh actually now that I've done some assuming, what do you want to be when you grow up?

C: I was premed but literally about a quarter ago I decided to do pre PA just because it aligned with what I wanted more, it was more the program I'm looking at is MEDICS and I really like how they're orientated towards serving Washington and like the NW which is where I'm from and that's where I wanna live, so, I'm trying to go on that path and that made me have to take what is it, physiology and anatomy this quarter together, and I also after I graduate I still have to take an extra class cuz I missed it, but it's all right I mean I'll get there eventually.

L: Excellent

B: Nice! Um I just got into psychology so that's really exciting, so that's why I'm taking this class. And I definitely want to probably do a masters after graduating from here, and hopefully something with children, not sure what that is yet, so yeah.

L: Excellent

C: Congrats of getting in / L: Yeah congrats,

B: Thank you, thank you thank you.

L: That's, that's very cool because I feel ike between the two of you I have sort of overlapping interest in sort of serving and healthcare but from very different perspective which is cool for me Because the next question I'm going to ask you is, has the exams specifically—and like the course as a whole—has that helped you do you feel prepare for what you actually want to do or is it just a school thing that you have to jump through hoops for.

C: Um me personally I'm loving this course more than any of my others, cuz the others I feel like it was just kinda jam information into your brain and this one I feel like I'm really learning it, and it might help that I'm in anatomy and I can apply that to my other classes but it's definitely things that I know I will be using within the next few months

L: That's awesome

B: Ok, I think I have quite the opposite answer

L: Great

B: Um.  I feel like I've been jamming just a ton of stuff into my brain, ha, and this definitely is one of the classes that I just need to take to fulfill a requirement, um I do enjoy the class I just think it's definitely something I just gotta check off the box for me and it's a lot

L: Yeah, it's definitely a very intense class, Sorry what was that C?

C: I was saying that makes sense, that was exactly how I felt about my biochem and ochem, I was like this is brain torture I just wanna be done with this

B: Yeah,

C: It really depends on where you're trying to go in life so

B: For sure

L: Yeah.  Oh great well this is fantastic because now I can get you to sort of argue it out

B&C: Hehe

L: While I try to workout well what is our class doing to serve students with such diverse goals and it sounds like a little bit of a different background as well, cool.  Ok so back to the exams, because one of the things we were hoping to be able to do with this public exam is go a little deeper and have something that feels a little more relevant on the exam.  And what I've heard from students is that this makes the questions really hard.  So I wonder if you have thoughts about that, both the difficulty and utility of that kind of question

C: Um, do you wanna go?

B: Um yeah, I definitely feel how it can be harder with you seeing the questions before, and I see it in like a new light and you sorta have to re flush out all the answers again, I feel like that can sometimes be especially challenging for me, like ok I feel like I've prepared for this as well as I can and now it's something and I just gotta do the whole thing over again which is sometimes pretty hard But then on the flip side of that it is nice to be like, ok, I know I recognize this and I can do the same process of what I've done before um so I do kinda feel the love-hate with the public exam but I do think it's helpful in the end.

C: I agree with that, I think it's really a balance between like memorization and understanding, you can't be memorizing the public exam because then like when they give you something you didn't expect you need to be able to understand the information, actually interpret it, because yeah if you memorize you're just going to put down the wrong answer and I've liked learned that with my biochem, They didn't really give us a public exam but it was kinda like similar where they like try to trick you—not that the public exam is like tricky but that almost like, I thought oh I have to memeorize it I have to memorize it and I found myself doing bad.  So maybe like for people who havn't experienced it um telling them this is NOT for memorization, you need to understand the under, like, the meanings of everything to really apply it to new scenarios.

B: Definitely that's absolutely exactly what I went through the first exam I was like, oh, memorize, easy peasy, no, completely bombed it, horrible, did not go well.  So then I tried more of like conceptualizing it and I feel like I understood things better and I feel like everything I studied made sense, still didn't do too great so that was frustrating But yeah public exam, not for memorization which I found out the hard way

C: I feel 100% that was me in biochem and I just did horrible my first few exams I really had to like learn how to go through it wo maybe if we have like, not someone like explain it but maybe a disclaimer note on it to be like, these are not for memorization. Just cuz people who haven't experienced it might thing they are

B: That was a big shock for me.

L: Yeah. Good thought there, something like "Here's how we're hoping you'll be able to use this public exam, to help guide your studying, like don't try to memorize it cuz it won't work"

B: Yeah

C: Yeah exactly word for word on the exam, it's just you learn the scenario.  I know that sounds dumb but that's what my mind went to I was like oh

B: No I did too yeah

L: Oh, good thing I'm recording this, I'll put that down. Um Something else you said, C, sreally struck me, you talked about like they're trying to trick you.  What is it—and that's something that we do hear from students, it's like they're in competition—not in competition but like in conflict with the professors.  I was wondering what are the kinds of things that make you feel tricked on an exam

C: I haven't really felt it out in this class but more of our higher level weed-out classes where the average is a 2.8 or lower, it's definitely they have to make the wording so intricate that it feels like you're being tricked even though you're not really being tricked but maybe in the use of double negatives in the sentence, maybe you didn't catch it when you're reading it but that's kindof a lot I have to deal with and I'd get bad scores just because I didn't fully read it, I'd go back over it and be Like, I didn't read it like as good as I could have, and I felt like I was being tricked because I was being graded on not being able to read a question as well as I could have rather than if I knew the content, that's kinda how I felt about it and like my biochem and all that

L: Yeah, B, I saw you nodding along.

B: No I definitely get that where sometimes I feel like I learned the concept but I just didn't read the question how it was supposed to be interpreted, that feels kinda hard

C: That's why I like that this class they allow you to edit it and look at it, you're like I have no idea what that means, you're like, cuz I know a lot of classes—not that they don't care but they don't take that extra step to see like can you understand this beforehand

L: Like they're not paying attention to the fact that their exams may not actually be testing the content because the questions are too convoluted.

C: Mmmhmm And like maybe I know now that I've been in science longer I'll say things and I'll expect people to know but they just don't because like they haven't gone through the classes I've gone through So I think a lot of the times in biochem I did running start so I was starting out as let me see I was a junior but not really, I was a freshman, and I didn't know what half the words I was supposed to know and I felt like I was being tricked but that's another issue kinda with transferring in

L: So like, saying things that other people don't understand like just using terminology or vocabulary that you picked up?

C: Um yeah I think more the issue was just double negatives and just wording, but that could also be another issue I guess there's a mix of different things, I'm trying to remember cuz it was two years ago when I did biochem. But that's not really—I'm trying to think if that's an issue in this class…for me I haven't had an issue with that but I've also been in a lot of classes. Have you, B? Where you couldn't understand terminology or anything?

B: I think this for me really is the first class where terminology I've been like ok, before I even try to answer like what are they saying, like what is this word, how does this related to that. Um so I think that's also been another of my biggest struggles for this class is just trying to figure out what they mean by something.

L: mm. And is that more vocabulary? Like from the course, or is it that sortof double negative type stuff that C was talking about

B: I feel like in the public exams and the exams in general I feel like I haven't felt like there's too many double negatives or things like it kinda felt like they were tricking at all, So I would say more just the terminology and vocabulary in general.

L: And then I guess ideally you'd be like, oh I'm gonna have to know this when I take the actual exam, like I'll learn it now

B: Yes. And that's where like on the public exam I can be like ok, so that means that.

L: Have there been questions that you've taken advantage of the editing process of the prereleased version?

B: I think I did, I didn't on the last one but on the first one I believe I did, um, and it was I think it was one of the general ones that I know a lot of other people did as well.

L: Did that work?

B: Yes. No it was really helpful

L: Yeah C sorry what

C: Oh I was just saying I haven't asked for anything to be changed I just looked over and it seemed like understandable so I thought it was ok. Different scenarios

L: And I'm curious now, especially for B, because you're not finding this class particularly relevant it sounds like for what you want to do, and so I'm curious is there something that is like the type of thinking where you're sort of like, I'm learning these concepts it sounds like very deeply, and learning to think from, like a foundation perspective, do you think that's gonna be—I mean it sounds to me like that's a useful skill so I'm hoping that feels like something you can hold on to and take from this class

B: YES for sure.  No I think after finishing this class I think I'm gonna be really grateful for the way it was set up and how yes you can to memorize the most, I feel like I have to conceptualize everything, I think that's going to be really helpful for me oing forward but I think it's just really hard because it's my first time doing it

L: Yeah yeah definitely.. how about for you C?

C: Um what were you asking again? I was just thinking like once you're done with this class all your others will be super easy, that's what happened to me.  I went through some hard classes now I'm like, these classes are super easy, I'm chilling.  What were you asking earlier

L: I was wondering if the way you think based on how you have to study for these exams in a very deep and sorta conceptual level is that gonna help you do you think, and that you can take that with you and learn that as a skill

C: I definitely think yeah because—I'm trying to see what you mean, I definitely will have to use these concepts specifically for me like wanting to go into the medical field So either way I have to like learn them and memorize them and keep them for future use and so right now, like in anatomy and physiology it's really important to me so I'm spending extra time like paying attention to concepts but I mean if I wasn't in my shoes and I didn't really need this class I can see how you wouldn't find it, like you really need to know the concepts and you need to know Um for the future like you might just run over and not try to understand it if it's not something you need to utilize for the future.

L: Yeah, so it sounds like for you um, you do need the memorization, whereas B not so much, like you're not really gonna be using, I dunno, the different, like the individual molecules that don't go into like neurobio or something like that but C, you're like yeah I'm gonna have to have these at my fingertips. Well that's good, I really admire, again, that you're sorta noticing like yeah, I'm gonna need this and you're doing the extra work for it so that's really great.  So ok.  I have two different students here with two different goals, at different points in their university careers um. I'm gonna make you do my job for me.  How could we do you think, how could we as instructors how could we improve an exam so it would feel more relevant to YOU noticing your improvement rather than for US to grade you like we're the big powerhungry maniacs that we are

B: Um.  I think, yeah, cuz the first two exams not been great for me, um even though I do feel like I'm learning so much all at once um and I don't know if maybe that would just be better on the exam to make things, I don't want to say not as detailed because details are important, but sometimes like the super super specific ones are a lot sometimes so I'm not really sure how to help that translate better on exams, um. I dunno.  C you can speak on that at all

C: I kinda feel the same way, it's just really hard to know what's good and what's not um I think for this particular class how the exams are set up I like how they are just because it's not really right or wrong answer and we've seen that our teachers had to go back and been like, oh people fought for these answers so we're gonna let you guys have it and it's awesome that he does that but since they're like concepts and scenarios you can have more than one right answer so I kinda like that and especially if you defend it, rather than this is a molecule, tell us what it can or can't do, like right or wrong answers, I like how they're set up so I'm not sure how else to change the exam for this class.

L: Yeah, I like that, where it's like your process gets the credit because if you've thought through it and it makes sense then it's right, even because I guess especially in medicine there's not always one right answer anyway

C: Mmmhmm definitely

L: Yeah, good, good. Um. Ok, so one of the things that has come up in other focus groups when we've been talking to students is that they have also been interested in healthcare, and they were interested in having more things that were drawn from real life, so like case studies and then questions about that, and I was wondering what you thought about that

C: I really like

B: personally, oh sorry go ahead

C: I was gonna say that I really like that cuz I'm going into the medical field but I know for other people like maybe half the class like B, she's not in that she wouldn't like that as much, so I think how they have it is nice, they have a good like separate like 10 of the questions are like around scenarios which is good like if you want to go into the medical field here's some like for your practice, but I know for other s it doesn't work as much But personally I do like them and they help me for the future.

B: Yeah for me those are definitely the ones I struggle with the most. I do totally get that for a lot of people that's really helpful, like especially you C where you can really benefit from that. For me and where I hope to be heading, definitely those are the ones I challenge, they challenge me the most.

L: And I'm kinda reading between the lines here when you say they challenge you the most, AND it doesn't feel that useful is my guess. Like it's a challenge that—ok I'm wrong?

B: Yesh, yes, though definitely I feel like it may not help me the most I do think like later down the road working through that type of problem will benefit me, maybe it doesn't feel like it right now. Trying to be helpful about it

L: Fair enough. Yeah, um so another thing that from my background as like someone who has been in education because I like learning and I like teaching, and for me one of the things I hate most about exams is that they reduce your learning to something that's just one number, which is silly. But I remember in the courses that I liked the most, exams were kinda exciting because—and not just stressful—because it was like, did I learn this, I like this subject, it's gonna be important to me, did I learn it and I would try to tell based on what my exam score was. And so my question is would, is there something that could make exams feel more relevant as like a personal tool to assess YOUR improvement, rather than making them feel oppressive and stressful?

C: I would think maybe having supplemental tests that are worth no points

L: Oh

C: So maybe with having an exam then maybe after either giving like tweaked questions that's worth no points that you can take for yourself and just see how you did on that because that alone would tell you, you just did your exam, you went over the correct and wrong answers now here's another exam just for yourself to see if you understand the concepts again but in a different like way

B: That's a really good idea

L: All right, so you like that one too.  I like that as well

B: Yeah

L: Cuz it makes it seem like the content is the important thing rather than the grade.  Especially because it's not actually graded at all.  If we did something like, I know some classes that had like mandatory regrades, where you can like do point recapture, would that be similar or does that sort of still tie into the exam, like points for everything philosophy?

C: I think that's also helpful sometimes because then you could go back and like, I know I've had teachers who they made us find the correct answer And then write out like a full paragraph on why that's right and why we decided the other over the correct answer and then explain why we were wrong or something like that and it's honestly helped me to really think, oh why did I pick that and then determine like this makes sense and why this would be more correct, or maybe explain like why I still think I'm correct

B: I think that's actually a really good idea and might be really beneficial to this class because there are so many concepts that you really gotta understand and that might really help me specifically too to learn better or if I thought one thing be like, ok so that's NOT right and here's why, or like if it is like here's why I think that, I think that would really help me to actually learn it so then by say the final rolls around and like I learned that and it's in my head for sure, not just memorized kinda thing.

L: Mmm hmm cool.  These are really good ideas.  Um.  I really like that like ungraded exam idea, I'm gonna bring this up.  Cool.  Well, I'm conscious of our time and I don't want to keep you here for too long and I do just want to ask before I wrap things up is there anything that you want to bring up or anything you think I should be asking other students who do these focus groups

C: Um I don't think so other than just like what we talked about and like maybe adding in that extra like exam or just a way to re-get your points if you thought your answer was like more correct, or it was just worded weird, a way to explain that you thought you were right because it was worded this way, something like that

L: Like something that To kindof acknowledge that part of test taking is writing a good test and that doesn't always happen

C: Mmhmm so like acknowledge you knew the content but just came out wrong I guess

B: No I think that would be really nice for the students and also be a way to get some points back, like I DID understand the concept but it didn't quite work out on a multiple choice test or something

L: Yeah cool.  All right well.  I guess the last item is business is your starbucks cards so…

### ***Interview #5 May 5 2021 @R1 site (Leah Lily interviewer = L, with Students Y and Z)***

L: …Any questions before we get started? Great. Well so I guess my first question is, how is Bio 118 treating you

Z: Um well Y you can go first

Y: Um Sure, I really liked Bio 118 so far. I haven't taken a ton of biology courses before I think the only other one I took was 108 and I think it was only like 3 credits and it was more introductory and just talking about evolution so it's definitely been like a hard class, just in terms of a step up. But I definitely enjoy the challenge and I think that especially like professor Wiggins has just done a really incredible job of the zooms, I know they're optional but the 9:30 Zooms every morning are just really really helpful for going through material and He's always there to help in any way and also like Piazza so I think just the overall involvement is just really key and has helped a lot.

L: Great well that's good to hear

Z: Yeah and uh I kinda have the same thoughts as well. I did not have any bio experience, I left Bio in grade 10 sophomore year and I did not study any bio in junior or senior year so this was my first bio course at UW. Um so yeah first two to three weeks were quite difficult because before this I was studying Maths And I think there is a lot of different, there is a lot of difference in approaches to both clasese and maybe the first three weeks that I took were just to reset that approach and kinda change a lot of things and I feel the first exam was also was a basis because that kindof helped me understand what sort of questions are going to be asked so I know how to study, what parts can be of importance And how to actually watch the lecture I think, there are multiple ways that anybody can watch the lecture but there are only very few ways that can actually help you score well um and that is different from understanding well. I think uh I think Bio is one course where you feel like you have understood um but that does not necessarily mean that you will be able to prove that on the exam. Um. I must say that uh the exams kindof replicate what the SAT is, in the two exams that have happened sometimes I can just see and tell that this will definitely not be the correct answer choice and um it's just right there, when people say that extreme answers cannot be correct, and if the option choice is [something] will definitely help you do this, that is not going to be the right answer, I'm just giving a very vague example. So that is the actual problem with multiple choice questions, and which is why it's not a great way to actually do the testing. Um But overall yeah the course is good, um the lectures are very important um I don't read the book. I, the first time I read the book was I think four or five weeks ago, I have not read the book as much as I have watched the lecture um so yeah, I think the lectures are important, they're good to watch, you have to pause and take a lot of notes. So yeah. And synchronous sessions, as well, I attend all of them so they have been very helpful too.

L: Great. Um I'm curious about—Y, I do want to hear from you but I'm curious Z, you said that you kinda got like a wake up call from the first exam and I'm wondering what specifically changed for you, the way you studied or the way you prepared for the next exam.

Z: I think I looking, so there are two things, one thing is that you kindof have to be clear on the big picture, but you also kindof have to know the small details as well. And if you know the big picture you will probably get 60% of the questions correct but the remaining 40% are small minor details that you kindof have to just remember somehow, sometimes they are not even connected to other details as well, so, of course that requires good memory, and good memory comes from revising before the exam, re-watching the lecture, writing while you take the notes—like while you watch the lecture and I think note taking skills have to kindof change, when you, as you progress so you will be able to understand that yes, this is something that might actually show up or this is something that might actually not show up. And yeah I think it takes time to identify that, of course nobody can be completely accurate, you cannot completely predict that yeah definitely this is going to be on the exam or definitely not going to be on the exam, everything's important and um some things are just a little more important, and I think the trick is to identify what is more important and then you can get a little edge over the others, because the edge part is important. This class is not curved, It's not graded on a scale, it's graded on how good the others are doing, so if you want to get that edge it's important. Had that not been the case the edge part is irrelevant.

L: Wow yeah, and Y I was seeing a lot of nodding.

Y: Yeah not to be for lack of a better term like creepy but I have been in many of the synchronous sessions so the one thing that pops out to me is when Professor Wiggins is going over the type of test, and he was telling us that we weren't going to be able to see the answers—or like not the answers but from the first question to the second question you can't go back to the first question, And how Z was talking about how that was like a bad process because you just like, it limits you and also if you have one question and you're stuck on it and you wanna have like time if you come back to it but you're just not sure and you wanna go answer the easier ones you DO know or you have studied well it doesn't allow for that, and I remember that you brought up that point and now our exams are like that but I was also just nodding a lot because the first exam was also a wake up call for me, and I think definitely Everybody in the class our lowest grades will be that first exam

Z: Yeah again I totally aree with that, I think the first one, I wish we could have like a practice run through exam kinda thing but I think dropping one exam solves the purpose, So that's good too, but yeah I think this class is online so I don't know if they can make it theoretical like free response questions, but yeah sometimes I feel like they could have asked different things, 25 questions is limited. And it's like you get it right or you don't get it right, so it's right you get 1 or you get zero. I think free response has more possibility of partial grading, so. Just because I got something completely incorrect does not mean that I do not know anything about that particular topic, so yeah, I don't think that's a great way to—especially biology, I don't think that's a great way to judge. Physics maybe, maths maybe, but Not bio! That's-- I don't think that's fair

L: Yeah, it's kinda like what you were talking about where if the answer is too certain if the words are "definitely" or "never" like that doesn't really happen in biology

Z: Yeah exactly so. I mean kindof you can use that to your benefit but if it comes down to actually testing real knowledge uh yeah, that multiple choice questions is not the right way to go in this class.

L: Mm hmm yeah. Um. Yeah, so I'm curious also so one of the things that I know the professors have been trying for this particular class and with these styles of exams is they do the prerelease version where they release some of the answers, and I wonder especially since it sems like the first exam was sort of a surprise and you had to Adjust to ok I need to approach the lectures in these ways, and I need to sorta study in these ways. We were hoping that that prerelease was helping people to understand what they need to do so I'm wondering did it help at all, or not so much, did it help the second time or Is it helping with the third exam that I know you have coming up—not to scare you

Y: I think that for me I didn't know how to go about it. I had never received a prerelease of an exam and so I was very used to things like going into an exam and not knowing anything and although we've addressed that that's not the best testing strategy or way to measure intelligence in general, I I think that although it was like helpful—and there are some like omitted answers as well, Which also was strange for me because you're not able to work out the full answer you just know like part of it, and so I think that that confused me a little bit but I think I just didn't know how to handle it. Like I went over it with a study group and I kinda skimmed it on my own but the understanding behind it wasn't quite there because it was like ok I guess I understand what this question is asking but I can't answer it in any format because there's not enough information for me to go off of.

L: So you could, it was sort of like a study guide but not a practice

Y: Yeah I guess, I think that I was just—because I had never seen it before I just had no idea of how to interpret it and so now when I looked at the second one, the diagrams he had on those ones those stood out more to me, because I was like ok, I need to focus on this part of that, and so when he just puts like kinda half questions I get the purpose behind them but I think for me it's easier to go over my notes again before the test or to rewatch the lecture to make sure I understand it or go to a study session or study group than the prerelease really does as much for me personally

L: Yeah and Z you're nodding at that one

Z: Yah I kinda agree with that the first time the prerelease was a little bit difficult to work with, um. Again because I had never seen one before. So what am I actually supposed to do with it? And I think that question was answered after I actually saw the exam, so, then I knew that this I the prerelease and this is the actual version and I was able to see what bridges I would need to get the gap, and I think that knowledge I applied on the second exam and I almost kindof knew the questions already, in the second exam specifically for certain questions where because the diagram was there and they kindog used the same diagram from the lectures so you could just go back and rewatch the lecture or even go into great detail of the lecture, actually write down every sentence they speak because that will maybe definitely Get in your head. So yeah of course the preexam was difficult to work around with in the first exam but in the second and I hope in the third as well that it's actually helpful and can actually help you direct your study in particular way. Although I feel that kindof defeats the purpose of a test because you are biased to only focusing on the parts that are going to be tested. And that kindof defeats the purpose but I guess it's the same you know, if everybody has that then I mean then the average will go up for everyone, if nobody has it then it's the same for everyone so, uh that's those are my thoughts on it. That you can direct your study which kindof defeats the purpose of actually learning

L: Yeah, that's definitely I guess concerning and the other thing that is striking to me is that—so I have a masters of education and I did it because I really like learning and I really like teaching and I sort of think about what if we started the system from the ground up and built it in a way so it would be accessible and useful for all students, So like, an ideal world, what would it look like. And one of my thoughts is that in an ideal we'd be preparing everybody, everyone for example who wants to take physiology and go into healthcare, Learns physiology and they don't compete with each other they just all learn. And yes some people are better than others at doing certain things but everybody gets ready, and so I'm wondering about the competitive aspect and it sounds like Z for you you're worried about Finding that edge and so I wonder if thinking about like your goals, what makes that something that you needed to concentrate on

Z: Um yeah actually the thing is that this class is required for my major

L: Ah

Z: I do not have to go into medicine or public health or anything related to biology.  It's required for my major and if-- the edge part actually only comes in because of the grading style.  If there is a certain percentage I have to meet for a certain score the last thing I will care is what others are doing in the class and what they are getting.  Or even the average for that matter, if I know that a 95% is going to get me a 4.0 that's all that actually matters to me, and then the approach is very different, then the focus is on what you are doing , and um the edge I think only matters in the grading style and the honestly this makes it very competitive, that you only know that there's only a few percentage that you're grade is dependent on, and if people have different skills then there will be people who have those skills and are being benefitted for it, and just because you don't have those skills, and even if I realize that, let's say that I don't have good understanding skills, And I know that I don't have them, I can try and develop them but 11 weeks is not a long enough time to develop those, so even if I do realize that I don't have the skills 11 weeks is a very short time to even work on them, I mean I would do anything but that's not how it works.  But if I know that there is a certain limit that I have to reach then I think I can work on them, I think I have to reach that level of understanding Or whatever it is to make it to that level, but in this case I'm dependent on others, my grade is actually dependent on how good the others are and I think that's not fair.

L: Yeah that does sound.  That sounds sortof counter productive to me so yeah that's a really good point, I will note that, thank you.  I'm actually curious, what is your major, if you don't mind

Z: I don't have a major yet but I'm thinking engineering school and studying HCDE and this is one of the prerequisites and I will also be taking Bio 180 in the next quarter

L: So you're doing that, like you have to apply to your engineering program and so the GPA is very important I know about that

Z: Oh yes definitely it is important, but it's not about the GPA, I think I need it for applying to my major but I think I chose this class because it's also related to my major in some way, it does require a good understanding of humans, and this class is important that way.  I'm also interested in it so it's just not all about the grade, but I feel that it is also very important, it's both ways I think.  It's very totally said so many times oh you should just go with your interest and just learning, but not in a class in Bio 118 where your learning is not all that matters, It always comes back to the point, your learning versus how much the others are learning, and I don't think that's very healthy.  I think weedout classes anywhere are competitive enough and the grading style does not make it healthy at all.

L: Yeah I hear that.  Y I see you nodding pretty empathetically too

Y: Yeah I think that at a competitive school like UW, like UW is my state university, I'm from Washington but it is a very competitive school and Ranked very highly in general and I think that there's enough competition to go around that not necessarily everything needs to be a competition as well as I think it's--Z correct me if I'm wrong on this I think it's the top 5% gets a 4.0 is what he told us

Z: That's right

Y: So that was really strange to me because I've never been in a class where I was told like the top percentage of students, because I could see to the advantage that if nobody got a 4.0, like no one outright got a 4.0 that then that would benefit those top 5% of people really just they would automatically just shift up to the top but in terms of everybody it also like if you're working in study groups with somebody and it's like, ok, now it's just you against everybody, it's like the 5 of you or however many people in your study group but it's still all of you against like everybody else trying to do your best and I would say the same for like synchronous sessions like people who show up for those are the ones who are showing initiative and then are able to do that and are going, But I have no idea what it would be like if I wasn't going to the synchronous sessions because I think I would be very lost especially to what other people around me are doing, and then I would have no reference as to like grading at all.

Z: Yeah that's true, that's actually true, um the synchronous sessions are actually not required but uh yeah it is actually very important and I think there's going to be a stark difference in 2 people with the same skills and one attending the sync sessions and one not attending the sync sessions and there'll be a lot of difference in their knowledge and their approach, Which is crazy because this class is supposed to be 5 credit hours in a week and that adds up to more than five, so right out 5 hours of sync sessions and then lecture are 45 minutes, roughly about 45 minutes each so that adds up to What, 8 to 9 hours of credit teaching which is not fair for a 5 credit class

L: Um yeah, yeah I mean its definitely a lot of work, we hear this from students who are just like so stressed out because there's just so much material. Yeah that's an interesting point of like your team, your study group against everybody else, and then I guess, it's like you're having to spy on other people and think like "do they understand this more than I do" even if you feel like you have a good understanding it becomes not do I get it but do I get it as well as everybody else. That sounds really frustrating because you can't just take charge of your own learning. Um hmm. Well so that's a grading issue, and Definitely important. I'm curious a little I'm gonna change the topic back to exams, just because um grading is hard, you know, in education we know that it becomes destructive both for an individual competing against themselves and worried about points, and as you have been describing, when the class is competing with each other. So we don't really know what to do about that and if you have any ideas please share because you know the—I think the idea of the top percent is exactly as you were suggesting, Y, which was you know it means that if the top 5% only gets an 80% it means that people still do get a 4.0 instead of just the top student gets a B which is very unfair…but um. In terms of the exams, for you in particular Z I'm curious about it seems like you have kindof a clear idea of what you wanna do, and so I'm curious Y, do you know what you wanna do with yourself, you know in the short term.

Y: I'm hoping to do psychology

L: Oh great, um so my understanding is that's also-- 118 is a requirement for that program as well?

Y: Um I think so yes but I was also just interested in anatomy and physiology so like I took anatomy in high school and I just wanted to take it in college. So I think it is part of my major requirement then I just--tentatively because plans always change so I don't want to necessarily say anything yet and I haven't committed to the program, But a neuroscience minor, and so the anatomy and physiology would be really key for that because that is all bio classes and so yeah I think it's it's more [something]

L: Great, that's wonderful because I have two students here who are interested in the material for its own sake which is great because what I'm wondering about is if the exams, because you suddenly have to apply your knowledge, if those help prepare you for what you think you wanna do with this content at all

Z: Um can you explain that question a little more

L: Sure yeah, it's a little convoluted. Um, what we're hoping for from an exam is that not only will it tell us whether or not you're learning, but also does it help you understand how you're gonna be using that material in your career. Does that show up at all for you when you're taking an exam

Z: Oh yeah definitely, yeah I would definitely say that because as I said 60% is actually big picture, maybe its just specific to this class because it's a lower level class and it's a survey of physiology. So yeah the 60% that has like bigger detail I think that's important even, just on personal lines I think it's important to know those things, you should know your body and I think if you know your body you are saving yourself from a lot of things that are just plainly lifestyle generated, so yeah I think the 60% of the material that pertains to the bigger picture on the exams I think yes of course, that's very helpful and again it's like back-tracing, so if I know that 60% is going to be a big picture so when I'm watching it automatically I will focus on the big picture and I believe—and this was surprisingly not, or maybe it was mentioned many times but actually after the first exam and first two exams it actually became more clear that yes, that 60% is big picture, I kinda noticed that pattern and then that's how I changed my approach towards lectures. I'm not sure if it was blatantly said that yes, 60-70% will be big picture. I think if it is already told that 60-70% is going to be big picture people will start looking at the lectures in that perspective, so that can alter their approach but I also believe that that will be giving away too much because I think from a class it's just not about what you're studying but also how you're studying and I think it's just very important for this class, how you study kindof makes or breaks this entire class. Like for maths you know it's definitely practice, if you don't practice you're not getting it. But that's not how this class works. So yes so the exams, to answer your question back to that, I think the exams actually do help because if I know that they require a big picture understanding and an overview understanding and then some in depth understanding, yes of course that's important and you should have knowledge about your own body, yeah.

L: Thank you, yeah and how about you Y?

Y: Yeah I would agree with that, I would say that somethings, well I would say probably specific to biology in general but also a lot of other subjects, they build on one another and so that's been very evident from the first exam to the second exam, Yeah we learned about things in the first two weeks and then we were tested on them but—and we weren't tested on them necessarily on the second exam, like that wasn't the focus, but it still is like the background to like THE next thing. So I think that's very key in not necessarily remembering from like exam 2 to exam 1 Being like, oh this is on exam one I should really know about that but it's like, it keeps being cemented, and you're like ok, this is an example of this term that I've learned earlier and it's a different type and a different body system that I'm learning about but it's still, like, Homeostasis, that's going to come up forever and ever and ever, so like that's going to be cemented, so I think in that sort of way being tested on it does come back to different things and He also includes I mean I'm saying he, I know that Dr. Hennessey is also teaching but I feel like I've mostly interacted with Dr. Wiggins so—that he puts more example questions if that make sense, so he'll talk more situational, he'll be like this and this and this are happening, what's most likely the problem, or what could fix that, or kindof leading us to the answer but not like giving it away but just to kinda cement that, so I think those questions are very helpful versus necessarily being like What kind of, I don't know, reuptake inhibitor is this? That's just memorization but when he talks about the situational things that's definitely something that I'm like ok, I can go back and base it off of my knowledge but then It's also my thought process of what is exactly happening, and I think that's going to happen a lot—I'm studying psychology to be a therapist so I don't know exactly everyday that I'll be dealing with something where I'll have to be like oh what exactly did I know from Biology 118 but it will come in handy for more general just knowing the body, and I think like Z said it's just Important as a human to know things

L: Very cool. Um you both used a little bit of different language but I wonder if they're, if you're sort of talking about similar things, Z you were saying sortof the big picture and Y you were saying like building on foundations.  Does that sound like kinda what each other was talking about or am I getting this totally confused.

Z: No yeah of course it's sort of related as well because if you understand the example of homeostasis, If you understand exactly what homeostasis is and just from the top of it uh if you understand the big picture you will be able to identify in other parts of the body as well, so even if it is say diffusion or diffusion through membranes it will start I think in um the digestive system but it was, it's used in respiratory system as well and so I think that amount of, that knowledge of big picture is what at least I was mentioning and The homeostasis part is actually accurate here, you kinof have to build on each other and if you go into the very specifics of the building on each other part does not actually happen, if it's very specific it won't apply to other parts, so yeah I guess, I guess your understanding is accurate.

L: Oh good, Y?

Y: Yeah I think so, I think that it's. I think that like the key difference between it is I think that Z was more talking about everything that we could grasp from Bio 118 but I was more talking about the timeline of it but I think that yeah we were referring to like a very similar thing Just in a slightly different like manner of thinking, if that makes sense

L: Yeah totally thank you that does kinda clarify it for me, yeah it felt really similar so I thought I would ask about it.  And something else that sort of struck me actually I was wondering Y when you're talking about sortof starting from here and building, is that referring to the course as a whole or the exam questions

Y: That's more of like the course material itself, but then it also applies to the exam questions because you're building on that knowledge each week and whenever, like--each lecture may not directly relate to one another like if we were talking about hormones versus the digestive system and we're only talking about the liver for example That might not be a direct connection, we were talking about at least in Bio 118 but in a sense that if we're building on the material like I might see something on the final exam that I have built knowledge from. So I don't think necessarily any of the exam questions require you to, I don't know that you HAVE to build on, it but it's a natural progression, it just like happens.

L: Cool, I think that's probably something that the instructors would want, so that's good (cough) excuse me. So I'm thinking about something that I think you both have touched on which is that this course is sort of teaching you, it's teaching you how to study for this course but it's also teaching you how to study in general, and it sounds like—and again, correct me if I'm wrong here—it sounds like you both feel like taking this course has helped you refine study habits or learning habits. Yeah I see nods

Z: Yeah I have to say that--because it's my first year, I'm a freshman, I have taken two math courses, a philosophy course, a psychology course, a nursing class, and uh I did not really have to change a lot of things in my approach when it came to those classes, but I kinda had to change a lot of things when I started to take this class. And psychology it was psychology 101, which is kinda close to biology, it's not he same but it's somewhat related to each other, um but that class was kindof a breeze for me and this class, yeah it, it was, I think it was just difficult to kindof—you just had to change your approach a lot maybe because, I'm sure you find somebody else to say the opposite thing, that for psychology it was so difficult to have my approach work and then for bio was so much easier, I think it depends from person to person a lot so, specifically for me the changing approach part is important and I think there's only a certain permutation combination That you have to use when taking college classes, I think, I think in a few more quarters I will have reached all those permutations combinations and then I will have all the kits, all the tools in my toolbox for any other class as well so. I guess if you ask the same question to maybe a junior they'll probably say oh no it was totally fine because they probably have already taken a lot of different classes and know a lot of different approaches and I mean we're both, Y are you a freshman as well

Y: Yeah

Z: So yes I think our responses will be based on the limited number of classes that we have taken, so yes, I'm sure that every class requires a different way of approaching it, and yeah, I think if I've taken 118 I think I might do ok in 180 as against to straightaway taking 180 I would be saying the same thing or even more things because 180 is a little more difficult than 118. So yes I'd say that this class might possibly be preparing me for other courses That kinda require the same approach.

L: Uh, you're guess is correct, some of the uh uh students who are further along in their college careers have said yes this class would have been really hard but I've taken like biochem and that was the reall hard one, and now this one is ok, That was good because it means the rest of your classes are going to go like that (snap)

Z: Well that gives me a lot of confidence in the future so thank you for sharing that with me, thank you so much

L: Of course, how about for you Y?

Y: I don't know if you took any AP classes, Z, but I took some AP classes in high school and for me it's just a personal note I'm a very bad procrastinator, I tend to be A perfectionist at certain things and I like to aim for the highest that I can, strive for, and that means I avoid a task that's not necessarily that difficult but I wait until it's like absolutely necessary to do it

L: Spoken like a psychologist

Y: Yeah, based off of taking my AP classes from high school I was able to, I think I was able to get away with certain things and also just because it was a high school level and not college level, even though I got college credit for different things, and so I took AP psych which counts as pysch 101. So I would say that it's much more intentional and I think Z referenced this a lot earlier but it's more of like 9hrs a week, depending on what you're doing I think you touched on—like you havne't done the readings, if you're oing the readings, which I'm not caught up on them either but if you're doing them they take an additional like at least half hour, if not more depending on if you're skimming or taking notes on them, But it can add up very quickly depending on what you're doing so I would say I don't know that the class itself---the difficulty level, but my level of intention in studying and making sure that I'm doing different things and that I'm interacting with the material on more of like a daily basis rather than like Oh I haven't really done anything for bio this week maybe I should pull out my notes, no it's definitely like an every day working on it, and I would say that it's maybe not the level of difficulty of understanding because we tend to go a lot further than that, like as a class, we'll ask questions during the synchronous sessions that Dr Wiggins is like well That's like way beyond what we're doing but I'm gonna answer that because it's fun and I think that Biology is cool and so that's kind of a neat feature that we're going like Beyond that and so um I would say that it's the amount of course material that's just like, you just have to take it and that's part of like the quarter system as well like we have 10-11 weeks to get all of this down and so I would say I just have to be more intentional about what I'm doing And be aware of how much I have to learn in a period of time.

L: Mmmhhm

Z: Yeah that's, I agree with that yeah. Uh yeah this class, the class is at 9:30 but you have to watch the lecure before the class in order to actually be in sync lecture and also for participation points, uh so usually I watch it over the weekend and only then I'm able to attend the sync lectures in time with full preparation so, yeah I think it's it can be a little too much for a 5 credit class which is probably why I do not do the readings, there's only a certain amount of time I can assign to a class, it's not about having 24 hours in a day it's about How much you can actually do in those 24 hours and how much you would want to invest in a particular class because you have 2-3 more different classes other than the class that you have. Yeah I think, I also feel like I the lectures are made in a way that a sync session is not required to get a better understanding of the lecture, and I can actually, I say that because sometimes I feel in the lecture there could have been easier ways to either teach something that has been taught or to either structure it in a way that it was easier to relate with other things and I'm saying that why because I'm actually an [something] student, I'm from India, And the education system in India is very different, they will actually tell you things and you can actually draw a conclusion very easily because of the way things are structured, it's called spoon-feeding and I think that's not done here, there's a strong difference between the education system and the way teachers teach in India and the way they teach here, and I can so many times easily spot that There was definitely a better way to actually explain this concept that you've just explained and I can spot that because I have been wired to understand things in that way, I'm from India I studied 18 years in the Indian education system, so So many times I can spot that yes, if this was being taught by a teacher in India, he or she could have approached it in a very different way and a sync session might not even be required after the way they've taught it, but I guess that's how things are different here, That you have to establish those connections by applying them, and it—I'm not complaining or anything, I'm just explaining the differences, and so many times how I can tell that yeah a sync session might not have been required if the professor had chosen to just kindof establish a connection on himself Like just like that um

L: Like as you say like spoon-feeding telling you like this is how it connects instead of making you work it out for yourself

Z: Yes yes exactly but I understand if that is not the purpose of education that that is here. I think You kindof have to do it for your self in order to actually succeed and I think this class specifically focuses on that, how many connections you can establish by—not even unrelated but just things that are given to you differently and how many times you can just seamlessly see ok, yeah, this is exactly why this is happening and then this is happening. Structure, probably I would say, could be a little bit different, so that a sync session is not needed at all. Um.

L: So, so um it sounds like some of what—this strikes me as similar to some of what Y was saying where you have to apply those concepts across different things, like we learned Homeostasis here, and then we learn it over here, And it's like what you're saying is that you could teach it together and then the students wouldn't have to do the work of saying oh it's this and its this, oh it's the same. Does that sound like what you're telling me?

Z: Um it's not more about how you can cross reference so lets say I'm in week 6 and then they teach you something and then I establish a connection from say week 1. It's also about how much I can even in one lecture so many times I can actually tell that it's good that they explained it in this way but I believe there was a different method possible that you could have well structured your lectures. So many times the reason that somebody has to go back and watch is either because they're not paying attention or because they have not understood. And those can be two interrelated things but More often than not I can actually tell that I have just not understood because it was not structured properly And then I go back and I structure it in my head, so I do it on my own because the spoonfeeding part does not come in here. And I can just draw that contrast because I have been in a different system and now I'm in a different system, so this might now be relevant to what…

L: This is fantastic, first of all its really interesting so I love to hear it, and you know, you know we certainly in the world need to be learning more from each other so different systems is great.

Z: Yeah it's just that I'm used to something different, and this is different so yeah, I see that this is something that could have changed and I could have understood better.

L: How does that strike you y?

Y: Um…the one thing that I would say is I also have like a different background I guess I was adopted from China but I was raised in the US since I was 1 so that's all I've ever known, but I've been aware of it because I grew up in a Mostly white town, like it was just not a big deal but it was what it was and my family is white who adopted me, so there's that distinguishing bit so when I came to UW it wasn't necessarily that I was more aware of my identity but it was definitely the idea that--And I think because I've taken political science classes as well, that has been emphasized, the idea of yeah we're at uw, we're learning in Seattle, in Washington, but That we should be looking at more than just Western ideals and I don't think I had really looked at it like that before, I hadn't necessarily been like, I don't know, I don't typically think of something and say Oh I think this because I am an American and this is connected because of that, so I think when we talk about not just international issues from a Western perspective but We have a lot of international students and we can hear their first hand experiences I think that's much easier and there's like the emphasis that's put on it and the respect Is really nice to see as well as I would say the one thing for me that stuck out about the lectures and like the order of them is that between Dr. Wiggins and Dr. Hennessy they're very very different, and one week is one teacher and the next week is a different teacher and their styles of teaching are very different but also, not just them interacting with students But also I think kindof Z was talking about earlier but the structure necessarily from Point A from Point B to Point C Dr. Wiggins is very different than Dr. Hennessey and he tends to go much more in detail about certain things And um I think it's supposed to be like 4 questions that we answer every lecture, he usually has a lot more that we answer as we go about them and she typically has like 4 or 5, and I would say that's the biggest difference for me.  But um yeah I think that that's really really interesting to hear coming from a different country, and my only other experience with that is I've hosted exchange students so they were from Indonesia and Pakistan so that's my only other reference points, so I think that one of them, Kiren from Pakistan was less prepared than Isna who was from Indonesia so it was interesting to see their teaching styles because—and I was a lot younger, second grade maybe, and Isna came to our high school and it's not super super rigorous It's just average level of difficulty and She found it really really easy and the only hard part was really the language barrier, whereas when K came and she's from Pakistan it was much more difficult for her as far as how the class was set up as well as the language barrier so I would just say, I think each country tries to emphasize education but we all go about it such different ways that its hard to just like put them together, Like there's no good shift from one to another, it's not like natural it's just like oh yo have to adjust and adapt on your own to this new system

Z: Yeah I guess, I think Y think that's a good way to put it, maybe it's not just me taking the first bio class but it might be me taking the first bio class in a different education system, that can also be a huge factor in my responses. So yeah I think the background part is necessary, That was important to point out, yeah

Y: I also just think it's really admirable, to just even if it's not always a language barrier or culture differences-though there's culture shock even if you come from like the east coast and you come to Seattle there's culture shock, but just it's I dunno, it's just very admirable because I were to think about going to a foreign country, I studied French in high school and If I were to go to France and suddenly try to take college level courses I would be so lost that I just yeah I just think it's very admirable

L: Yeah

Z: Thank you

L: Oh my goodness this has been such a fascinating discussion with two very insightful students who are clearly, you know, very hard workers and very invested in what you're doing which is just, this is the reason that I love my job is because I get to talk to the next generation of creators and workers and professionals, and so thank you so much.  I don't want to keep you too long and I'm conscious of your time, we're almost at an hour, oh my goodness I feel like I could ask so many more questions, but again, I know, as you have told me, you are very busy so I'm gonna sort of wrap it up here.  Unless there are any last things you want to share or if there's anything you feel like I should be asking other students who come to do this study

Z: Actually I don’t think so, I think I’ve spoken a lot of things but I think things like these go better if the person whose asking questions asks the right questions instead of the person who gives the right answers, and you have been great with asking questions obviously, you did not really, I mean if you have more questions we can probably schedule another hour but that would go differently so yeah but for now I don’t really think there’s anything I want to add um to that.

Y: I think that I’m good and for my own personal curiosity I might add like, experience in education because this is just again a personal kindof more thing but I come from a family of educators so ranging from my grandparents and my mom and my aunt they have all been an educator at some point in their life up to like from a teacher, like paraeducator up to a superintendent and so my background in education is very different and how I look at especially like policy, because I did like programs in high school for whatnot so when you said that you have a masters in education it piqued my interest because of my background in it and I avoid education as a topic sometimes because I know I can get very into it and heated on it if someone’s going to debate me on like specifics of it just because I have so much respect for education and people who work in education and I think there’s a big discrepancy about how much we—especially speaking as just like the US but we put a big emphasis on education and there’s not as much follow through so there’s the idea that you should get a high school diploma, definitely and if not like a GED, and you should go to college, and There’s not as much follow through and you’re kinda left to do a lot of it on your own, so I would just be personally curious to see peoples background like how much emphasis their Parents put on education and how much it was personally like they were like oh yeah I really wanted to come and study these things and I’m really self motivated and so just like that kind of factor but I don’t know how much it would relate to the study more than just me being interested in that perspective

L: Yeah it’s definitely in these sort of comparatively short windows I have to talk to students often it does just come up as students talk about themselves and their backgrounds and what influecnes how they study, like it came up for both of you as just part of the conversation, And I think you’re absolutely right, it’s very important because how people view education or what education system they come from or just how they personally feel about the purpose of learning or college or things like that, there’s just a Huge variety and so it is defineatly very important, again I say very insightful, you both have been just, it’s really been a pleasure, and so I will wrap up. Your starbucks cards … etc

### ***Interview #6 May 7 2021 @R1 site (Leah Lily interviewer = bold)***

**…And What we're specifically working on for this study is actually trying to improve the exam system, um and I understand you uh just had one**

Yeah just walked out of one

**How did it go?**

Oh um well I don't really know how-- I can't, I usually can never say after an exam cuz I'm just afraid to jinx it um but cuz right now I got, we have our grades but sometimes like Ben goes back in and fixes the grades like to, like sometimes it's not or a lot of times he tells us it's not accurate in that some questions have multiple answers and he just couldn't put down multiple answers there so he just goes back so He fixes it so like the answ—the grade is reflecting the correct answers. Um but like I don't really know yet I did feel like traumatized by it, but I felt like it really made me think um haha. For like a Friday morning but there wasn't like there wasn't much hassle I guess that's the word for it, um, it was pretty easy breezy in terms of process.

**Wow there's a lot to unpack there because you say it's easy breezy which is great but you also said you were kinda traumatized by it so I'm curious about like**

Oh I guess just for exams, it's just something I always, something I always say about exams just like traumatized by exams, the exams are like always bad even if they're easy or even if they're fun like it's just like I guess student culture, it wasn't like traumatizing it's just something I say

**Oh yeah ok, well that's fair. I would hope you're not actively traumatized by an exam because that--**

No

**Ok all right yeah I that's I mean that's as educators we know about like exams are stressful and I think um we all wish we could find a better way to measure learning. Is there something that has worked for you to make it at least less traumatizing?**

For exams?

**Yeah**

I guess one of the things that the—one of the upsides of the pandemic was that exams, like you can take exams at home and in the comfort of your PJs, you don't have to wear your makeup, you can wear, like you can be snuggled in your blanket, I think that was the best part of the pandemic for me being able to take exams in the comfort of your home and it makes it, it makes it like after exam you like kinda close your laptop and you're done with it and you're not like kinda still in like the clothes of school., like that mindset of school there's still so much to do you're just kinda like "I'm done with the exam, I can go treat myself, I can go do like things that like that comfort me". Um but like I think that's like I guess like the pandemic is not really a good like haha solution to like getting traumatized for an exam but I guess like that specific like being comfortable like taking exams when you're comfortable is definitely a good solution for helping the trauma ahaha

**Can you think about like different types of exams you've taken across your academic career and like were there some that were more upsetting or less upsetting**

Yeah, um, well this is definitely like—I know like Ben, I've taken classes with Ben in the past before, like before the pandemic and this is his, This is his style too, like I think 2019 I took Bio 200 with him um and I just thought the style of um of having, releasing the exam and having students like work with each other and like kinda wonder you know, what is this redacted part of the exam, wonder like together and then like that studying is just really helpful. I think that, that's like a really powerful, or I've always thought that was a really powerful way to like study, it's like when you don't know in this so you just like explore all these options and there's like so many possibilities and so many avenues you can travel, and in that way you even like learn some more, I had—I studied with friends in Bio20 but it's like the same exam structure, um, I studied with friends then and it's probably like some of the best most productive study sessions ever, because you're just like wondering and you kinda know what's on the exam so it kinda alleviates the stress sort of, um, whereas in other classes I've taken um I've taken like basically memorize and regurgitate exams, like classics 205 I guess, it's you memorize all these terms and then you like, the test basically—it's basically like quizlet but you don't have access to your notes, test is like, what does like, I dunno, like what is a certain suffix or a certain base mean, and then like there are four multiple choices and you pick it out. Those kinds, those are pretty like straightforward um I guess those are less anxiety inducing and more just put in a lot of effort, but for other STEM classes I've had, like Ochem, um, like it's like sometimes you never like they give you a preexam or not a preexam, a pretest or like a practice test, and um you think you're doing good and then, and then they hit you with the exam and it's just super hard and it's like, it like is completely different than the practice exam so you can never really know with a practice exam, so that's why I like Ben's format, because—it's not like a practice exam but it's not like completely different from the exam so yeah that's –I feel like those are the three main types of exams I've had, it's like, regurgitate exam, like memorize and regurgitate, ben's exams, and just the other exams where they give you a practice test and then like an actual exam. But yeah I, I 'm a biology major so I've taken a lot of STEM classes heheh

**And are you a, let's see if you took Bio200 a couple years ago you'd be a junior maybe…**

I'm a senior

**Congrats, during a pandemic year, sorry about that but**

Yeah

**Glad you made it through. Ok this is cool, you first—did you first encounter that prerelease exam style in Bio200?**

Yeah that was my sophomore year and I thought it was like super novel and super cool cuz like, I know all the bio classes I've taken we've always done these like cool and novel like discussion based learning, and I've always liked that, I like that like teachers and professors are putting in effort like trying to help us learn better, like trying to help the material stick to us better instead of like better their teaching style, I guess those come hand in hand actually, Like teaching style and like the way it sticks to us um and I think the bio classes are the only classes that—or yeah, I think there's also psychology but like bio class are like one of the main departments that are actually trying to help us so I guess in that respect it's kindof like helpful too in its own way, like knowing that people are trying to get it to us, like do the big vast abyss of communication heheh

**Oh yes, that vast abyss, we try to cross it in different ways. Well it's good to hear that you feel helped, I think certainly a lot of instructors are trying harder to focus on students and so I'm glad that's working for you, it sounds like so it sounds like that prerelease exam is one of those differences that did work for you, and I, what I'm curious about because I've talked to a lot of other students in 118 who had never encountered a prerelease exam before, and it sounded like a lot of them were sort of confused about how to take advantage of it and so I'm curious about when you first encountered one was there a learning curve of letting it be useful?**

Yeah yeah I remember we were so confused, like I remember the piazza had like billions of questions like "what is this" and like what is "redacted", um but yeah I think um I see that learning curve happening now, like all the people in bio 118 that I talked to they're like what the heck is this exam, this exam style is so different and I'm like, with my experience with this I was like Yo take advantage of this, this is amazing, and I tell them like just like study group, it's supposed to like—cuz biology, and here's another aspect I like about it because biology is like a communal kind of thing it's not a lone wolf standing at the edge of a cliff it's like, it's teamwork, like in research group you work together with other people, You're bouncing off ideas off of them right, and like in the healthcare field you are working with other healthcare professionals so this is kindof like applying all that, or helping us practice those skills that we need later, like collaboration, and being in that mindset of like we can help each other. So that's when I tell all my friends and they're like Oh that makes so much more sense, and I dunno, after that first quarter of Bio 200 I was able to embrace that, like, I think Ben told me that cuz I took another class with him I took bio 313, I had Ben for a lot of classes, I took Bio313 and he told me in retrospect that that was his mindset when he was putting that exam format into place and I thought that was really beautiful and yeah, I think like a lot of students—I feel like Ben should be more clear about this, like in class this is why I'm doing this. He's very transparent about everything but I feel that he should definitely emphasize more that it's about like collaboration and you know that's how like STEM like fields are and like healthcare fields and all that but yeah that's—but like back to your questions, it's um I feel like the learning curve was steep but once you know like what they have in mind, like what the purpose of it is, then it's like super beautiful so

**Cool. Um yeah I wanna poke at that a little bit more because it sounds like you have figured this out, you've made the system work for you and I just—I've heard from students who are having trouble with it so I want to get at a little more of like what is your process, now that you're used to a prerelease exam. So like for this exam that just happened, what did you do to prepare for it?**

Oh I um I don't know if they told you, there's a discord server, so with everything that's like online, the discord server is basically how we talk to each other---even though there's a Piazza I think discord is the thing nowadays, all the students are on it. But basically like someone will—there's this girl named Alyssa actually, she's really, she's very vocal about setting up study groups and I really appreciate her for that, so what she does is she's like I'm gonna set up a Zoom meeting at 3pm today, so just like hop on, but lately now she's been like ok I'm gonna set up a Zoom meeting, anyone who wants to come Has to come with like ideas of what they think this question's about and like more preparedness and more engagement, and so I go to those and we just bounce ideas off of each other and it's cool because some of us are taking it—well I'm taking it as a perquisite For my pharmacy school, and some other people are doing that as well—or not for pharmacy school but for other like school they want to go to that's in the healthcare field so we're coming with all these other experiences from like Other classes, other professions, and um even like nonSTEM majors, like they have a lot of experience with like in their day-today life just like hearing things, so all of us just come together and we have this rich study sessions and we're just hearing things from each other and learning things from each other, so yeah that's a study group thing, that's my approach I just do that.  And I look through notes again but that's-- I don't really do that as much as I like put more effort into my study groups, and that's like—I feel like I don't, I spend less times studying for this class but I feel like I do well studying in terms of this like class, in terms of all my other classes.  But just like a note, I am in my senior year, and I am already in pharmacy and I really just need a 2.7 to pass this class so I'm putting in low effort for that reason too hehe

**That's very resourceful of you!  Save your energy you can use it for other things, and congratulations for being in pharmacy school.  So tell me if I'm getting this right.  It sounds like basically ou get together with some other students in 118 and ideally they have diverse goals and have diverse backgrounds so they can all bring a different perspective to like what do you THINK is redacted, what do you THINK this question is. So you try to guess what the question is going to be?**

Yeah, Basically that's what we're doing.  We're trying to guess what the question is but in the process of guessing it, like some of the questions they're not completely redacted they're like, oooo I can't remember an example right now but they're like what would happen if redacted was broken and there'll be like three multiple choices and it'll be like the system will fail completely or it'll be normal or something like that.  And we'll kinda like go through the process and kinda break each step, like say Thrombin was not being produced, or like Factor 8 was not being produced or something, so then we would see like this is how it works, this is what would happen if you break it and we just go through each and then break it because this is really helpful because we're seeing all the possibilities of the question and we're studying at the same time because we're just reviewing all the steps.  Or we go through the multiple choices that are available, so like in the example question I was telling you so if the system stays normal, what would be broken and what would make the system stay normal kinda thing, kinda like that.  So it's not really like—oh what if it's this,--actually it is, it's like what if its this question but it's more like we're using that step, that question as a stepping stone for our studying, just like a study guide I guess. Yeah

**Yeah.  Like a study guide but it sounds like for you this is different than a study guide so I'm curious about like—I mean I guess I understand the tangible differences like one is a list of things to study, yeah?**

Yeah

**A study guide, And this is sort of a partial exam yeah?**

Yeah

**I guess have you done any exams where you do study from a study guide?**

Yeah there actually like plenty—like microbio I think, microbio, there was like a study guide that was like just like posted online and then I think uh were there, were there practice…it's been a while but I think like sometimes it's just a study guide and you just study that way um but sometimes like um how do I say this, the professor is like pretty clear cut about like what type of questions it'll be so writing in a practice exam, but study quest—like study guides I don't really pay that much attention to because really it's just telling me, review all the material again, um I don't really use that, I don't sit down with it in front of me I say oh cool that's a study guide, like scroll through for a few seconds like ok that's gonna be on there I should probably review that because I don't really know that, Um but I'm not gonna be like um this question and then I'm gonna—or like, I'm gonna say like, um, if there's a study guide for Bio118 like TPO I'm not gonna sit down and say like TPO is this, and here's a diagram of that, I don't use, I don't do that, I just go through the material again if that makes sense, did that make sense?

**Yeah, yeah absolutely. Um cool. I'm gonna circle back to you were talking---the process of studying from one of the prerelease exams, because it sounds like what you're doing is basically imagining different scenarios that could come from a process, and as you imagine these different things that could happen, you keep coming back to that process and like every step of the process and the fundamentals of it. Does that sound like I'm getting it right?**

Yeah yeah exactly.

**Ok**

Yeah yeah it's kina like in that sense it's the study guide sense of it, you can kinda understand like you go through that process and then we like study it um but like it's not like, it's like different than a study guide like you were saying

**Can you elaborate on how? I know it's kinda tough to pin down but**

Yeah um hmm.

**Like what does it feel like when you're studying from a study guide or when you're studying from a prerelease exam**

Mmmmm I definitely don't feel as alone as if I was just studying by myself, it's more like a group effort which I really like. Also like during the exam when I'm taking the exam after a prerelease exam like I, it's kindof exciting you know, it's like what is this redacted question and then Ben releases it by like while we're taking the exam and Ben releases it and it's like Ah Ben I got you there, or like ah darn we didn't cover this at all, sneaky Ben You knew we weren't gonna think about that and you just put that question in there, it's like you can kindof appreciate the exam in that respect, does that make sense

Yeah which is amazing because, from talking about how exams are like always traumatizing to being able to appreciate an exam sounds like for you this is really heping and that's fantastic! Um appreciating **the exam. Can you tell me more about that, like, um, what do you appreciate about it?**

Ok um I guess it's more like the journey, like the journey that it takes you through like um, or like the experience I dunno like with Ben's tests I never, I never feel like unprepared for it if I do like the study groups and all that I don't feel unprepared for it, and I appreciate that he like, I guess I appreciate just the fact that he prereleased it, made sure we weren't stressed for it and made it clear, like on the exams there are things like, for other exams I've had they never really show the process that they're trying to test us on, so like for example platelet production. Um so he showed a picture of the platelet production on there, so it seems like very deliberate that he's not trying to like test us on memorization but whether we actually understand it or not and that he was actually true to his like goal by putting that exam on there, you feel like I dunno like maybe back to what I was saying earlier, you feel like he's being very thoughtful about it, very like that he was thinking of US when he was writing that exam, so I feel like in that way I kinda appreciate that because not only is that different than like other exams I've had professors who are just like you know this is going to be a memorization question because kindof like not help us at all, it is kinda like uhh like a special teacher-student dynamic like relationship kinda thing that makes me like appreciate the exam. Um it's really hard to put into words, like I just appreciate the gesture, like it just seems so nice, you know?

**Yeah, you know like he's not just doing his teaching job but he is actually caring about his students**

Yeah yeah exactly. And I know that with all the classes I've taken Ben is like very thoughtful about his students, I just feel like compared to all the other teachers I've had like Ben is probably like my most favorite one because he's very student-centric, it's not like um I had a professor in ochem who he just teaches and has his office hours and then and then like, it's just basically like a standard teacher thing, like teach test teach test, and it's like there's like a wall between it, but when you feel like you're connected to a teacher you feel like the words the teacher says like resounds with you more, or it sticks with you more, like I can remember things that he said in Bio200 um like because it's just like his like humor, his like like the relationships like he has with his students, it's just that connection it's just more established? Yeah that's the word for it yeah I guess ueah. I guess this is like Ben is super cool as a teacher, maybe that's like kinda taking you aside, or taking me on a tangent from like actual exam practice.

**No this is really useful, I think—I know that one of the things we're hoping that comes across for students is that when they see the—kind like what you were talking about of seeing like what the teachers goasl are for the exams in relation to the students, hopefully students seeing that will make them feel like they're—as you say, less alone in the process**

Yeah, And makes it less traumatizing, because like you're suffering together, you're suffering haha

**Haha Do you feel like—so I'm gonna circle back to something you talked about, you talked about how he would—you weren't sure about HOW you did on this exam because sometimes he goes back and changes things based on how the answers work out. How does that like is that stressful waiting for it?**

Mmmm well ok, my answer is No but there are many factors that go into it I guess. Like first of all I know like Ben's doing his best, we're probably gonna get more points anyways so that just makes me feel happier. But also no the point is that I am already in pharmacy school, I don't really um it doesn't really matter to me! Um but I guess like putting myself into the shoes of other students, um I guess it could be annoying that you don't know your score right away, but it's um like with like the experience of the test itself, or the like knowing Ben as a person like having that kind of connection like I feel like I wouldn't be too stressed out like knowing that he's doing his best to like have a fair exam um and like like right now it's to give us more points, so I'm like fine with that haha, that's totally fine with me

**And it sounds like in a way you, your uh, it causes you to think about the process of making a fair exam, and like maybe like you've gotten some insight into the process of like I guess**

Yeah

**How the exam is made fair**

Yeah, you can kinda be like more empathetic because like he like explicitly says like exams are horrible at gauging your progress but they're what we have, and he makes an effort to revise exams, he releases it ahead so we can look through it and see if some wording is weird and so he's like obviously making an effort so we can't, we can't really—it's like really hard to hate on him as a professor because he's doing all the things that we want him to do, he's like hearing us basically, he's like I hear you, like exams are hard and so I'm gonna, like you're gonna help me with it, um I think that's like like it makes it you can't really be like stressed or annoyed about it, like the waiting process in my opinion, I don't know what other students have said but that's my opinion like having taken classes with Ben before like a few times so

**And it sounds like he explicitly asks you like (cough) excuse me, he explicitly askes you like help me get this right, is that? Yeah ok and how does that feel**

I dunno I just feel really really like happy compared to all my other classes the professor is just like you don't like the exam then deal with it haha, don't take my class or something like that. But Ben is like let's make this a team effort kinda thing. I dunno it just feels, I feel more respected as a student like kinda thing, it's not like—like for the other classes it feels like, or sometimes it feels like the professors are just there to get paid, do their research stuff like adverstise their research, it's more than, It seems like other classes ists professor-centric kinda thing, um like but for like Ben's classes seems like it's like a community sort of, like I dunno, if that makes sense

**Yeah definitely, that's, well that's great, I mean that's great to hear because that sounds like really valuable and hopefully effective too**

Yeah

**Which I wanna aske you about now, so you are a senior, you're into pharmacy school, again congrats, like you've got your pathway going so I would hope that this class in particular would feel really helpful and really relevant. Has that been the case?**

Yeah um I've I'm seeing like stuff that I have learned about in my other bio classes, in like my microbio classes, even my ochem classes, I feel like it's a nice way to end up to wrap my senior year just kinda going through, actually like even like seeing like bio 119, 118 aren't all stem majors and so they're like, I can see like growth in the people like work with who aren't like bio majors, who haven't had that exposure previously, and it's kindof nice because it looks like I'm just looking back on my bio adventures as a bio major and seeing like I was there, I was there in their shoes like learning this for the first time, like most of the stuff here I'm seeing for the second time, but it's kinda nice to have it all wrapped up at the end for me. And I feel like looking at all the inside look videos that he posts I feel like is really valuable as alike a sendoff gift into pharmacy school, I dunno, I feel like it's like taking all the important bits from my entire undergrad and putting it with a nice little bow, haha at the end of the line. But yeah I do feel like it's really valuable even though I've learned all the stuff before.

**All right, well it's good that you feel it's valuable but I'm gonna push you on that. Since you've already encountered all or most of that content once before, what specifically is valuable about it?**

Hmmm that's a good question. Um. I think it's valuable because in pharm—so I'm a pharmacy assistant right now, and a lot of stuff that we're learning like he goes over and he kinda even like puts the drugs on it like, what was it, cyclobenzaprine, or like some like drug that I've that I've sold at, as my pharmacy assistant job. Um I think it's just like it's valuable because he's going through all these steps, and going through all the ways that you can break it and all the ways that you can save it kinda thing that you can fix it. And he sometimes puts drugs in there and I'm like wow that's up my alley, I'm going to need all that stuff. So In that way it's valuable because it kinda segues into pharmacy school

**Like real world examples get used**

Yeah exactly! And I'm selling it at the store too, and I'm like this is a real thing, like this is cool I like that it kinda clicks with me it's a good headstart into pharmacy school because I'm going to start my pharmacy internship at my like job at Safeway now, safeway pharmacy and they're going to make me memorize like 200 drugs and so Bens class is a good way to remember like all these things and, like it's valuable I guess another way it's valuable is it takes all the big bits and summarizes it for me cuz you know like people like I always forget, like the reason why I'm not like acing every exam right now is that I always forget, there's so much I biology I'm always forgetting all this stuff and he just like reviews it for me, it feels nice, To be able to have that again, cua I'm always like insecure, like what if I forget my entire undergrad, kinda thing, cuz there's so much, and it's been a while too. So he kinda just sums it up, and I dunno I feel like it's nice to have in the back pocket just to have that overview again, um

**Yeah Well good. In terms of like remembering the stuff from your undergrad once you're a pharmacist—is that, you want to be a pharmacist specifically is that**

Yeah

**Ok great. So once you're a pharmacist and you're keepingup with the latest drug releases and trying to sort of solve problems and think about drug interactions and All these complex scenarios that you're ogin to have to deal with, um is—it sounds like you have encountered some ways of like, this is how I remember something. Is that something that you've taken from something that was done in class, or like something you learned on your own, or something you're going to keep doing, something you're going to change…that was a complicated question sorry I'll give you**

Yeah what do you mean by that

**Yeah I guess I'm wondering when you think about the job that you want to do and you think about the skills that you need for that job, has your college career and any specific classes, like hopefully 118 have they helped you prepare for what you'll actually be doing on the job, not just the content but the skills**

The skills, well yeah, like what I really like about the exams is like the social aspect of it, and pharmacy is very social it's very community centric and the work I do right now as a pharmacy assistant at safeway it's very collaborative, You're talking to doctors all the time and insurance companies and um yeah you have to talk to a lot of people and kinda like put the pieces together kinda thing, and literally that's like Ben's exam format, you're piecing things together like this person thinks that this puzzle piece is actually this, and then you think it's actually this, and then you put it together, and then You kinda have this question, this awesome study session I think it's like the skills, yes, that's something that I can take away from like any of Ben's classes, all his classes have been collaborative, bio 200 bio313 and this class, which is why I'm like a huge fan of Ben's classes but um I guess like other skills, like studying skills are you talking about or like what other skills do you mean?

**I guess I'm curious about what skills you think that you'll need at your job, and where those are going to come from, where you think you'll learn them. Does that make sense**

Yeah, I guess yea, skills, when I think of skills I think of it less as knowledge and more as like ability to like get information kinda thing, um, I don't think of it as like I can memorize all the Krebs cycle stuff, or um I can like list all these things off of the top of my head, I think its more like knowing who to talk to, like at least in my job right now you gotta know who you wanna talk to in terms of getting prescriptions for patients, or like talking to the doctor to get some sort of like change in a patients' medication, um yeah I, I think like the skills are like communication skills, like the collaborative skills, are the skills that I will take away, in like my entire bio classes like that has been a theme, like, like collaboration is the way to go kinda thing. Haha but yeah I don't know like what skills you're looking for but for me it's like collaboration

**Yeah that was exactly what I was looking for, actually I have a couple chronic illnesses so I love my pharmacists so it makes me very happy to hear you say that because you know the communication between all the different pieces are so frustrating**

Oh yeah

**So I really, it's very, I'm so happy that you know students like you are going into pharmacy, it give me a lot of optimism, I love my job because I get to talk to the next generation and you're all awesome!**

Yeah I appreciate that

**Yeah. So ok. Let me see if I got this right. In terms of like of some of the things you take from the class, so it sounds like you're presented with a whole bunch of different problems when you get the prerelease exam, like puzzles and you work those out as a team, where you all offer different things, You have different perspectives because you've learned different stuff, you've mastered diferet things. You work together to communicate and piece—and think about ok, this is a possibility that someone brings up, how would this then work?**

Yeah

**And then someone else brings up one, and how would this then work**

Yeah, and then we just go through like the system, like you know if we're learning about like this certain step, like step 1 step 2 step 3 kinda thing, like we're learning about that kinda system that just goes through it and like break it

**Go through it and break it**

Yeah, which I guess is like the essence of physiology, like that's good foundation of like physiology class, like, right at the beginning of a pandemic, and the professor also a pretty good professor, she told my like physiology is basically like learn a system, break it at any step—because that's what humans tend to do we just tend to break the prcesses and that's why it becomes issues. And um then you figure out what happens afterwards, how do you fix that and as a pharmacist that's going to be important too, like which step broke kinda thing and which step needs fixing. So yeah this is why physiology is pretty important

**Huh, this is very cool, all right. Thank you so much for talking with me, this has been really valuable, really insightful to talk to a student who has figured out how to use the tools that you're being offered. And so I wanna ask, you talked about watching the learning curve of some of your peers, trying to figure out the system and I wonder how could we as instructors help them or you know, help**

**yourself back in the early days, to figure out how to use a tool like the prerelease exam um effectively and in a way that will help them to learn long term skills for their jobs.**

Ok yeah First of all I really like the inside looks, I fell like that's just really good for getting more people to jump into healthcare especially if they weren't even considering it because even as an incoming pharmacist myself I took some insight away from that, there was an inside look for like emergency like emergency practices this emergency doctor she was like, or they I don't really know like the pronouns, but they were like I—when I'm off of work, I'm off of work and I don't have to do anything and they were calling from some vacation spot which was like really cool. So yeah even for me, someone whose already figured out kinda what I want to do in life, those were really helpful. And even like I'm thinking now after watching that video About like emergency pharmacy, which is kinda cool but I don't know anybody in that spot ss I'm just gonna like go talk around but I really appreciate those. In terms of helping with like the learning curve I feel like talking to students who've already worked with Ben, like some of the students had already worked with Ben and the fact that they're coming back shows that, the fact that they're still in contact with him even, shows that he's a really good, like he's really effective at keeping student-teacher relations beyond the classroom, Which is really powerful, like when you're a student, like a current student and like looking to him as your professor, like it really helps with student dynamics. Um I think maybe getting those students to talk about how you can be successful in Bio118 or in Ben's style classes um like how you can use all these formats, Im pretty sure like you can like tell them to like come back and have some sort of like panel, like during class or something, like this is how you can be successful, like make use of these tools. And I'm pretty sure that like even if I had that panel that kinda resource for me back when I was in Bio 1—or Bio200 when I first had Ben, I probably wouldn't be able to understand that full, like kinda the knowledge I had now if I tried to pass that on to like my younger self I probably wouldn't be able to understand it fully but I would have like that seedling growing, like Should I do this instead of like you know, studying and trying to memeorize all those steps, like study, I'll start taking the steps that I need to get here, like get to the place where I can kinda understand and like fully take advantage of the tools that are available to me. But yeah, I just feel like jus knowledge and insight, enlightenment from like the older classes is cool, um in terms of making use of all that and appreciating it, cuz I feel like I'm still in contact with a lot of friends who like took his classes and they like, he's probably one of their profe—like favorite professors too and they know and appreciate his teaching style um but yeah.. Um I just feel like kinda like knowledge and wisdom from older like the people that have passed through kinda thing. Cool yeah.

**All right I'm gonna do something a little mean which is I'm gonna put you on the spot, ok, you're on the panel, it's you, what are you gonna tell them**

Oh ok, Well I'm gonna tell them what I already told you, I guess. Like this class is not supposed to be like professor looking down upon his students kinda look, it's more like a professor helping his students so you think about it, um yeah I don't like being put on the spot

**Sorry**

No it's good, you think about it more as the professor is there to help you which is unlike all the other clas—or actually which is not shared by a lot of other professors in other classes, I'm not gonna diss all the professors. Um you'd think about it more as he's trying to make the experience much more lighter on you, much more helpful for you and look for those places where he's trying to help you, then you will learn to appreciate this like everything more, but I guess like as a student in this class just appreciating the fact that he's trying to be there for you is not enough as a student but if you take the advice that he gives and just follow it through just, source, trust me on this, um, it will help, um, and a lot of skills that he will teach you in this class will be helpful in in whatever profession you want to go to in the future because you learn a lot about collaborating, um together um in like while working together for those study groups, and you learn about—you learn a lot about physiology so you can dispense all the false media, false news, fake news out there. Um but yeah I'm—I feel like I am horrible at summing up exactly what I told you like if I'm being told um just give me advice

**No that was great**

Ok

**No that was a really good summary you know collaboration, realy world thins and take the Take the advice of someone whose trying to help you.**

Yeah I was just agreeing with you

**Ok um I'm gonna press you again just a little bit, you said take his advice where he's trying to help you, what kind of advice does he give?**

I guess he's like very transparent about his teaching process, oh yeah that's another thing I would say, he's very—his—he's very transparent bout his teaching processes and he says I am giving you this so you can do this kinda thing, like he's giving, well I guess he hasn't been very I dunno if he's been very transparent about his, about the reasons why he does this style, but like if he says like "I am like being transparent about this, I am giving you the exam ahead of time so you can like discuss it" oh wait I think he did say that once in class um but yeah, you should go and discuss it kinda thing. Um or he's saying, I'm giving you this exam not so you can be, or not exam, I'm giving you this diagram because I don't want you to be scared by it, it's a very mean looking one, but I want you to understand it, like I remember this one time he said in class, um like he gave this really complicated exam that had a whole bunch of like drugs on it and I really liked that diagram and he was just saying this diagram is just to show the many ways you can break this system then fix it and I was like I really like this diagram Ben because all these drugs are on it and how to fix it, And these are drugs that I see like in my job as like a pharmacy assistant

**It was not just like oh let's theoretically learn this thing, it was like learn this thing using real examples**

Yeah but he keep saying like don't memorize this but understand this, and in like understanding it I kinda just memorized it, like that's his advice to me, like don't be scared just understand it

**So you're saying that the process of understanding it led you to memorize it?**

Yeah, just accidentally like not on purpose but like Because I understand the diagram like I understand all the processes that like that follow and then I can just like associate names with it and memorize it and he's, he said don't memorize it but I accidently did because he said don't memorize it which is like, which is what I really like the kinda like low stress if you say memorize this people will memorize it and forget it, but you say understand this people will understand it and memorize it just accidentally which yeah I think is cool

**Yeah that's awesome.  Well I uh we've been talking for almost an hour and I want to just thank you again, this has been so useful for me and really as I say very inspiring to think that students like you are going to go be in the real world making changes and working, so that's awesome.  Gonna kind wrap it up just asking you, is there anything else that like I should ask other students about, or something else that I didn't ask you about that you want to talk about**

Um not particularly I feel like I think like I kinda went through like my like my adoring of Ben's style very thoroughly haha and yeah it's kinda nice to tell people about it I just really like his style.  I think these questions were cool

**Ok great, and then the last thing I'll tell you about is your Starbucks card blah blah…**

### ***Interview #7 May 12 2021 @CC site***

### ***(Leah Lily interviewer = bold, with 3 students S, T and [redacted because 17yr old])***

L: Any questions before I move on?  Ok

[]

L: Ok Great Let's see, and I do need to check and make sure that, are you all over 18?

S: I am

[]

L: Oh, um ok.

T: Yes I'm over 18

L: Ok thank you

[]

L: Uh your video is frozen but your audio is ok

[]

L: Yeah so you're 17, that's fine, you can participate and your input is going to help steer our conversation as a group but I just won't be able to use your data on the research. Does that make sense

[]

L: Um T, I think let's see, I think I'm getting some echo from you, um from your audio there, um so I don't know I hope you all can hear me ok, and it's not too, it's a little confusing for me so I'm just going to try my best and T I don't want you to feel like you have to stay on mute all the time so we'll just kinda work with the technology we are given here and yeah.  All right, Well.  To start out with what I'm curious about is, how is Bio 213 treating you, how is it going?

[]

L: Yeah, thanks.

S: I would say to describe like the best way to say it is like this class is at least for me been very memorization based.  Um Feel like there's a lot of things that I've had to know just off the top of my head for other quizzes or especially this last exam that we took.  Um. Not necessarily like greater concepts but kindof specific cases, if that makes sense.  And definitely feels in this case like the expectation is that we are on point with reading the textbook and getting as much as we can from that information because we are held to that like pretty much all the information that is present there, which I believe is fair but which is a little more difficult in kindof an asynchronous environment

L: Yeah for sure

[]

L: Yeah T

T: Oh sorry yeah, sorry I was driving and I had it on mute but I guess I have the same sort of experience as they did, um, I think coming into this quarter I kindof expected it to be a bit different than the last class that I took for 212, um I think that was more lab-based and I guess I should've kindof expected it to be this quarter to be a bit more lab based. I feel like a lot of it self-t—like I'm teaching myself a lot of the material based off the study guides and stuff like that and Its' kindof made it harder to understand without talking directly to Dr. Gwen, well the professor, about what 's going on in the chapter

L: Yeah yeah I'm curious about that like, so you feel like you're teaching yourself, what kind of materials do you feel like you've been given to help with that cuz it also sounds S like you have to go through the reading on your own and master that on your own as well so it feels like a bit of a similarity there.

S: Yeah yeah and I would definitely agree with what T was saying in saying like we're given kinda each week we have a chapter that we're going over and we have usually a video lecture made by the professor to go over which is covering the content of the chapter as its kinda presented in the textbook. And so between that and the lecture that's kinda the study material we have for the week's content, I would say so it—at times it's um a little tough to um look through those two and then get all the information you need and then at the end kinda when we're going to take kindof what we call the post lecture quiz we're given a study guide which is kindof a, the content the professor wants us to kinda hold on to the most, um, and that's kinda like at least in my experience what I've used the most in kindof just to study what I need to be learning, is that study guide.

L: And T is that, that's the study guide you were referring to as well?

T: Yeah yeah

L: So you feel like you're teaching yourself from that—how does that look like when you're teaching yourself?

[]

L: Oh haha yeah sorry that was, I worded that really confusingly, sorry about that. Um ok um what so I guess my question rephrased less confusingly would be, um when you are trying to teach yourself that material, what do you do?

T: Well at least for me when using the study guide I kinda just read through the study guide entirely and then read through the chapter and try to reference back and forth between what sort of questions she's asking with the study guide and what is talked about in that section for the chapter and sort of try and understand what exactly the point is that they're trying to come across um and then when taking notes for like I usually will take notes directly onto the study guide, just to write what my understanding of the answer is or my understanding of what she means in regards to our understanding of what she's looking for

L: Ok

[]

S: I would agree as well, yeah, I would say the main focus this quarter has been our group lab project as opposed to kinda our course work and kinda content that we would be getting on our exams

[]

L: Yeah and you um you mentioned you did want to talk about exams and that's great, I would love to hear about exams Especially because it sounds like, correct me if I'm wrong but it sounds like the exams are a lot of your grade but it feels like the course is a lot of the lab which is not very connected to the exams

[]

L: Yeah yeah totally, how—and ok so a couple things I wanna focus in on, one—and S and T I do want to hear what you think about his ask well, you said she makes the exam harder and I wanna know I guess like how? And I do wanna hear more about that what did you call

[]

L: Yeah the preexam as well, just how that works for you.

[]

T: No no I was just gonna talk about the structure of the exam, I think it started I don't know if it started before in regards to like 212 because we were taking our exams via zoom she kinda—the setup was not like the, the normal exam where it's like you get a bunch of questions and then some answers and then you choose an answer, it's more like essay based pretty much, so you get a compare and contrast essay as the first question and then she has a few other, like the phylogeny question where you answer you know different parts of the tree and your understanding of it and you have to Find where something is on there, and then a graphs sort of question and then the last one is like the drawing so it's kinda hard to like look for the answer because you have to know everything off the top of your head to complete the exam pretty much.

[]

L: Um yeah S does that resonate with you

S: Yeah I apologize I'm trying to gather my thoughts on it

L: Oh yeah take your time

S: You know like the best way I could describe like the questions we were given on the exam is that they're very specific.  Um and it's kinda like what I was describing a little earlier on is that the questions that were asked both on the exam and on the preexam that we're given is they require quite a bit of kinda memorization of specific things and many times especially kinda on the—I would say this happens most commonly on the compare contrast essay—is that it's, it's a very specific um set of things that she wants us to talk about and sometimes I don't have as great um memorization of so I I think that would be the best way I could describe the exam, is like very specific

[]

L: How did, um for the yeah for you T did it is that very specific knowledge and that memorization does reflect your experience?

T: Yeah yeah I would say it was definitely the same I think with the preexam I've noticed from previous exams she will normally pull, or at least for the one that we just took, the two questions on there were, one of the questions was definitely one that she put on the exam so I was kinda expecting to see uh primarily the compare and contrast question, I was kinda expecting to see that So I know that my focus was on like understanding what exactly the question—I think in the preexam she took out, there were parts that were taken out so you kinda had to guess what exactly would be compared but yeah I would say like a lot of it would be memory based and things like that

L: How did you decide what you were going to focus your studying on?

T: Um I think I just referred back to a lot of the study guides I used before and kindof set those aside and created my own sort of sheet that compared and contrast whatever was on each study guide to understand how those releate and how they're different Just in case I needed to use it for the exam

L: So it sounds like so it sounds like you used mostly the study guides that you'd been given throughout the course rather than the preexam

Mmhmm well yeah I would reference the preexam for like the structure of the –cuz like I already sorta knew what the setup for the exam was gonna be, I knew there was gonna be a compare and contrast, there was gonna be a phylogeny question and interpreting graphs question and then a drawing question so, a lot of my focus at least was on--

L: Hello? Oh you're frozen, oh no that seemed really key haha well I hope she'll come back oh no, um

[]

Well oh I hope we get her back, I guess while we're waiting to you two how did you decide what to study for the exam, those specific things that you had to learn.

[]

L: So it sounds like you didn't have a particular focus based on anything you were just like I will study everything

[]

L: So you're saying—T we got you back!

T: Sorry my phone overheated and it shut off for some reason but I'm here now

L: Oh no, yeah no worries, you were telling us what you, how you chose what to focus on

T: Oh yeah So yeah I based it basically off the preexam sheet because I already had a history with the previous exams in 212 so I kinda knew what the setup would be and how the—in 212 we went over how to write the essay portion and what she expected from us so I sort of based it off that knowledge that I had from the previous quarter and then already knowing that there would e those type of questions on there I didn't feel like I needed to understand the interpreting the graphs portion because I kinda thought that that would be—for me at least, it was  little easier to, I think that was the easiest part of the exam so understanding the graphs, I think the drawing and the compare and contrasts were probably, has historically been the hardest part for me yeah

L: I mean those are the, those are hard, that is totally legit. And had you, so, had you had you had practice of interpreting graphs of 212 or your previous classes

T: Um I think we did a bit of it for the lab, I'm not sure and I know we did a bit as far as like the research project if I remember correctly, um she wanted us to have that, or we were supposed to have that on a few, depending on your research project you had to have an understanding of creating a graph for your research project so that was something that my group sort of talked to her about, In regards to how we would formulate that so there was yeah, there was a bit of background in that one.

L: Mmhmm and um it sounds like you said you pretty much knew what the structure of the exam was gonna be like, and that it sounds like was more from you taking 212 with that same instructor, yeah? And not so much the study materials that you were given but that you'd had Dr. Gwen before

T: Yeah yeah

L: Thanks good to know. And S did we, I think we didn't get to you yet

S: Ok, yeah I, I mean I do want to freely admit like this past exam not sure if I'm a great data point for, I was going through a lot of events outside of class that compromised my studying especially for this one but I can kinda talk about my experience going through 212 previously cuz that's kinda what the same kinda format for this, and as I was thinking about it I definitely agree that like this exam definitely kindof is more…more understandable having taken 212 previously because there are certain things that I can completely understand []'s kinda fish out of water feeling with it. Um. But definitely what I normally do with studying for exam like these is the first step is taking a look at the preexam and kinda what, what the preexam is offering because she'll give us kinda options that we need to be aware of the questions, like we said before, and from there I kinda see what kind of focus areas I need to be looking at in the textbook for studying, just getting familiarity with specific cases of what this compare and contrast might ask for, and I defiantly find myself paying attention more to kina like more likely the exceptions to the rule, often times if that makes sense, kinda the corner cases, the more irregular occurrences because I found like more often than not those seem to pop up a little more kindof what we're asked of

L: T I see you nodding about that so it sounds like your previous experience just in school has taught you that on exams you usually get asked about the exceptions is that right?

S: Yeah

T: Yeah

L: Yeah nods, ok great, that sounds, that sounds frustrating an it sounds like my experience as well you know they're gonna ask abut the hard one hehe

S: Hehe

L: And I do I also wanna just affirm, S, it's it's um we are looking for a wide range of students to talk to and it's very much a common thing for students to be going through other stuff in their lives so it's important that you know we as researchers know what works for students who are like really busy and can't study as much as they want to , what helped you succeed anyway so you know I really appreciate you sharing that information because that helps me think, like when students are struggling what 's a helpful thing

S: Ok ok that's a good thing to think about, I didn't really think about it in that way

L: It's hard to be a student you have a lot going on. So I'm glad we're talking about exams and that that was something that came up for you all because the specific thing that we're actually trying for to change in this course to look at is the pre, that preexam. And try to use that to basically what you're talking about which is, Does this help you get a better understanding of how to prepare yourself for the exam. And it sounds like yeah, pretty much, But it also sounds like there's some drawbacks and so I'm curious about—and limits, as well, so I'm just curious about like how do you think we could improve that preexam so that it would solve some of the problems of like, for you [] for example like you should learn how to study a graph and you're gonna have to know that because you saw on the preexam, as an example.

[]

L: And T and S it looks like you were kinda agreeing about that, S you specifically talked about –I'm going to say specific again, the specific memorization, And T you talked about going over like the specifics of the study guide so does that feel useful, does that feel relevant um

T: I think it really depends up at least for me it, I felt that it helped a bit but trying to memorize a lot of the material in such a short amount of time and basically trying to retrieve that material from your brain during the exam is hard for me and I feel like if we had a better understanding Of each part of the the what we're learning conceptually as [] said it would make it a bit easier for us to take the exam and actually know what we're being tested on and understand the material a lot better

L: So like not just memorizing, but being able to use what you already memorized.

T: Yeah yeah

L: Yeah that yeah that makes sense. And I think um yeah there's defiantly a lot of memorization in biology or in like any subject but definitely in biology, but it's also very important that you know what to do once you've memorized something so do you feel like you're getting that knowledge maybe from lab or another part of the class

T: Um I think most I don't know for S and [] but I feel like most of us are getting that from actually having to read or watch the short videos that she sends out or actually retrieving the information from outside sources like youtube videos and stuff like that so not necessarily--I believe last quarter we did sort of a review at the end of what we felt should change at the class and I think a lot of people brought up that we wish there were more lecture based class—er time in zoom where we just go over the material with the professor and get to ask questions and kindof understand what the point of each chapter and what we should know and things like that

L: Mmhhmm and those are the synchronous sessions right?

T: Uh yes I believe so

L: So and then the youtube videos and the research you did was that stuff that was assigned or stuff that you had to go find on your own

T: Um some of it we had to go find some of it she would post for each week but if you still didn't understand exactly what you were watching or reading you would have to of course go and look it up and see if there were an explanation somewhere outside of the material that was given to you

L: So it sounds like we're kinda circling back to what you all were saying at the beginning which is that you kinda have to teach yourself a lot the material, yeah, seeing some agreement, that sounds sort of frustrating and I wonder if you have ideas about what would help you at least feel supported in doing that kind of work. Because that's certainly a useful skill, and something I've had to do as part of my research is finding new information but I also think it's also a skill particularly if you're sort of new to college or your university career that needs support.

[]

L: More specific study guide questions, so like you'd know exactly what to study?

[]

L: Did the preexam help with that at all, like give you an example of "here's how we're gonna ask about your knowledge"? And then you can figure out from that? It sounds like you did a lot of work with like, some of you if not all of you started with the preexam and worked it into your studying, did that help at all to know how to answer a question and how to concentrate your studying?

S: I would say yes, um that is kinda the…if I think I would say like that's where at least for me the true studying really began, it's not in kinda the week by week chapter work but it's actually kinda preparing for the exam where I get the most studying in and I was kinda, I was turning this over in my head and one of the things I thought was the best kinda going through this past year and covid and turning into a more asynchronous learning, in my organic chemisty class kinda each week we were given kinda a worksheet to print out and kinda work through as we were going through each chapter and it kinda covered a lot of the concepts, we were writing in the big ideas, Performing kinda processes that they were teaching in that chapter, and at least for me that was very beneficial, I feel like if we were given that as kina an aid week over week I think that would give us kinda a better idea of like these are the concepts that we should be turning over in our heads, these are the concepts that we need to be paying attention to, I would say like outside of that how we're going about it sort of right now is Kindof going through everything kindof trying to figure out like what information is more important, which what information is a little less important and I don't know if [] or T you guys kinda feel the same way or

[]

L: Understand it enough to be able to apply it, and it sounds like one thing that the chemistry worksheet did was like it walked you through the processes that you have to understand in order to apply the thing you're memorizing

S: Yeah that was at least in my mind it was kinda a little more structured reading in that case than like ok I'm paying attention to these things as I'm reading through the textbook at least for me that kind of additional structure is really beneficial to kinda my studying and my retention of a lot of the key concepts.

[]

L: Yeah yeah more conceptual, more about the process that's happening rather than having to just memorize

[]

L: I'm seeing nods across the board, yeah tell me a little about what walking through the –I guess, more structure more guiding more walking through the what's happening

T: Yeah yeah I think that would, I would defiantly agree that that would be a lot more beneficial to the understanding of the material all together. At times I kinda feel that we are going through everything kinda just to have some sort of knowledge of it but not necessarily understand how they relate completely because they don't have that set time to kinda sit together and discuss it. I think she has like lab gabs or something like that, I've never been able to attend it because I'm usually working during the time, I'm not sure if that's usually when she goes through the material and explains it further, I'm not sure if you guys have attended any of those

[]

L: Yeah that's definitely an issue. I'm curious, I you mentioned something about like studying in a group T, do you do that, do you use a study group?

T: No I did last quarter but I haven't been able to do it for this class specifically, I feel like adding on a study group last quarter had really helped to me to understand the material a lot better cuz then I was able to sort of shoot my ideas back and forth with someone else and see if my understanding sort of matched theirs and what exactly I was missing but I think this quarter it's a bit harder to do that, or for me it's a bit harder because I'm working and on top of that I'm taking another chemistry class so a lot of my focus is between the two and trying to understand the material in both classes yeah

L: Yeah um is that [] would that be kindof an option for like you know if you can't actually ask your question to the professor um you know can have you tried asking your peers, does that work the same way or

[]

Gotcha, yeah that makes sense. Um I'm conscious of our time, I don't want to keep you here too long because I know you're busy, so, anything any thoughts just to wrap up with or um either about the course or about like questions I should be asking your classmates about how to understand this course better?

S: I mean I think one thing that I definitely want to mention is kindof at least coming from 212 where we were taking all of our exams online we took our exam in person socially distanced um this quarter and at least in my mind it was a marked improvement and I think that's also just a consequence of like getting back in person, having more actual in person interaction with the professor, peers, all of that. Definitely, definitely helps. I um honestly can't say that enough, I think at least for me I feel a lot more discombobulated when I'm interacting with everybody through a screen.

L: Yeah I hear ya. Hah I think your classmates hear you too

[]

T: Yeah definitely agree with [] that I would hope that in the future when we take more in person exams that the structure will kinda change to sort of match our understanding of the material, or at least the way that we're sorta studying it, because we're kinda just studying it all together to sorta memorize and understand all of it but we're Only getting tested on certain aspects of it so it's kinda like are we wasting our time with studying everything, could there be a point to where we are being testing on what exactly that we're studying or memorizing

L: Mmhmm feels like a lot of work that may or may not actually be relevant to the exam for sure and my I guess my pushback question about that is since you're taking the course for a reason, um, like is it useful to have motivation to actually study everything in the course even if you're not tested on everything?

[]

L: Yeah.  That makes sense. Ok well I will wrap it up, um, so your starbucks…

### ***Interview #8 May 12 2021 @CC site (Leah Lily interviewer = bold, with students M, N, P, and Q)***

*Note that audio-recorded voices for M and Q are occasionally difficult to tell apart*

L: …any questions or concerns for me before I jump into our conversation

All: Uh no

L: Ok cool let's see here did I cover everything, oh yes, your consent forms, I do need those in order to get your data included in the research, if you're not ok with consenting that's totally fine and you can still participate in the conversation and what I'll do is I'll basically … is anyone under 18 here?

All: No

L: Great, lucky me, cuz that's hard and sometimes it's hard to be like oh wow you just made a really good point but I can't use you, so … one last call for questions before I give you the first official question. All right well so what I'm wondering just to start off with is how is bio 213 treating you?

Q: Um I guess I'll go um so far it's ok I just feel like since we meet in person at least once a week that's so much better than bio 212 was just because it's like I'm more of a hands on learner so like going in person is really helpful yeah so I honestly like it better than last quarter just because it's more hands on than last quarter was so

L: Yeah that's good, I'm glad covid seems to be hopefully winding down

Q: Yeah

N: Yeah I think going in person is also like it's very helpful because I can like I can ask the teacher questions in person and for clarification, it's a lot like faster understanding when you don't understand something and you can just like have a one on one with her like on the farm

P: Yeah and for me like I feel It's easier than last quarter cuz like because we have research that we have to do it at the end of the quarter so we can meet our group member so it's easier

All: Mmhmm

N: I'm just glad we can meet in person at least once a week, any other subject I'm ok with doing it online but anything science related it's better to do a hands on in person cuz just some of it you can't try to teach yourself it's a bit too difficult so I'm happy we can at least meet once a week

L: Yeah good good yeah it's harder when you're—partly you don't have that sense of community as well and yeah, asking the professor for confirmation is always really important. Good well um I'm glad you're liking it better than last quarter, how is it going, like, is it hard is it easy, frustrating, fun

M: It it seems like it was uh it's pretty easy so far for me like I feel likehte exam was easier than it was made out to be like, I thought she was like memorize all these things cuz like tha'ts what I would have done then I did but then it was not those things, it was like easier things than the stuff we memorized, so I fee like it is definitely easier than it is appearing

Q: Um me personally for the exam uh mine was the exact oppostite of M's, I had the exact opposite problem I studied a bunch of things and then on the exam there's like stuff that I hadn't like thought about in like weeks so it was kind of like difficult for me personally, so I kinda was like stressed because of that but yes it was ok

N: It was the same for me too, it wasn't too easy but it was a little more on the difficult side there were definitely things on there that I hadn't thought about for a while or I thought I had down and once it was in front of me I was a little stressed out

P: Yeah for me I feel the same, it's less stressful than last quarter but in the middle, not perfect

L: Ok.  Um so I am curious about when you are studying for your exam, what do you do to prepare and what is helpful

M: Um for me I started like a week beforehand and I spend like a day on each chapter like going through my notes on the chapter and making sort of like a study guide with all the information so I had three days for the three chapters and Afterwords I used the study guie for like the remaining four days and like whent to the lab gabs and anked her questions about what to do so like memorizing the life cycles and I feel like that process is really helpful for like remembering all the older chapters Cuz sometimes it's hard when you have new information you forget old information

N: I used the active study recall, that's really helped me a lot in like STEM classes, I don't know if it helped me too much with this exam but it helped me a lot in the past

L: Active study recall, what is that

N: It's like as you go through the chapter you write your own study questions or even based off of the study guide and When you're done reading it you go through it whichever ones you know off the bat you put in green the ones you struggled with are in yellow and the ones you didn't get are in red so then So then you really know which parts you need to focus more on

L: Oh that's really cool that's an interesting system Where did you learn that

N: I learned it when online learning really first started cuz I didn't know how to study so I looked up on youtube best way to study online and that was just like the number one that kept coming up

L: Awesome P or Q how are you studying

Q: Personally I use stuff like quizlet or flash cards in gnereral, I feel like flashcards really help me, and I usually write stuff down so I can like I guess go into my brain in a way Just so like I fully understand it so I like rewrite all that stuff just so I can fully understand the concept of it

P: Uh huh for me I am just using the study guide

L: The ones that your prefeossor gives you?

P: Yeah mm hmm

L: Yeah are those, how do you use that?

P: Um cuz s it shows you what you have to focus on

L: Ok

P: So yeah I just read what I have to read yeah

M: Oh um for this study guide I thought it was interesting she did it differently so the last quarter it was like we could add our questions and like she might choose our questions but in this one she sort of like she put exactly what was going to be on the exam but with a fill in the blank kind of situation so it was like sometimes it was easier to prepare but Sometimes it was way harder because it was like compare and contrast blank taxa so that would be way more like vague so it’s Hard to prepare for but then there was like the other prompt would be like compare these two kinds of endosymbiosis so it was a very interesting way to study off of it felt a little bit more reliable

P: Mmmhmm

L: More reliable, tell me about that

M: Like um like we knew exact- like in some ways we knew exactly what we needed to understand and study cuz sometimes ther’s like so much information or like small facts that we thing we need to know but then in the big picture we don’t actually need to know it

L: Ah

M: And like it was a lot more clear I guess

L: Was that useful for the rest of you as well

Q: Yeah

Q: Yeah it was it was nice to know like what exactly we should like look for or like study for so I guess that was like the helfpful part of it so yeah

N: I feel like when I was in bio 211 we would have a study guide for each chapter but the questions were pretty vague especially something before a midterm or a final and even me, everyone in my discord group in 211 we were like there’s so many questions We don’t know which one to focus on, which one is gonna be on the exam, which one is not, so this was better because I knew what to focus on

Q: Yeah it was like the singular document, in 211 like she would just give us a link to every single chapter we did in the entire quarter and it was very hard to use

P: And there’s mini lectures, you can watch them or look at the back of the book there’s some questions you can use that summarize the Chapters

L: Uh huh

P: Uh uhu

L: It sounds like you didn’t get a practice exam, because it’s sounds like the questions were partial but you could do those practice problems in the back of the book

P: yea yeah kindof

L: Kindof?

P: Yeah yeah yeah that's right

L: Ok

P: Mm hmm

L: Ok and um Q I'm gonna circle back because you were like that was the good part of the prereleased part and it sounds like maybe there's some bad parts too

Q: Um ok so for 212 when I was in 212 she would like for example there's the compare and contrast she would put like 4 different questions so you knew What exactly it was but for this one it was like compare and contrast and then blanks so you were kinda I guess stressed because you didn't know exactly, so you had to study it like like all of it Overall and be like oh is it gonna be this or is it gonna be that so you kinda have to know the concept of everything which kinda stressed me out because there's a lot of information so I had to like study extra

M: It felt sorta necessary to me I dunno

L: Yeah I guess as a teacher my instinct tis to be like ha! It made you study more so it worked! But also if it was stressful that's bad

M: Yeah it was just, I feel like it was just too much information to like grasp so it I feel like it was too much information, I dunno that's what I think

N: Yeah like general taxa, If it was just plants and algae and like protists or fungi instead of like all taxa because it could have been any of those phylogonies which is like 60 options about and then compare and contrast any of those, it could be any of that it was really stressful, I'm glad I didn't get that prompt so it was ok

Q: And I understand the fact that we should like learn it all, like we should have the idea of it but at the same time its like a lot when there's like compare and contrast and graphs and like all that different kind of stuffs I just feel like it's too much for your brain, like I understand like Oh you need to know this and you need to know that, like I understand that we need to know it at the same time it's like a lot at once.

L: Um is there, Im kinda thinking like, it sounds like there's sort of two different things you have to study for, 1 you have to know the material because you're taking this class for the reason, and 2 you're taking an exam, and you have to pass the exam, and it sounds like that prerelease was useful for the exam, or more useful for like knowing everything.  Did that question make sense

N: I think it was useful for knowing everything in my case cuz I was personally less worried about exam taking but I know a lot of people like they're more focused on um preparing themselves mentally for an exam which can be very stressful

L: Mm like being calm?

N: But there's a lot of memorization

L: Sorry say that again

N: For some people I feel like, they feel like they gotta prepare for the exam more in a mental way and less for just the knowledge part

L: Oh gotcha

L: And what helps with preparing for the exam in a mental way?

N: Well um for um some people I know with test anxiety it was more like being absolutely sure you know all the material but it can be overwhelming because you never feel like you know all the stuff that you need to know but that's why I sorta go overboard with my studying and Try to like start really early because I want to make sure there's like no possibility that I'm not gonna know what's on there but you never know til It's in front of you.  But also like memorizing a lot of stuff for an exam is also it feels like It's more for the exam and less for knowing in general because you forget everything after you finish that exam, like memorizing nine different life cycles And now it's like gone

L: And like if you're not taking an exam on it you can just look it up

N: Yeah yeah

L: So so what I've been talking to other students too they're talking about like the memorization is for the exam but then what happens in the real world is you need to apply your knowledge and so I'm wondering is there something about this course that's been helping you sort of prepare for the real world, like apply that knowledge?

Q: Mmm kinda thinking about it I'm not really sure how I would—well for me personally I don't know how it's going to help me in like the real world um cuz the field I'm triyng to go into is not really about plants or like fungi or all that, it doesn't really apply to my like course that I want to go to so that's why I'm kinda like eh, but not yeah, you know

L: Sure sure are there oh go ahead

M: Oh I was gonna say It's the same thing for me I don't know right now I can think fo how it'll apply to anything I want to do in the future just cuz of the field I want to go into But it's definitely helping me build on what I need for the next course

P: Mm hmm yeah for me like I don't like to memorize things, I like to touch things, I like to work with the grove Or something I can understand more for the future

M: For me I feel like it is sort of what I want to go into because I want to go into biology wth like animals and plants But um I feel like going to the farm has helped me really understand the knowledge like in the real world because like she can point out and be like this is the sporophyte and this is the gametophyte and I'll actually be able to use the knowledge and comprehend it in real life So I feel like that's helpful for me at least

L: Mm hmm so it sounds like well this is really cool for me because it sounds like I have a range of student goals and since what we're trying to do is make these courses more useful and relevant to all students, well, what do you all what to do if you know already. If you don't know that's totally fine, there's no need to know yet trust me on this

Q: Um I'm personally trying to go into like dentristy, like that field, so I'm not sure how plant biology really like works with my future goals though

L: Yeah that's fair

Q: Yeah

N: I wanted to go into medicine to become a physician's assistant

L: Awesome

N: So that's why I'm not 100% sure like I can't say that plant biology is going to help me in my future goal

P: Mmhmm yeah I'm going to go to dentistry too

M: I'm going to go into Zooology so like exotic animals and so this is like what I

L: Cool cool well then I guess, hmm, one thing that we've been trying to do with how exams work is we tried to match them better to make you students feel like what they show on an exam is going to be a skill that they can take with them like a generalized way of thinking or a way of studying or keeping up on the latest in dentistry or what's happening in the field, in PA work or in Zoology or whatever. Have there been any of that sort of generalizable skill that you've been able to work on in this class

M: I feel like there was with the life cycles of plants that I was memorizing for the exam, I feel like that was really something I'll get to take with me because I actually understand it when I look out at plants outside now and I feel like that'll definitely help me understand like horticulture and like animal based biology

L: Oh we lost P, oh well, I hope she comes back. Well yeah, carry on

M: Oh that's ok that's mostly what I was trying to say

L: Yeah ok like you can look around you and apply your knowledge to the things you see around you

M: Yeah and I feel like it's really like, it's helped me understand things way more deeper than I did before, like, understanding the life cycles and the reproductive systems and the same with like in 212 especially because I want to go into animal biology it was very helpful and I was Like I just think it's exactly what I needed

N: Oh P is back

L: P glad you're back

L: Um yeah and so how about for those of you who aren't studying exactly what you want to do in this class? … mmm not so much

Q: Mmm I don't know it's kind of hard

L: Yeah I mean do you mind talking it out a little, thinking out loud like what what do you think you're going to be needing, what kind of skills do you think you'll need to be successful as a dentist or as a pa yeah go ahead

Q: I was thinking more like I guess the human anatomy or I guess more like I dunno When I thought of biolgy when I was younger I thought about the human body or like bones or stuff, I didn't really expect like plants or fungi or whatever, I didn't really expect that when I was younger when I thought of biology it was like oh ok I'm going to learn about the human body and stuff like that I just feel like if it was more about anatomy that would be more helpful to me personally I think

N: Um for me bio 211 connected more with like what I wanted to do just because I learned a little more about cells, cell cycle and everything so I could at least somehow connect it to like the medical field or the PA world, one thing I like about 213 is I work, you know we work in research groups, and In PA you have to work in a group and work with people well so that's definitely helping me to like build my skills to lik learn to communicate in a group, like figure out schedules, stuff like that

L: Yeah that's been super important, we've been struggling with that in my research group right now so. How about you P?

P: I don't know I'm thinking

L: Ok

P: Um I don't know, no idea actually

L: Yeah that's fine it's a complicated question because there's a lot of different ways to think about what do you need when you go into the future especially if you haven't had a chance to work it your field yet, it's really hard to know. So I am gonna circle back, what we tried to do with some of the changes in this class that we thought might be helpful is we worked on that, what you were talking about earlier, that prerelease sort of study guide preexam thing, We thought that might be a useful tool for students to use and it sounds like maybe eh. Like yes and no, so I'm going to pick on you a little bit an ask you to do my job. Do you have any ideas for how we could improve exams. Especially what you were talking about, like having to be mentally prepared. Exam stress is a totally real problem that students struggle with and As instructors we would love to be able to you know make that less of a problem for students so they can focus on learning and not focus on anxiety. Any ideas

Q: Um so I know this is gonna sound really bad but um I feel like so for example this quarter we have two exams, right, I feel like in the exams there's gonna be like a lot of information that we need to know. I feel like if there were smaller exams with less information on that It'd be a lot easier to I guess focus on it because last exam it was just a bunch of information into one test and it was a lot for personally my test anxiety, I have like really bad test anxiety so it was really bad for me, I just feel like it was too much information for one exam, I feel like if there were like smaller exams I guess with less information on it It'd be so much more helpful

M: I feel like there would be a lot more frequent exams if there was less information on each of them and I feel like that would be also really stressful. Like I feel like after I finish the second exam it's lieka rel

N: I personally liked in 211 that we took three exams, had two midterms and then the final at the end of the quarter, I liked that more than only having two just because at first when I read the syllabus at the beginning of the quarter I was like ok only two that's better than the three we had before like I'm ok with that. And then once I saw the first one I was like ok maybe three exams wasn't too bad now

Q: Well like I feel like with it being really research based it's better because if we had the research and also had the to study for a cumulative exam it'd be like really stressful, I really disliked the full cumulative exam cuz I feel like it's impossible to remember especially in biology, when it's like such a big concept every single chapter that we learned in the class it just feels impossible for me

P: Yeah um for me in 211 we have three exams which is the first and second and final, and you know life happens, maybe you're gonna miss the first exam or something so the final gonna replace the grade with the first midterm so in this class you just have two exams and research, if you miss one of you did bad on one, this is your grade and the percentage on exams is so high it's gonna affect your grade so much so yeah that makes me so stressful about that like you have to do really well on every exam

L: Mmhmm

P: Mmhmm

L: Yeah I've I know about some classes they do something like three midterms and a final, and you can drop your lowest score

P: Yeah

L: Just as you say, P, life happens and so the professors are like you know if you don't show up you'll get a zero But your zero will disappear and if you do badly on the first exam when you're still finding your way in the class-I'm seeing a lot of nods, yeah

All: Yeah

L: So though again, like M brought up that does mean more exams, more frequent exams so I guess that's the other side of it

M: I don't like that at all

L: Yeah and but it sounds like so that's not M's cup of tea but maybe the rest of you would choose that if you had the chance

N: I personally would

Q: Yeah I feel like if it was like a there was like multiple I guess exams or tests I feel like it would lessen the anxiety because I would know exactly what was going to be on the test and I know exactly what to study for and there's not like 4 different subjects to study for It's like one or two like specific topics, so I definitely like there would be more I guess tests and exams and all that but I don't know I just personally feel like it's better because I know exactly what's going to be on it, it's not a bunch like 6 different topics in one test

P: Mmmmhmmm

M: Yeah

L: So it's like you can study fewer topics more deeply?

Q: Correct yeah that's what I yeah, I feel like that's so much better for me personally I would want that instead instead of like 6 different topics in one exam

M: Cuz I feel like at that point you're just trying to memorize it so you can remember it long enough for one exam and then you can just like forget about it until you have to study again for the final so it's more at that point I think about memorizing for short term rather than trying to understand it for long term

Q: For me like um it’s like the oppo—it’s like the same thing but if it was for multiple exams and for a smaller amount of information I feel that allows me to Flush the information, like have multiple chapters like how we did in this exam, it made me sort of focus on the big concept and how everything relates to each other but when it’s stuff like the quizzes, sort of get it done with and then focus on the next thing and I feel like that, like it doesn’t help me, it like encourages me to forget the previous information in a way but that’s just a personal way that my brain works

L: Mmhhmm huh.  Um in terms of knowing what’s like , what you’re supposed to be knowing deeply for the exams, other students have talked about a study guide, not the preexam but an actual like list of I think it was it sounded like it was a list of questions?

Q: Like the learning objectives

L: I think that might have been it

Q: Yeah we had that in the 211 there was like a really long list of every single learning objective for the final and we memorized all those and she’s like If you can answer all of these without looking at anything you’re prepared for the final, is basically what she said and it basically was like that, once I knew the answers to all of that I felt like I could do the final

L: Did that work

Q: Yeah it worked, like it worked way better because she only did that for the final exam in 211 so lots of people were struggling with the midterms cuz she’d just give a link to every single chapter’s study guide but Once it was just a list of like all of the stuff we need to know on one space it was a lot more helpful

L: And do you have anything like that for this class

Q: Yeah that was sorta like the preexam thing

L: Ok so it sounds like that that was uh it’s from what you said it was not particularly helpful for telling you what to study because there were just the blanks that you had to fill with like everything

N: I liked the study guides more personally cuz for my—I had a different professor for bio 211

Q: Yeah stacy

N: Uh no I had doctor miller and he would make us study guides for each chapter and then have a completely different document for the final that he would already have released during the first week of the quarter it’d be like ok compare and contrast photosynthesis and this, how are unicellular and multicellular different.  So I like study guides in that way more than I liked the public exam because at least then you know maybe in once chapter there are four different questions about compare and contrasting but he’ll put it in the study guide so it’s like ok there’s a chance at least one of these will show up and I’ll know them

L: Is that the same as memorizing

N: For me personally I didn’t think so I felt like I learned more with the study guides than I did the public exam because I could go through each one as I would like read the chapter or watch the video and just like ok, I would type in the answers on the document and then whatever by the end of for like chapter 9 I think that was photosynthesis, if I couldn’t answer any of the questions that’s when I would email him or maybe talk to somebody in my discord group and say did anybody miss a question, was number 6 hard for anybody cuz it was hard for me and we'd just talk it through and people'd be like well this video helped me, or try looking at it this way. So personally for me the study guides are better than the public exam. But that' sjust for me how my brain works

L: Well and it sounds like that was more of a guide that you got before you even started the learning, before you starting studying period whereas it sounds like the preexam or the public exam you got it like very close to the actual exam, so that was more of a review tool rather than a like here's how to learn tool, am I getting that right

N: Kindof yeah, it was like he would li think he would put on top I can't remember off the top of my head by the end of like this week's module you should have a general idea of How to answer these questions or understand these concepts cuz theses will come up on the exam or, you know this is going to connect to the next chapter the next module so if you don't at least understand it a little bit you're just going to be lost the next week and the week following and again and again

L: Mmm so it was like telling you sotrt of what you deed to understand deeply so you could continue to grow in the class

N: Yeah it was like if you want to get from point A to B you need to understand these other tiny points otherwise you won't understand B at all , you'll just be kindof confused as to how you got there

L: So it sounds like that's the anti-memorization because it's like you can study B but you won't understand it unless you already understood A

N: Yeah

L: Cool yeah that sounds really useful to have that upfront

N: Yeah I kept all my study guides from 211 so they could help me in this class and 212 as well

L: Q you're like nodding along, now I want to know about your experience

Q: Yeah I actually had the same experience because I had Dr. Miller as well. Personally I honestly really loved his class, I don't know, out of all the biologies that was probably my favorite because it was like how I studied, I guess I actually like fully understood it Like the way my brain just completely like wrapped around the concepts of it and it was really nice to be like, oh yeah, like I just personally thought it was really nice it's just how my brain worked it was just yeah

N: Yeah I liked Dr Miller a lot

Q: Yup

M: For I don't know someone else is going to say something but um for me I had uh stacy alverez and it was uh a lot harder of an experience like a lot of my classmates as a whole did not do well on the midterms because like she wasn't very like helpful with the like, we didn't get study guides we just got like the general release chapter questions and stuff, and so like overall it would be like 60% is like the average grade for the midterms and so like lots of people were struggling so I sorta had to like teach myself in that class and I feel like a lot of people had the same experience, like we all talked about that but I feel like in the more recent classes it's been a lot more pleasant, preparing, I was sorta jealous of you guys having miller it sounds like he was great

N: I had stacy too, I had to retake bio211 because I was so lost in her class but once I was in Dr. Miller's class it was a complete switch, like I was able to understand everything and really get it ingrained into my brain so I'm really grateful that I was in his class

M: It's um I remember stacy saying like so like when everybody didn't do well and usually like what I've been taught is if like every single persona in the class is doing bad then it's not really the fault of the individual we're jut all not being provided what we need to be provided but She told me I was like studying wrong and I should have studied this way instead so I studied her way and it didn't go, like it basically wen the same, so then I studied like my way for the final Then it turned out really well but it was a roller coaster taking her class

N: I feel like Dr. Miller would at least warn us, he'd say ok the first midterm is gonna be a little relatively easy and the second one is going to be harder and he'd warn us before we'd start studying for it, like Before we'd go into the next few modules and I kindof that helped me mentally prepare, like ok, the next one is going to be harder which means there are going to be more detailed concepts that I'll have to be ablet to not memorize but like understand more, So it would help me figure out my study schedule more, be like ok, this is gonna be hard so I have to fully prepare myself

P: Mmhmm

L: Um I'm curious about kinda the difference between understanding and memorizing because that seems really key, a lot of students talk about that so I'm wondering have there been different courses you've taken where it's important to memorize and important to understand?

N: Um for me um memorizing has been more important in math classes, like statistics or calc just memorizing certain maybe equations or how to put things into the calculator. science is I'd say is pretty important to understand it more but you know sometimes you find yourself trying to memorize something just long enough for you to take an exam or quiz and you'll forget it right after, like you might get to your car right after the exam and be like ok I don't remember anything anymore

P: Mmhmm

L: What feels more helpful to you? Like personally, not as a student

Q: Definitely I think

P: Understand

Oh sorry you want to go.

M: I was just gonna say the study guide helped me more than the public exams because I can just take my time and go through it and those one day a week I could probably say like hey this one question on the study guide I was pretty confused Versus like the public exam you can't really ask too many questions but they have to be pretty vague because she can't give you like a detailed answer about it because there are those blanks missing so it doesn't give you much room to ask questions

Q: For me I feel like memorizing is a good milestone for comprehension cuz even if I do forget like all the details stuff and the memorization it helps me like I still remember the general concepts and all the main ideas that I need to know even if I don't remember every single step in that life cycle still

N: Um yeah so personally I um I honestly feel like memorizing is something that you have to do for some kind of exam or some kind of small quiz that you have to do but understanding is like you actually understand the subject, like you know what's happening instead of like oh I have to do this, I have do memorize that just so I can pass this

P: Mmhmm yeah I like understanding more than memorizing. Understanding makes you feel like interesting to this topic so you want to know more. But memorizing is you just want to pass the test that's it yeah

L: Yeah definitely. It keeps you excited

M: Like it keeps you eager to learn more, like you wanna learn more

Q: Like I feel like you're more likely to see how something in one chapter is connected to the next chapter

P: Mmhmm

L: And it sounds like for you all, the preexam was like, like memorize things that fit into the blanks

Q: Yeah

L: And the study guide was like I need to understand this list of things is that right

All: …yeah

L: So hmmm. Are there things that are helpful or aren't helpful on a study guide like I know that I remember study guides like some of them were like these are a list of subjects, like learn these, and some of them were like—and it sounds like you had this experience too, some were like know all the answers to this question, so is it like Do this sort of like a pretest on the study guide, is that useful or is it more useful to have like a link that you can use to study your own way, or other things that you've encountered

N: For me in Dr. Miller's class his study guide was like kind of a mixture of things, he'd be like you should be able to define like um I dunno like certain terms, and be able to tell the difference between two things, they weren't like specific like specific questions, but more like ok by the end do you know what this word is, can you define it for me, can you tell me the equation of photosynthesis, can you explain this,

L: Like it sounds like kindof not questions but like a list of skills that you should be able to do

N: Yeah

L: Ok yeah that does sound helpful

N: Yeah it helped a lot especially for the exams and to be able to see how he would word things in the study guide and in his quizzes helped me prepare for how he would word things in exams

L: Did the prerelease do that as well, cuz it's like almost the exact question right?

N: Kindof. I dunno I think cuz I really held on to how Dr. Miller taught with his study guides I tried to mimic the same thing cuz it helped me so much

L: Yeah that makes total sense, once you find something that really works for you you want it to keep going

N: Yeah you just kinda want to keep holding on to it

L: Yeah

P: Mmhmm

L: All right well I guess I'm I don't want to keep you for too long because I know students are so busy, you have much to study for, so um I wanna just wrap up is there anything that you feel like like I didn't get to that you want to share, or something that I should be asking your colleagues, your peers in order to better understand your class?

Q: Mmm I think you like pretty much like got the whole concept of what we want to talk about so

L: Great great

Q: I feel like asking about how everybody individually studies and Notetakes is very helpful in understanding cuz I just like there's a lot of different ways that people do it and if you can find like ways that some people are struggling less than others then that could be like something that's provided to the class at the beginning or something

L: Yeah maybe everyone's gonna share your, what was your method called N?

N: Oh active study recall

L: Active study recall that looks really interesting I'm gonna like google that

N: Ok

M: Yeah I'm gonna try that too

N: Yeah it helps a lot, I love it

L: Yeah so it's really cool, it's cool for me I want to thank you again for participating, for all your input, for helping me, this has been really thought provoking, it's very actually—I love my job because it's so inspiring to … Um so again thank you um we wish we could give you more but instead we just give you these $20 starbuckcs cards…

***Interview #9 May 14 2021 @CC site (Leah Lily interviewer = bold, with students C, D, and E)***

L:…We are studying some things about your Bio 213 class …and what happens is I record the conversation and then I transcribe it so basically after I get all your words down on paper I destroy the video, I destroy the audio so no one will…so there's no way that your data will be identified with you, it's all confidential all anonymous and no one affiliated with Edmonds or … I may go to your instructor if a lot of students are saying like hey we have a problem with this … yeah so any questions on that? Great. Think we lost E but hopefully she'll be back um oh there you are

E: I have a 4 month old puppy who just got past our gate so I had to go get him back to where he should be.

L: Fair enough Yeah go for it if you need to take a break during our conversation that's totally fine, I just didn't want to get started without you um ok well so my first question for you all is just how is bio 213 treating you?

E: I mean it's it's challenging to do any class in a remote setting, it's challenging I think to get all the information without true lectures um but I am enjoying doing our lab portion on campus um so I don't know. It's a challenging class but it's fun, that's what I think

L: Yeah it's nice to have that in person time again

E: Yes

L: C or D? Yeah

C: Uh yeah I was gonna say yeah like E said any well for me any science class that's online is kinda challenging because science classes are meant to be I'm pretty sure in person, are meant to be taught in person most of the time Cuz it's you know there's the lab aspect of it and everything and to take in all that information it's a lot to learn on your own I think and the videos help and stuff but I feel like the majority say when I did take science classes in class it was always a little easier because of the lectures because then you learned in class and you went home and just studied what you didn't understand or did the homework, but here it's more You learn it and you you know you do the all, you put in all the work um so it's definitely more challenging but like E said I think the lab portion of it going into cl—or not into class going into the farm doing all that definitely helps you know, lets upi breathe a little so that's good yeah

D: Um I do really enjoy the lab, for this quarter cuz um I love going out and doing some stuff outside but for the lecture I'm kinda struggling to understand the information that I read in the book and try to um memorize how to word the life cycles example like for the last exam is very big challenge for me because I basically it's really hard to understand how to draw and how to um find a good way to draw the life cycle and you know the lecture was kinda a lot and like if we have like a lecture day so you go over everything then Pick in the important details information so you know when I read the book again I think it's better for me to memorize the information

L: From the lecture, not from the book?

D: Of the lecture yearh cuz you know once she talks to you have like to pay attention to something that she picked on yeah.

L: Yeah that makes sense um and you mentioned how you were doing it for the exam, do you feel like the exam is is, do you feel like you're getting prepared for the exam by the work that you're doing beforehand

D: No haha

L: No Haha ok

D: No I'm always struggling how to study for the exam

L: Yeah yeah I mean that's the difficult part of any class right so that makes total sense. And for you others do you feel like the exam is, do you feel like you're getting prepared for the exam by the course activities?

E: I feel like for me and to get prepared for the exam I had to do some of the like well I guess I mean even watching the lecture videos isn't necessarily required but I don't learn super well-- sorry I'm very distracted by the dog sniffing everywhere, I just took him out-- I don't do super well with reading and comprehending really well, I do a lot better with hearing it or seeing it so um it's very helpful for me she has um a lab gab like office hours time and I go to that on Wednesdays cuz it's really helpful for me to be able to like interact with the content of the chapters, talking to her about it and hearing her explain it and seeing her draw it on the whiteboard and um the best thing for me though is to explain the information to somebody else so I mean sometimes I'll just, I have a seven and nine year old and I explain to them life cycles because like they don't really understand what I'm talking about but I mean they also think it's cool and their brains are little sponges so maybe if they decide to go into science one day they'll you know already have a good foundation But that's really helpful for me to explain it back to somebody else but the way that the, the way that learning the content of the chapters is set up is not very helpful and it's hard because currently I'm taking um 2 science classes and a math class all remote and um so it's hard to do things like outside of what's required cuz there's just not time. And I just have zero time in my life so um yeah. So it's definitely hard to learn the content and at the farm like when we're doing lab there's not a lot of time to ask questions um within our groups At least my group we talk about some of the information and that's helpful um and but yeah I just I wish that there was more interaction with each other and the information and I have to go get the puppy I'll be back

L: How about you C?

C: Was the question about studying for exams

L: Yeah

C: Yeah um I feel like if we had some kindof like way to practice I know that there's like no way to practice unless like you give the exam and then but if there was like you know maybe like a group activities or something more of like interacting with others and discussing what you've learned I feel like that'd be helpful, like the study guides are also helpful I feel like cuz they focus on what you learned in the book but it's difficult to just sit down and read the book like I don't it's hard you know you get distracted or Me personally I get distracted a lot reading a textbook so I feel like yeah if there was like a-- or and like E was saying the lab gabs those definitely help because you know you're able to ask questions with wherever you're confused or everything and then Dr, And then you know she discusses that so that's definitely helpful um but yeah it's it's difficult to study for you know over what you've learned cuz it's like you have to read the whole textbook and that's tough but yeah

L: Are there opportunities for group studying apart from the lab gab

C: I mean yeah we can, she encourages us to you know get in our groups cuz we have like little small groups On the lab or at the farm and to study with those people, to discuss what we've learned so yeah, definitely take the opportunity for that as well so that's also helpful yeah

L: Um and any uh any of you also take advantage of the small group studying sessions? Studying with your small group or otherwise E you mentioned talking to your kids about it but are you doing any other like group study activities

E: Yeah I might sneeze in a minute um. We have a discord group chat with most of the people in the class and a lot of us have taken 211 212 and 213 together um so especially before exams um it probably makes it more sense to do it like weekly and not just before exams But um like we there's an opportunity in there to ask questions of each other and help each other out with the information which I felt like we utilized more last quarter than this quarter but um and then within my research group we definitely talked about the information And in our research group we have a discord chat just between the five of us where as we go through some of the information or study for tests or before we take some of the post lecture quizzes cuz those are definitely the more challenging quizzes that we take making sure we understand the information cuz those quizzes are a lot about comprehension And taking it out of context in a way just out of the context in a way, just out of context that they put it in in the book and being able to apply the information um in a different context and being able to understand if we really understand ha how the information fits in outside of the specific example that they put in the text um so yeah we have study groups there and then the lab gab which this week it ended up just being my research group so we ended up talking more about our research than the chapter And then before tests also dr gwen normally does a study group or two and I feel like those are pretty well attended within the class and fairly helpful to get more specifics on some of the harder content.

L: Yeah that makes sense.

E: And the puppy's in its crate now so

L: Haha

E: Kids are outside, puppy's in his crate, hopefully I can focus

L: And probably a rare moment of calm [cough] scuse me. And you D have you taken advantage of any of the group studying opportunities

D: Um I think I have not cuz um cuz I've been working a lot so I really don't really have a lot of time to go to any zoom meeting especially when dr gwen have like a lab gab is very rare for me to [something] the meeting because I try to really make time to go in there But still I just too busy with my work

L: That's hard, it's hard when you're working a lot outside of school too. For you what do you think has been the most helpful thing helping you get prepared for exams?

D: Um the way she put out the outline for the exam but the thing is like even the outline so the outline for the exam still have like a lot of information to get in my brain and plus like I speak a different language so it's kinda difficult for me to digest some kind of words so I can try to understand it in a good way for me to do the test. Plus um there's a graph and a phylogeny which I did not have any practice on it, especially for the graph so the one for the graph that I drew need some practice before I can get into the test because Basically I know nothing about it

L: Oh no

D: I cannot just like um didn't know, no practice and to do with the test this is a new hassle for me, I listen to her try to explain further but still I don't know what's the, I need to look at or what information I need to put in the test to answer the questions

L: Yeah I've been hearing that from students that the graph and the phylogeny both were really hard questions and you didn't feel like you had a lot of like preparation for those particular types of questions in class

D: Yeah yeah

L: Is that yeah. And uh the outline that you mentioned is that the preexam I've heard people talking about

D: Yeah, it's really [something] to Give us like information she wants us to know will be in the test but it's still a lot

L: Yeah definitely. So it's kindof a study guide

D: Um yeah I think so yeah

E: Well and this test on this quarter it's not really a study guide like the questions on the outline are the questions that will be on the test. So last quarter the compare and contrast part of that test study guide, There was like 10 different questions and she normally would pick one of those 10 questions but it was like 10 very different questions whereas this one there were two questions and she was gonna pick between the two. One was a lot more specific of primary and secondary endosymbiosis so it was like very this is what you want to know but the other question was much more open, of comparing and contrasting different taxa Within I dunno fungi ugh I don't remember anymore um haha but it was like there was a lot of different species that she could have asked you to compare and contrast. So that one was just a lot more open ended and between the three chapters that the test was on It was just a lot of information. With the graph, I do with Edmonds would require students to take probability and statistics before they do this cuz like 212 and 213 is much more research based and it is very hard to, I mean I did take probability and statistics I wasn't super, I didn't love the way that Edmonds teaches it but um, and I still struggle with some of the graphs and, but I feel like at the same time in comparison to some of the students that I've talked to who have not taken probability and statistics there's a much more like lost feeling of how to do this, The math to display research in an effective way and how to read these graphs cuz like the graph on the test was like a box and whisker graph which you would know if you had taken probability and statistics and if not it's up to the student to do the research to figure that out because We're not learning types of graphs and how to read them necessary in biology.

L: Yeah and D it sounds like you had to basically learn how to do deal with graphs on your own

D: Yeah I have to do my own because I basically don't know what's the type of graph that is in the test and I tried to understand how to read the letter and numbers on it it's kinda difficult for me to learn, I wish she could like have like example for of graph and then have an answer them all so we just like try to do a test on our own but look at the key answer and compare it and that's way I think I would learn the best

L: yeah, Like a practice test?

D: Yeah

L: And it sounds like C that's what you were looking for too, the ability to have practice problems.

C: Yeah I think that'd been very helpful because We'd know what we were doing right or wrong if there was an answer key whereas if it's just you know open ended then you have to do all this research and at that point I don't even know if like what I'm searching up is correct or not you know like so if there were like an answer key that would've been helpful. And then um like um E and D were saying about the graphs, yeah like I had to search it up more so you know find out what the R's mean like standard deviation all that but I think after a while it kindof it kindof made sense but still it would be nice to have more practice on it definitely.

L: Yeah and it sounds like reading a graph or doing a phylogeny, that's a specific skill where you kinda, it's like doing a math problem you kinda just have to like practice doing the problem to kinda learn it. Seeing some nods

E: Yeah the phylogeny we do go through very in depth especially in 211. The students who are in 213 who have skipped 211 I think they're struggling more with the phylogeny, and she did, Dr. Gwen did put up a couple videos and post a couple videos from last quarter about phylogeny but that is one that we did go in depth on, the graphs we have not gone in depth on, like that one like I I said if you haven't taken probability and statistics I can, I mean I felt a little lost on some of them and I did take probability and statistics, So um yeah those ones are a lot more challenging

L: Mmhmm D I saw you unmute there, something to add?

D: No no I was trying to say the same thing as E

L: Ok all right

D: Yeah

L: Um all right so this gives me kind of a picture of what your exam was like, and I guess some of the materials you got to try to prepare for it. So I have kindof a bigger question. For this class, is it—does it feel like it's helping you prepare for your life after this class, like your career or, either your college career or if you know what you think you wanna do afterward Any of those skills carry over or what you're learning going to be useful to you?

D: I think I learned the most in the lab, especially how to do the research and um work as a team with everybody um try to communicate with everyone as a team and then try to do all um um sorry what's to say, separate work and try to work together as a team, I just all about teamwork and research, that's where I learn the most

L: Yeah absolutely, collaboration is a crucial skill. I experience that in my research team all the time. Um yeah absolutely right. And how about the rest of you?

C: I think that the research definitely is something I'll take away from it cuz we did it both in 212 and 213 so learning how to write the abstract you know the whole research paper and all that and also collaborating with Other group members in order to you know complete the research I think that's definitely a helpful task for the future, and you know in a science career and everything so I think that's' definitely something I'll take away from this

E: Um I'm in a slightly different position than most students because I already have a bachelors degree and I'm working on prerequisites to hopefully get into the genetics counseling masters program at the UW

L: Oh wow

E: So um like I already can see, I do really like the way we did this exam because even in the career no boss is gonna ask you, they're not going to surprise you with something I guess in a sense where you have, You haev the opportunity to work on skills to build skills if I wanted to move up within a position or company or whatever like I have an understanding of what's expected of me in that position and the opportunity to learn and to grow and to take classes and get that experience or knowledge that I need to be competitive to make that next step. And so I liked the exam, I mean I have a hard time with tests in general just because no job is necessarily going to- you always have the ability especially in our day to grab your phone and google something, so You have the information accessible to you and yes there is the need in a program like this to show that you have like an understanding and you don't always have to go back to the information to get that, and so I liked that because I felt like this balanced it out. Where I had the information, like the graph that was on the outline is the graph that was going to be on the test, the phylogeny on the outline was the phylogeny on the test, so I had that ability to have that information, See it ahead of time, google, read, whatever I needed to do to have that information and then be able to take the test on it and so I felt like it was a much more realistic view of what life looks like and a lot of this I will take, I mean I did a lot of research in my undergraduate degree cuz it's in psychology but this is very different research that is more I guess objective and science based whereas psychology is much more subjective and so I'll take that as well but a lot of what I'm learning is the foundation of just the math and science that I need to do and like for this class I'm only taking it to finish my AA in Biology cuz it'll look better on my application. So somewhat I've been enjoying it, I'm a horrible gardener and I'm enjoying learning about plants it might make me a better, I don't know, I do a lot of running, I ran through a park the other day and I kept like looking at the plants and like iden---kinda identifying if they were vascular or nonvascular and I was like what is wrong with me and but for me it's more of the experience and the--I already have the prereqs done that I need to take and so for me this is just more like getting that foundation in science that I need to get into a program that's gonna bring together natural science and psychology

L: Yeah that's really good to hear, um we, one of the things that we tried with this particular course is we did, we wanted to try that sort of outline or that prerelease in order to see if we could make the exams more, I guess a better experience and it sounds like for you that was effective or at least in some areas. And I'm curious I'm gonna circle back D to something that you said which was that because of struggling with Understanding the language, one of the things we were hoping with releasing some of the questions on the exam early or at least part of the questions, is if That would help with strugg—like because instead of struggling to understand what the question is having you do, you could figure that out beforehand and then go into the test already knowing.  Did that help at all or was that just Not effective?

D: You mean like she give the specific question like 10 questions and pick one of them for the test?

L: Yeah

D: That's helpful because I know exactly what I need to study and getting definite otherwise if the questions for the last exam, for the outline I'm kinda going around and try to guess, maybe she gonna pick this maybe pick that and I kinda mix them up together so I can get them in my brain cuz it's so so similar it's real hard for you to get something if this is your second language and um and it's something like you never read about this before. I tried to read, I tried to understand but you know I still have to go to Google Translate a lot

L: Yeah

D: I really struggle for it

L: Yeah I really admire you taking especially taking a science class in your second language so well done, good work um yeah but it sounds like it sounds like you were like able to go to google translate and look up some of the stuff in the exam beforehand

D: Oh no I just translated words to try to understand what the sentence Tried to tell me what's the point of it and um you know I tried to get the general idea out, I cannot just go very in depth for some information because it feels too much, it feels overwhelming For some thing like like the um what is it called the contact or some such like compare and contrast different tax, there's a a ton of taxa so I have to know like the scientific name and what's the difference between them and what is common between them, it's really difficult for me to understand and know and memorize it So if I know exactly the question is I might try to get you know, try to memorize every day, I really don't like guessing around cuz it's hard.

L: No yeah it's easier to be prepared when you know exactly what it is, I guess my pushback is if you know exactly what's going to be on the exam then you study just that and you don't study the rest of the material so There's a tension there

D: Mmhmm yeah that would be the gap, I can tell but you know cuz biology is just a lot of informations, so um I think what's important to remember is that's that's going to if she can point out, that would be good

L: Yeah um yeah.  How about you C, was that prerelease the outline was that helpful for you in preparing

C: I think yeah, it was helpful especially since see cuz like one of the first

L: Oh I think you're cutting out oh no try again

C: Ok can you hear me ok now

L: Ok yeah

C: It cuts out I dunno I did like the way that she did it, it's um, or the preexam thing, um but yeah like for instance the compare and contrast it's just like one was more specific asking the question than the other one, the other one was more general and so the more general one You had to look at different you know taxa and that could be like anything you learned in the book over the past three chapters, cuz it could have been you know fungi or Plants or you know protists but the second question was more specific so I don't know if there were maybe like a balance of the questions asked so they are similar I don't know if you can hear me

L: Yeah I can hear you you're frozen but I can hear you

C: Yeah cuz your guy's faces are also frozen so I wasn't sure, ok, Now they're unfrozen but yeah if there were like a balance between the two so that they're either both specific or both more general so there's like a good in between of that. So that, Cuz like I focused for instance for the one that was specific I focused on that question because I knew exactly what she would ask pretty much, cuz it was like endosymbiosis primary secondary, and then Taxa it's just like it's general so you didn't know what exactly to study. But I think that this was definitely helpful. I also like the way she did it in 212 where it was like a lot of questions but they were specific, like they were the questions that she was gonna ask. So it kinda did make you look at the overall whatever you've learned throughout the whole chapters because it was you know Like a lot of questions so I feel like they're both good ways to practice for the exam because like the previous one it makes you look at everything you've learned because you don't know which question Out of say like 10 or however many she put on there you know so yeah, they're both good ways I like the way she does it.

L: So it sounds like that is helpful for preparing for the exam, and my question now is did the exam help you prepare for gaining this knowledge or using this knowledge in your careers or in your life? Like did the exam seem relevant to your learning, not just passing the class, if that makes sense

E: I think so, I think a lot of what we're learning um is um how can I say this so like our research this quarter was a lot more structured um where we're put into three different groups and I'm in a pollinator group and like there's stuff that we're studying with pollinators and the plants that they're attracted to but then at the same time we're planting a pollinator garden that doesn't completely, like we're not just putting the plant that we're studying in the garden and so, so I feel like with what we're physically doing in lab there's a lot of under--like what we're learning in the book and what we're physically doing in lab I feel like complements each other a lot and it's stuff that we need to know a lot of the stuff um in order to be successful with passing the class but then at the same time if anybody, I could definitely see if anybody is going to go into plant biology or something else ecology that's more within the realm of biology That this class is giving a really good foundation um to all the classes and things that would build upon it.

L: Yeah thank you. Other thoughts? Whether the exams help with your learning in general

C: I think or you mean the exams themselves, not the studying aspect just the exams?

L: I guess both, I mean hopefully the exam will feel not like it's just a tool to measure you but it's also a tool to help you measure your own growth or to help you keep track of what you've learned

C: Yeah I think the way she does them with the essays it definitely allows you to show what you've learned whereas multiple choice it's like you can guess and choose but Here you actually have to explain what you've learned and I think that's a good thing because even if you didn't understand like one thing for instance You could focus on something that you do understand and still get some of the points, you know, so it's not like point-blank you either know it or not you can still like show what you've learned so I think that's helpful

L: So it's like even if you don't get a 100 on the exam you can still look at your answers and be like look at all these things that I've learned

C: Yeah yeah exactly cuz like you said you look at it and you know what you've learned and what you still need to focus on So yeah mmhmm

L: I guess that's good how about for you D?

D: Yeah the exam really helps me to learn especially I tried to on the, amaze myself if I look at the exam and look at the score if I got a high grade it's like whoa look what I did, I just amaze myself how I did it. But the thing is like it's kinda a little bit stressful to look at the grade but overall it's really helping me do Deep studying, comprehension the material and you know something is new for me and so this kinda like um I just already tried and [something] I just try to read through it again and you know try to understand it and um then maybe next time I can do it better

L: That's beautiful, a growth mindset is what we call that in education, where you're saying next time I can do better, and it's Really effective, well done. Um let's see I'm conscious of time, I don't want to keep you here for too long because I know you're very busy students all of you and have a lot to do so I want to start wrapping it up and I want to ask is there anything either that you are dying to tell me about the course as a whole or about the exams, or is there something I should be asking you students in order to understand how the course is working for you?  I can wait awkwardly for a while

D: Can you tell me the questions again, I'm kinda confused

L: Sure is there anything you feel like it's important for me to know about the course, and is there anything you feel that it's important that I ask students about

E: I think if these, if the exams are going to continue like if this, continue to happen in this format it'd be interesting to know what students think of doing this in a remote setting where a lot of the content from the text is being learned remotely versus you know whenever we get back to in person settings and if that's gonna change the way that people see the exam and the content

L: Yeah good point, I think it's pretty different for students in the digital world just like you were saying And other students have said that as well, yeah good point.  D or C, anything that I should have asked about that I didn't?

D: I think you asked pretty much so I think you covered a lot haha

L: Ok good to know

D: Yeah yeah

C: Nothing that I can think of most of it yeah you covered a lot of it so yeah, all of it you know

L: Good to know, thank you.  So then I have one last question where I will make you do my job for me and think about if there was one thing you could change about the exams, um and that includes the material you were given to help prepare for the exams, if there was one thing you could change, what would it be?

D: I would love to have like example exam with an answer key like below

L: Ok So like a practice exam with answers, with answers. thank you.

C: Yeah I agree with D, to have a practice exam with the answers I think that definitely will show what also you know help you understand what you've learned and what you still need to focus on because you have the answers, and probably just having the questions at the same like level of difficulty for you know, so that when they're both, not the same level of difficulty but at the same, Man how do I say it, so that one is not as specific, or one is not very specific and one is more broad but so that they're both around the same like, they give you, like, Man I can't say it haha, like you know what I mean like so that one is not very specific and then one is very broad because then you don't know which to focus on but so they're like equally distributed with the material

L: Yeah so like what you were talking about where with one of them you could study exactly what it was gonna be and the other one You had to study like 60 things

C: Yeah exactly eyah so that can be more balanced and yeah

L: So like, so like the example, or not the example questions, but like the questions that show up on the outline on the prerelease like they should, you should sortof treat them in the same way like you don't want one question to be like "use this to make your exact answer" and then the other one to be like "use this to decide to study 60 things"

C: Yeah

L: Is that correct, what I'm saying

C: Yeah that's what I was trying to say, but I couldn't phrase it correctly

L: Yeah that's a good point because if you have to think about like how you use the tool then it's much more confusing

C: Mmhmm yeah

E: I think like with that one playing of what C said is um it be helpful with that much more open ended question to have help structuring how to study I guess in a sense so maybe it's an open ended question but it's like 3-4 different examples of like compare and contrast different fungi or protists or like what's different about this taxa protist versus this one Cuz it was a lot of, it was so open ended like it felt a little overwhelming to figure out how to like funnel in to different parts of it so, you can have an open ended question like that but maybe a little bit of help or a study guide, like I spent way too long going through each chapter and taking the like table in the book that went through you know this different part of protists and this one and this one and um and like just putting it into one document so cuz three chapters of all this information that isn't just the species and you know how they reproduce and how they eat and how they do this and that and that and can they do photosynthesis and whatnot, it like it just it was it would be nice to have something with an open ended question that would help focus the studying I guess is the short answer to what I was trying to say, to what I just rambled about um

L: Would that be kind alike what D was saying about actually providing answers, like example answers

E: I don't think you need to provide example answers, on the graph yes I think that would be really beneficial Cuz I feel like in the study sessions too that's what we spent the most time talking about which was a little frustrating just to me Only because I took probability and statistics and I understood a lot of that but I also understood that most students needed a lot more time in this part so I think it would be great to have a practice exam on like especially in 213 where students can go from 211 to 213 on the phylogeny and the graphs because that is a really easy one to do like an answer key to you know like sister taxa, maybe there's two or three on the whole thing and so it's easy to say what is, like identify a sister taxa on this phylogeny and then say this and this are sister, this and this are sister. Whereas compare and contrast it'll be really hard to give an answer key to that cuz there's so much that you can compare and contrast Within these different questions so for that one there was the endosymbiosis which was easier to study for cuz you looked at it and you're—and it was challenging to understand in the first place, though like That was one like understanding what was primary and what was secondary was, was a little challenging so you still like that question you still had to spend a decent amount of time to really understand what that looked like. The second one being so open ended what I'm saying is If there was the ability to have maybe not an answer key but just a document that goes through like takes the phylogeny of protists and then the example species like within the phylogeny and goes through different things of you know do you wanna focus more on like one question could be like compare and contrast the different reproductive systems Within fungi and which that was something we were doing any ways of like drawing life cycles, and or like compare and contrast you know protists and or like looking at which goes to endosymbiosis looking at the algae and cuz there's three different ones I think that the whatever however they discover the stuff, that their ability to do photosynthesis came from endosymbiosis of a green algae whereas the other ones were red algae, so like what is the difference between those two, what does that look like, you know, Do their, do the way that they absorb light seem different so there's a lot of different things that you can do within all those things Being able to like focus down to that of How do those secondary endosymbiosis organisms look different from just your normal average tree that does photosynthesis--which I feel like was a question on a quiz like somewhere, like I feel like I answered that. But being able to like focus it down onto some of the—cuz there's just so much. There's so much about reproductive cycles, there's so much about photosynthesis, There's so much about um what they eat and how they eat like fungi decompose things and then absorb it like --so taking that broader question and helping to focus down into even just 8 things or whatever that are still gonna force students to really study and understand the different taxa and organisms and life cycles and things like that But just have more focus on how to do this rather than 60 pages of information of like where do I even start to study for this question.

L: Like so it sounds like more, more help and more examples of how to solve the problem, rather than the actual solved problem, does that make sense

E: For the compare and contrast yes. For especially for that super open ended one

L: All right yeah so it sounds like, it sounds like some practice problems, with answers, and some examples of how to think about preparing for the bigger more like general questions would be useful for maybe all of you? Seeing nods. Ok thank you this was really really helpful, um ok um I guess that's all I have and it's been 50 minutes …

### ***Interview #10 June 14 2021 @CC site (Leah Lily interviewer = bold)***

*Note: This interview is with a repeat participant from an earlier interview*

**…all right well yeah so um you finished!**

Almost, like I'm real close

**All right well good luck and congrats because you're almost there, you can do it. And uh you just took a final yes in 213?**

Well it was like our second midterm so like instead of finals we have a research report that we like work on for the entire thing and then the report is due at the end so it's sorta like our final cuz it's worth 75 points

**Oh wow yeah /** It's a hefty paper **/ How did it go?**

So like right now we're working on it, our draft for the paper was due like a couple days ago so now we just have to refine it and stuff and since it's like a group project it's like a little scary. Shitshow

**Yeah cool and that's due Friday /** Yeah on the 18th I believe is when its like officially due and that'll be my last assignment

**All right! Well you're almost there cool. Yeah I do, I heard the research project was like really cool for people but as I think you know we are focused on the exams for our research and so /** Yeah / **So I'm curious about how the second midterm went for you**

So it was sorta like, it went less good this time only because like, so since there's like a couple options for the questions you can get in each section of the exam and I had like preferred questions because some of the –there's always like one that's very specific and one that's slightly vague But she can't tell you like any more than that obviously but like I got both of the vague ones for the drawing and for the um essay and I still think I did good but I was hoping for the other ones

**Yeah**

Those two were just harder to prepare for

**Yeah oops doodoodo sorry ok um yeah that's what I remember hearing from the focus groups the first, after the first midterm and so I'm curious about that if you could tell me a little more about like what did you do to prepare for a specific question versus a vague question**

So for the specific question versus the vague que---for the specific it was um compare and contrast vascular tissues of xylem and phloem so it was very focused on like a singular chapter that we had and it seemed mostly that it was only testing on like three of the five chapters Which was sorta nice but um so I basically went through that entire chapter on the um vascular tissues in plants and that was really easy to prepare for because I could just basically write down everything that it says about those individual vascular tissues and then I could write an entire compare and contrast but then The other question for the essay was like um what was it, it was like uh compare and contrast adaptation, phenotypic plasticity and acclimatization Which are like slightly like they're bigger ideas I guess,

**Sure**

And then using ideas like that to find like examples was slightly harder to do since like adaptation and acclimatization were like sorta more general things that we learned in like our first biology class and then comparing it to something like phenotypic plasticity which is sorta within acclimatization so it's slightly more difficult to prepare for Especially the drawing prompt for the vague question it was like um it was different because it asks us to draw a phylogeny for the vague question and the phylogeny-- but we had no phylogenies in those exact chapters

Oh ok

So the idea was like what she said in her zoom was like high school level is to memorize the phylogeny and college level is to be able to take the information and put it on a phylogeny you already sort of know but it was like a lot more difficult because the chapters didn't necessarily say exactly where those things go

**Ok**

And there was like a whole section of the question that we didn't know but, and it turned out to be like angiosperms

**Ok**

And so That part would like completely catch you off guard but um I don't think it was too difficult to prepare for at all, but it was still, it would catch you off guard

**Yeah, And I guess that's like some of the exam does have to catch you off guard otherwise it's just a homework assignment**

Yeah I definitely agree with that though lots of students don't but I think it's sort of the point

**Yeah yeah is so it sounds like you had to like apply a lot of knowledge for this question um how**

Yeah

**How did that feel haha**

It was definitely like it's slightly more difficult so it sorta, it like broadens what you're trying to remember so I guess it helps more with like understanding but sometimes when you're panicking about an exam you're like I don't want understanding I just wanna know, But at the same time you do have to understand things

**Yeah**

So it was sorta just like in the moment you see what question you got and you're like damn but

**Haha yeah the stress and the surprise and yeah it's sort of unfair that the emotional reaction really does take so much out of you like during an exam. I wonder if, did having that that, she calls it the preexam, right?**

Yeah the public preexam or something

**Yeah the public exam, did having that help make it less of a surprise or was it**

Yeah

**still like well it's gonna be one of these two**

It definitely is like less of a surprise because if we hadn't had the public exam I would have studied All 5 of the chapters and had like less knowledge on each of the things and I don't feel like I would have remembered the exact definition of phenotypic plasticity as well as like when I saw the question and was like, I really do need to know this for the exam. Or I dunno like it definitely it focused on like 3 out of the 5 chapters so I would've, all the effort I put into those other two chapters would have been lost and I could have spent my time on the others, so it was good to narrow my focus a bit.

**Gotcha, my pushback, just to argue for the sake of arguing, would be, what about the knowledge that you would have gotten from studying those two chapters? It wouldn't help on the exam but what about for your knowledge?**

I think that specifically just because of the two chapters that were ignored I feel like it is ok if it was any of the like, if it was switched over, because it was plant sensory systems and plant nutrition which were like far more specific things that I already sorta knew because I took a horticulture class in high school

**Oh cool**

So personally those chapters I sorta like already had in my brain but it, I feel like it defiantly did focus on the most important chapters because it was like the angiosperm reproduction chapter is like 90% of all plants ever and then the vascular tissue is very important and like plant form and function, so I do feel like it was prioritized well but I also can't talk for like a lot of the people that might not know the other two chapters as well

**Ok but for you it felt like the, what you were told to study again for the exam was like more of the focus, right?**

Yeah it was sorta like what I needed to study and remember more because I didn't know it as well, it was like new stuff to me

**Cool. Um and how, for the specific question, that sounds like it was kindof a different—or yes, the specific question, that sounds like it was a little bit of a different experience because you knew pretty much exactly, like it sounded to me like you basically wrote your exam, or your answer before the exam yeah?**

Yeah basically, so like for the essay prompt xylem versus phloem I wrote an entire practice essay just to like, like first before restudying to just see how much I retained after the first time reading the chapter and then afterwards I like went through the chapter and sort of wrote down like a list of stuff for each of the xylem and phloem and then I was able to sorta incorporate more into the essay, and so I was really hoping I would get that prompt because I was like so ready

**Yeah that sounds, um excellent studying very good, hah which is I guess probably you know that really well now**

Yeah

**Even though you weren't asked about it but it's frustrating that you didn't get to show it off**

Yeah I would have liked to

**How does that feel to have done all that work and preparation and then poof**

Its it sort of sucks only cuz I know a lot of students say they don't study until the day before, and that would give me so much anxiety and stuff, Like I study for like weeks ahead and stuff so it's like, especially if I don't get the question that I'm like super ready for it does feel sort of sad to have it poof away but at the same time I feel like I have Bragging rights or like a conversation at a dinner party, like you know lots of stuff and you can talk about it

**Yeah bragging rights, I like that, I like that um.  And remind me what, do you have a career goal or a pathway that you have**

Yeah so I wanna be a zoologist and work with exotic animals and stuff so I was like really excited about these classes because I feel like they're finally like focusing on stuff I wanna do

**Yeah yeah yeah though I guess xylem and phloem aren't exactly animals but probably you have a lot of botany and zoology overlap I would think**

Like I also I love horticulture and I took a lot of classes for horticulture so I know definitely in the future I wanna do that but I won't make money off that probably

**Well I mean do what you love**

I love to know about it but I know a lot of people feel like if it's not like completely gonna be their career they don't want to learn it but I've always liked to learn things just as like a hobby kinda thing, especially biology things

**Well that's great, good, I feel like you're going to be an excellent contributor to biology and zoology with that attitude so that's really cool**

Yeah I hope so

**Yeah I think I said before but it's so inspiring to get to do these interviews and talk to students who are realy motivated So thank you um, I'm gonna go back again, so the specific question for that one did you get the whole question before the exam?**

Well um, so it was, so there's always in the compare and contrast essay prompts it was like Even the specific one would have a second part where she's like can you expand on this one thing and you won't know what that thing is

**Ok**

And so like there's still an unknown component but it's not like the majority of it.  Cuz this one it was like xylem and phloem and I forget what's, cuz I was so focused on the other one cuz I was panicked about the other one, but there was an end part I velieve, cuz there's always like a little blackedout spot that's like "unknown component" but it's not like the majority of the essay.

**Uh all right for I guess a little bit of a different direction, you mentioned getting pretty anxious about these exams, is that like something you've struggled with for a while**

Well it's like, like I tend to overprepare but I know there's like, like it's stressful but you, it's not bad to be overprepared but like it can like take a lot of your time and mental health which is why I feel like I wonder if other students who are waiting longer to study are doing as well or something but that's obviously like information that I would never be privy to so I'm gonna continue to overprepare

**Sure sure**

But it's also like when, my grandparents are paying for my classes so you also have the external pressure you know you have to do good cuz somebody else is paying for your education, like you don't want to fail the class

**Yeah you're gonna be like oh I don't wanna let my grandparents down**

Defeinalty since they're like ex professors so there's an extra level right there

**Wow, Cool. Um. Ok. Um and then so it sounds like for you the way you basically manage the stress of exams is you just work really hard, is that about right**

Yeah I just keep on going

**That's awesome, that can be very exhausting but that is really incredible**

I die for like a day afterwards every time

**Haha does the prerelease help with your preparation**

It definitely does, the first exam I sorta like, I changed my studying a little bit for this one, um, for the first one I did sorta like-- I wanted to make sure I had like complete knowledge for all of the chapters beforehand and all that but I feel like that's because we only had three chapters so it was like easier to do that and then I used the last like 4 days on the prerelease and like answering those questions and stuff. But for this one since I realized that Two of the chapters weren't on there, like I didn't do my full on like entire day of each chapter thing before using the prerelease because she released the prerelease a little late and then like it was more just like I'm gonna focus on that and also she was like, Just use the prerelease like what are you doing I dunno she

**Oh like she told you to study just for the**

Yeah she was sorta just like don't waste your time I was sorta like oh ok sure.

**That's sorta invalidating**

Yeah a little bit but it also did help in the end so I was like I guess so, I sorta feel like I'm still gonna do a day on those other chapters after the thing just because I have a collection of papers I composed for each chapter

**Oh wow**

Yeah because that's how I prepare for midterms so I have one for each of the midterms, or chapters through 211 through this class

**Ok all right so I guess if I'm getting this right, it sounds like the prerelease part of the exam helped a lot for studying FOR the exam,**

Yeah

**maybe like helped your grade improve um like maybe**

Uh you know

**Maybe but it sounds like it didn't, it didn't, it didn't really help with your knowledge acquisition, is that—**

It helps for like the knowledge specific to the exam cuz I won't say like it didn't help at all because like I don't feel like I would have remembered xylem and phloem as well because like when I saw that question it was sorta like a, at first I was panicked about it because I didn't remember too much about it, xylem and phloem that specifically, but now I have xylem and phloem absolutely down. So it's sorta like, it definitely made me know certain things very well and then the other things I probably would have known like the same amount regardless

**Ok ok. So basically helps you learn the things that were like highlighted way more in depth?**

Yeah

**The other stuff was kinda like, a little more up to you but YOU're like a really high achieving student so you do it anyway?**

Yeah exactly

**Cool, you know, again, I say like excellent work, it's just it's really cool to meet students who are willing to go above and beyond and take their own learning seriously**

Yeah

**So ok. Help me out here then, because basically when we were trying to think about what we can do to help students prepare for an exam, we're thinking, all right, we'll give them you know that prerelease that's exactly what's gonna be on the exam with some stuff withheld, ok. And what we're getting is it sounds like it does help people prepare but I'm, like partly I'm worried about students who are really dedicated already who are only gonna study for the prerelease maybe they're gonna lose something in terms of focusing on their knowledge, What do you think about that about maybe your peers' experience or other people that you have talked to about the exam at all**

I feel like it definitely helps um the students that don't really want to put in a lot of work into the other chapters cuz they just like look at in then they'll be like ok, but um. Yeah I definitely feel like it's a little bit of a short cut for a lot of people but it can be valuable if people are responsible with it

**Mmmhmm that makes sense**

Yeah I sorta like, I feel like people definitely can be irresponsible with it if they choose to but it's also just depends on somebody's motivations as a student, if somebody's not, I feel like if somebody is trying in class in general they're gonna try for the exam and like overall but it just depends on the amount of motivation that each individual has I feel.

**Sure**

I know I heard a lot of people like even though they had the prerelease they didn't really study until the last day anyways

**Yeah that's always gonna be a problem**

So I feel like regardless of it they wouldn't have changed much of their habits

**Yeah**

Unless those people feel like it's a scapegoat, it's a possibility

**Fair. Um yeah did you feel like like uh, cuz I know you had a zoom that was I think specifically dedicated to preparing for the exam is that right**

Yeah there was a few, she did more zooms that were like let's talk about the previous one and it'll help you with the future one Cuz There's always like a phylogeny section for both of our midterms so when she was helping us figure out our last, like our answers that we got wrong on the last one and how we should have answered it It was like a really good opportunity to know how to do it for the next one Because if you ask her a direct question about any of the questions on the public release she'll usually be like I can't tell you that.

**Ah**

Which is fair, which is why she's more focused on like ask me questions about the previous or maybe a clarification about the second exam but it can't be like which phylogeny do you want me to study cuz she'll be like I can't tell you that

**Right, that makes sense, yeah that sounds hmm**

So it was harder to get help on it just because like--But at the same time I wouldn't have known what to ask for help for if I didn't have it.

**So its like you can't treat it like a practice exam because she can't help you answer the problems**

Yeah

**and just like, you probably, have you encountered practice exams, like a full practice exam in your previous classes at all**

I, I'm not sure I don't think I have, at least not in like college classes, like there's study guides that are similar but that's, I don't think it's a full on practice exam

**Ok yeah I just was curious cuz some of the students I've talked to are like, yeah we want the ability to practice solving these problems so sorta playing with the idea there but it sounds like**

I feel like

**Go ahead**

I feel like it'd be way harder to do that with biology, if it was like chemistry I feel like it would be easier because there is a right answer but with a practice exam for like these kind of exams so many of them are like you have to explain why you think that and she's already so far behind on grading I don't think that she would have been able to give us the feedback before the exam to help us

**Gotcha**

Cuz there's like no solid answers on things besides like maybe a few questions that are like, where's the outgroup?

**Right**

But otherwise it's all like you have to give examples and all the comparing and even the drawing question she would have to like look at it individually and say if it's correct or not cuz there's so many options of what answers could be, like, but I definitely wish that it was possible

**Um so so you have kindof a study guide, is there a study guide as part of this class as well**

Well it's like there's study guides for each of the chapters

**Ok**

I don't really use them though because like I don't know I don't like them

**Well it sound like you make your own study guides when you make those like**

Yeah with the papers

**Yeah, So that's I mean that certainly works as an excellent excellent studying tool.  And then I guess we've, we've, we're trying to compare a little bit that prerelease exam versus a study guide so I wonder if you have any experience where you think it was different this way or it was not different this way.**

I feel like for most of the time like, for the classes I took in biology they either didn't provide like an exam study guide at all or it was like very helpful but it was like very clean cut answers kinda things, like if you know the answers to these 50 questions you'll do good on the thing

**Mmm ok**

But it was more like a cumulative thing, so I'm not sure how well study guides would work but I feel like it would get, it would be more like spread out knowledge and you might know the thing but you won't know it well kinda thing

**Ok, like you kinda gloss over it to be like "check, I read this chapter again" but you don't KNOW that you need to understand it**

Yeah it was basically like oh I know this and this and this but you don't know any of the thing well, it's like the Jack of All Trades thing but master of none

**Yeah, nice, well put.  Um yeah like like if you're gonna have that level of, that deep knowledge that you got with the xylem and phloem or like what you were kinda talking about of studying all the like the those chapters in great detail that were on the vague question**

Yeah

**Like instead of spreading out all your knowledge over the five chapters, is that am I getting that right**

Yeah it's like, I still study and like I still know the stuff for the five chapters I don't feel like I fully neglected them for the sake of those exam questions but I definitely didn't focus

**Yeah and that makes sense, yeah cool. Ok. Good stuff. Um, ok. I'm gonna have, I'm not gonna keep you too much longer because it's the end of the day and you've already given a lot of your time and energy and I know that you have lots more work that you need to do Um but I do wanna ask sorta one more tack, um, so as a very dedicated student, the and very skilled and experienced that you've talked about you've shown me that like you're very adept at what you're doing. And so I'm—AND you have a pretty clear pathway of what you wanna do, so I'm wondering is—do you use the exam for something, do YOU use the exam for something personally, like is it useful to you to have an exam to just prepare for or to take?**

Yeah I do feel like the exams are helpful personally just because I like to use the knowledge form the exams on like actual outside life cuz like I'm, it helps me take care of my ferns and make my snake's tanks better temperature wise just like I use if for a lot of small things in life just because I know things better but it's like not everybody wants to use that specific knowledge if it's already possible to do those things

**Sure so it sounds like using, USING the things that you learned for the exam is like, what you get out of the exam?**

Yeah like I definitely feel like I get something out of the exam but there's stress too but like, I do think like it helps me understand just things in life better especially like with my ferns like when I was doing plant form and function and the transpirational pole and I was like oh my soil is too wet and there's not enough oxygen in the soil and the roots are rotting and like I figured out what was wrong with my plants because of it and then I got a humidifier for my ferns So that there would be enough atmospheric humidity so that the fern wouldn't die

**That's awesome**

So that was also from the plant nutrition chapter so I do feel like it helps you apply things

**Helps you apply things**

Especially when you have to like write it out cuz I feel like knowing things and then having to write about something Makes you have like different levels of understanding cuz you have to make yourself sound like understandable

**Ah yeah can you talk about that a little more, kinda like what goes on in your brain as you do that**

Yeah I feel like, like part of when you're writing out like an essay is like it's the same as trying to teach somebody else that isn't taking your class about something and you have to teach it in a way where you're not using words that you learned in those classes so like I'll have my friends listen to me talk about a thing and I'll be like does this make sense because if it's not clear in that then it's probably not clear in my own mind and I'm probably just using a lot of buzzwords that I don't really understand as much as I think I do just because I remember them

**So having to break it down and explain it to someone else either in real life or pretending to explain it to someone else on the exam is like solidifying your knowledge**

Yeah I definitely think so

**Yeah I think so too haha. Excellent and um, let's see. Ok I'm trying to think if there's anything else I should ask, I feel like I've gotten a lot out of this conversation and our last conversation as well, so this is um--anything you can think of that I should know or an idea you had about how we could improve the course or the exam Or the process or like something that you wish had been included**

Mmmm that's a hard question um.

**Or something you wish hadn't been included or something you wanna change**

Well the, I feel like I don't, I don't think there's much that can be changed in the way of making it better or like just because like all the things that somebody would normally complain about are things that are hard to realize would make it harder, like less chapters on the exams means more exams, which means more stress, or like… so I don't think there's anything that could be changed personally just cuz I feel like it's a trade off and I feel like all the tradeoffs would be bad

**Sure yeah that's fair, yeah I mean ultimately school is hard, learning is exhausting, exams are stressful and there's not really much we can do about that. Ok well good to know. Um. Ok so like the like the very basics of the research we're doing, I'll just ask you a yes or no question, if you had the option to keep the prerelease exam or do a more traditional study guide or outline, would you do the preexam or would you do the study guide**

I think I'd do the preexam personally

**Ok**

Just because the study guides usually never really helped me that much

**Ok**

So personally.

**Great, great and that's the sense I've sorta been getting from our conversation too so I'm glad that's your answer cuz that means I've been understanding correctly. So unless you have any last thoughts […]Starbucks[…]so yeah thanks again it's really it's terrific to talk to motivated students**

No problem

**Huh?**

Oh I said no problem, I'm just glad that I could get included cuz I feel like my opinions are really rare in this class

**Well I hope they're not too rare cuz they're awesome. But it's really valuable hearing from student who can ….**

### ***Interview #11 June 17 2021 @CC site (Leah Lily interviewer = bold)***

**We're trying to talk to students about […] Your experience is really important and […] so how did it go for you?**

Um it was good, um I loved the exam taking in person a lot more, um. Especially on the second exam we were able to meet in actual classrooms and so this kinda gave us a space to where we could meet beforehand and then we could you know go to the classrooms after as opposed to the first one We just had it on the first level kinda like out in the open. Um and so we didn't really have like a space that we could study beforehand so we just kinda walked into it um not really having a conversation with anyone else or um practicing some of the questions that kinda thing. Um. The one thing, this specific course it's been a lot different for me because a lot of the information we had to learn on our own because all the time in person was dedicated to our research projects. And so I'm a very kinda visual person I like someone to be able to explain it to me and give me a diagram and um although the class was Tuesday Thursdays the only time the teacher actually met for questions was on Wednesdays and I work Wednesdays so I didn't really get that One on one attention—not one on one but I didn't get that classroom attention that I wanted. Um but overall I did have a lot of support from my other classmates, um, we actually have a discord group so we can you know communicate with each other, we can ask questions and kinda meet in study sessions so that was really helpful for me

**Great. Um how did you prepare for the exam?**

Uh so the teacher gave us a really good study guide on—and then I personally type out all the chapters so I have a binder—I'm not sure if it's actually on this table I think I moved it but, oh no it's right here, hold on. Ok so I have this binder, and it has the study guide in it, um I have like a diagram that we had to learn um, and then this I got actually from the chapter and then it kinda goes into like some of the questions and then my answer um and then it has like the tabs here for I don't know if you can see that the tabs for each chapter and so I typed out The notes for each chapter and I post them to the discord group and then I just studied from that so

**Wow that's a lot of dedication**

Yeah haha

**So the questions that you answered, you said, in your binder, where did—are those questions from the chapter?**

Those were questions that she specifically gave us on the study guide, and then I would just write out all the information I need to know about, you know the different things within the question. She would withhold—she withheld a little bit of the information so we didn't have the full question there but for the majority of it I was able to have it listed and then go back to the chapters to try to fill in the blanks

**Ok so did you successfully fill in the blanks?**

Um there were a couple questions where I had an idea that was coming and there was a couple questions that I totally missed. So yeah

**Um all right, that,s well that 's a pretty good average for fill in the blank**

Yeah

**How, did you study differently for the questions where you were like oh I figured this one out versus the one where you couldn't figure it out**

Um overall I kindof I mainly focused on the information given and then I would touch back on the chapters um and I would use more of like the larger subjects in the chapter that weren't already listed on the study guide to try to fill in the you know fill in the blanks. There was one specific where it was a drawing question and we had to draw out that phylogeny, that chart I just showed you that was handwritten, and it asked for what were I think they were monocots and dicots and I just did not even see that coming, um I should have, it was kindof a common thing within the chapters but I didn't actually think that that was gonna be a question so that one I completely missed.

**Bummer. So that study guide that you're talking about, had you encountered a study guide like that before?**

No not really. Um Ive had study guides where it tells you to study specific chapte—er not chapters, certain terms, certain areas within the chapters, um some have like just given a whole bunch of questions and I've been able to study from those but the fill in the blank kinda threw me off because we were given basically the questions but then there was this element that was missing and you had no idea, you knew, She let us know what chapter the fill in the blank was from but the chapters are so extensive that it was kinda hard to piece it together.

**Mmhmm what, do you have a preferred type of all those different study guides that you've encountered**

Um if I had it my way I would just have all of the questions and just you know be able to type out the answers and study that way, I like that, I also like being given like a list of specific topics to cover um, being a very visual person I don't like to feel like I'm missing something so to have all areas needed to be studied listed out in front of me is very beneficial

**Yeah that makes sense just like like an inventory of just like you should know all these topics?**

Yes

**And is that, I imagine that's useful not just for the exam but also for just like, what did we learn in this course**

Yeah yeah also going through each chapter there's just so much knowledge that having a list of like what's important to the instructor for that quarter is super essential I feel like, just so you know what to focus on more of um

**Oh sorry go ahead**

Uh no I'm I'm good haha

**I was wondering if knowing what to focus on, what the instructor thinks is important is that important for the exam or for the course in general**

Um I feel like the course in general just because if we have an understanding what topics are going to be emphasized in the beginning we can put more energy and more time into those topics as we go as opposed to trying to cram it all at the last minute

**Yeah totally cool. Well so I'm curious about, about you, it seems like so you have this study method that you've been using for a while now are you a sophomore junior senior**

Um I am …

**You're a student!**

Haha I am probably sophomore now? I'm almost done with my AA, I have I think two no three more quarters including summer quarter and then I'll have my AA. Um and then I'm not sure if I'm going for a bachelors of science but the end goal is my doctor of chiropractic

**All right that's very cool**

Yeah

So you have a vision of what you wanna do

Yes yeah it's gonna take some time to get there but I have a lot of people helping me out so that's helpful

**Yeah that's great. And I guess for this course, so biology seems pretty relevant but I'm not sure, does plant biology, did that kinda fit into your path for yourself or was that sorta out of the blue**

Um so plant biology was a requirement for my associate's, but it-- I also love biology um and so I was kindof excited to take it and it gave me the chance to go to the farm and like be outdoors around people for once, which being my, it was 5th quarter at this school it was nice to finally actually see someone face to face for a class so

**Yeah oh wow you've just been doing this all through covid, that's really hard**

Yeah haha

**Well we're almost through it now, and you're almost done so that's great**

Yeah

**Very cool. All right so I guess, I guess, something I was curious about is like so it sounds like you really enjoyed the course and that's wonderful, and you enjoy the learning part of it and so I'm wondering From this course, did you get any skills that are sort of like generally applied to what you think you're gonna need a s a chiropractor?**

Overall this quarter it was a lot different from animal biology in that I had a very opinionated research group and so having to navigate around everyone's opinions, that was definitely a struggle and I feel like I gained a lot of skills just trying to in a sense make everyone happy but at the same time you know to put your foot down and be like you know You have kinda the unpopular opinion and we need to come to a conclusion um so let's try to find like a a middle ground, and come to a conclusion so we can move on with our experiment. That was something that was much needed this quarter. Um. And then also trying to manage a schedule um that included you know work, and going to the farm, and working from home, that was um something too that I'll uh I'll definitely uh it I learned a lot for sure haha, chiropractors they have a fairly high paced schedule and so having to manage so many different aspects of my life although overwhelming at some points You learn to kindof get through it and um that was very helpful for me too.

**Yeah that sounds like excellent skills to have and it's great that you recognize that you know you're gonna need teamwork ahah**

Yeah haha

**And scheduling and balance and all those things, that's great. Well I'm escited for you to realize your career and realize your future that's very cool**

Yeah

**So I guess I'll go back to the exams again, you mentioned a discord, and like you put your own notes on there to share with people. Do you feel like you get anything back from um the rest of your classmates**

Um the only thing really that I get back I feel like is you know people's reactions to the notes, a lot of them are really thankful to have that material, some people they have the physical book but I don't, and for me to try to look at a screen and research stuff and learn stuff it wasn't working for this course. I kinda tried something similar with chemistry notes but It was so hard to type out every single equation that I just gave up on that and stuck with handwritten notes but I just felt like this course, and animal biology was going to be so tough in itself that I might as well try to help others out as well, so I just did that, I felt like it would be almost a little selfish to you know hang on to them when I put so much work into then so to be able to share that was just another benefit And I know we, we kept the same discord from animal biology to plant biology um but not everyone in this plant biology was in our last animal biology, so all the notes are there for them, um so I kinda in the back of my head hope that you know people will keep just being added to the discord and the notes and resources and questions will already be there for everyone so they don't you know have to go through it all by themselves

**Wow that's really generous of you, that's super cool**

Thanks

**Um what, did students give you feedback, what did they say about having someone else's notes to study from.**

Um so I had one student in particular he yeah last quarter I had gotten through like ¾ of the notes and then things got busy, my partner's uncle passed away and we had to go to Illinois to pick up a lot of his stuff and stuff and road trip it back in the middle of exam week

**Oh no**

And so during that time I stopped posting the notes and he had expressed at the end of the quarter just how, how much he gained from those notes and he thanked me and everything. Um I started the notes back up this quarter and I had so many more people reacting to them, and I think it was their way of showing that they didn't underappreciate my notes and that they were being used so that was really kinda nice to see.

**Yeah so ok I'm gonna dig a little more about your, like YOUR preparation now. So you write up these extensive notes, and you post them for your classmates which is nice, but do you study from them as well.**

I do yeah. A lot of the times before I have a quiz I'll try to condense them even further um a lot of times a chapter I can have up to 18-24 notes typed on them Um and so I try to condense it down to about two pages with the main topics and then study from those right before having a quiz um and then for exams I just go through everything, I don't condense anything down and I just yeah go through everything, try to find where the questions relate to the chapters, um so that I can kinda pull information from there, when writing an answer to each question. And so yeah um. And I print all of that out so I have it on hand and I don't have to look on the computer for it so.

**So it sounds like you have your answer basically done before the exam, is that right**

Yeah

**Did that work for you? Because you didn't know the entire question, but it sounds like it was like half and half**

Yeah it worked really well for me. In high school I took a public speaking class and so I kindof act as if I'm giving a presentation on the question and you know I start to get nervous every time I look down to try to find more information about it, so by doing that I sort of get it into my head, Um and learn, learn the answers that way.

**That's very cool huh. Um would, so would you say that's like more memorization**

Yeah a little bit, it's overall trying to get you know the facts in my head especially for the compare and contrast questions, um getting the facts in your head and then you know as soon as you figure out what questions you have, she let us have scratch paper so I could write down those ideas and then I could start to incorporate them into an essay

**Mmmhhmm and those were like, like the ideas that you had memorized based on your notes**

Mmhmm

**And just kinda jotted down to remind you**

Yeah

**Awesome yeah that's a good exam process I've always been told like write a little outline for yourself, like that's really good.**

Yeah

**Um all right so it sounds like this has been an effective method for you, it sounds like you work really hard and you're really dedicated to your learning […] difference in the world so you know you're being very inspiring to me. But I am going to push back a little bit. So it sounds like you do a lot of memorization. Is there a chance for you to practice applying what you've learned?**

Right now in my day to day life there is not a whole lot just because I'm not really involved with plants. In my house I don't know if it's the water or whatnot but we can't keep plants alive here

**Oh no**

We've tried a whole bunch of stuff including tryint to collect water outside and it hasn't worked for us so far, but I do have a lot of fish in my house and so um kinda learning how you know their systems work through animal biology has been really cool and it gives me a better understanding you know of how they work so yeah.

**Yeah yeah and I guess let's see that sounds like in your own life. Was there a chance to apply it you know, you took this course and you had this knowledge that came from the course ,was there a chance to poke at it a little more or a chance to apply it as a real life scenario as part of the course?**

Oh um yeah we had a, each quarter we had a research project so last quarter we had mealworms we were testing to see if they had specific color preferences, so we were able to do that, and then this quarter we did a research paper on um a pollinator's preference towards some native plants so that was cool too um. There were a lot of setbacks this quarter with our research project but it just kinda gave you that much more knowledge you kjow how to navigate around those issues to try to get a good presentation. Sorry I don't know if you heard my dog, the neighbor dog must have just gone out so

**Yeah I wanna play too**

Yeah he's tearing up our grass over there

**Oh nw, ah. Ok that's right, so you had that research project. People say that was really like very rewarding**

Mmmhmm

**Seemed like real life kinda thing**

Yeah yeah it's been awesome, we have so much information we have I think like 17 pages now for our final so uh we've really gone in depth with it that's for sure

**That's awesome. Ok I guess what I'm curious about now is it sounds like, it sounds like for you at least the exam was more about memorizing the knowledge rather than applying it. Is that right?**

Yeah last quarter we did have a question that gave us a scenario and how we would navigate around that. This exam it was more of like compare these two things, draw this, interpret this graph, um, there wasn't a lot of connections into the real world it was more just information regarding the chapters

**Ok. Um is do you prefer one or the other kind of question**

I prefer real world scenarios because I feel like there's a couple different ways to go about it, and so you can kinda better portray your knowledge about the subject instead of feeling like you should just memorize a whole bunch of stuff. So it's actually applying the information

**Yeah. I guess I'm also curious like if you had, if you had a que-- like it sounds like you had those questions partially released to you, and if you had a question that was like take this real world**

**scenario, you're gonna have to dig into this on the exam We're gonna ask you questions about it, does that sound like a better question or a more useful question**

Yeah I'd actually love that because then you'd actually have to go about and figure out all of the concepts that would tie into that specific population and it kinda it pokes at what the question is but it doesn't give it to you in the sense that you feel like you need to memorize those specific concepts That are gonna be on the question.

**Mm hmm yeah that makes sense.  Hmmm.  Would you study differently for a question like that oh yeah tell me about that**

Haha so if it was just a general population question you'd have to learn just so much more about I feel like the chapters and what might be going on with a specific population whereas the questions this quarter they just gave you a majority of what was gonna be on the question, so I definitely feel like if it was regarding a population I'd have to study quite a bit more and dig deeper into the chapters than I did this quarter.

**Mmm so my pushback would be, that sounds like a lot of extra work**

Yeah haha

**But for you it sounds like it would be worth it? You're a hard worker but**

Um so I think that in this specific class because so much is gonna be, or was dedicated to the research project I feel like it was kindof a good thing to have questions that were pretty straightforward, um last quarter um we didn't have so much tied into the research project it was just kinda a smaller thing than you know this quarter, um so it makes sense that the questions were so much more in depth last quarter so I think there's a little bit of give and take in regards to you know the chapters versus application of the data so, or the information.

**Yeah. Which class did you like better**

Uh I actually liked last class a little bit more um I think it's primarily because the group that I had to work with last quarter Um everyone just worked really well together and it just made the class enjoyable, um this quarter having a little bit more um feedback from a lot of our, our team members it kindof made me resent the course you know sometimes um. We just had, we had so many opinions and we had a lot of people that were very outspoken during the quarter and I think that was a major, a major negative, ha, part to the course.

**Yeah that makes it really hard.  Um I wonder if do you think if you had, say, like the previous quarter if you had the more in depth questions versus your more in depth research project does the group work um issue come up in around exam questions as well or is that just the research**

Uh it's different from the exam, so nothing within the research projects showed up on the exam

**Oh ok I see. Um.  Hmmm.  Have you, have you ever tried studying with other people? Not just sharing your notes but, oh yeah ok**

So there was, I think it was our first biology course in the series, um we had actually posted an online document that shared all of the questions that was gonna be on the exam and I think there was like 45 questions that could be on the exam, and just everyone worked together to tackle all the questions and it turned out to be really good, um it was really good teamwork on everyone's part um and then to actually you know be able to look at a question and be like oh no that answer you know is only half right and be able to adjust that and explain why it wasn't correct before gave us so much more insight Um and it gave us you know other peoples' opinions and um so much just factual data uh within the chapters that it turned out to be a really good um final exam so um I felt very prepared when everyone is able to chip in.  Previously I, like in high school like before this whole covid school thing we had a lot of study sessions and that was always nice too so.

**Do you intend to keep that up once we're allowed to after covid**

Yes ahah.  Yeah, When I eventually go to chiropractic school I'll have to move out of state because there's not a program here in WA so I hope to use as many resources there as possible when it comes to studying For you know there's an exam, if I end up going to Portland there's an exam after semester 5 so you really have to learn all of the information and that exam determines if you get licensed as a chiropractor so I'm going to be using as many resources as I can whenever needed so

**All right.  What , I'm trying to imagine like all these resources that there could be, would there be some that you would prefer that you're really hoping they would have available?**

Um I think just instructor availability so um as much information as I can get from the higher ups, I'm definitely gonna want that, being able to study with others just to make sure there's not a part of a specific question that I'm missing is gonna be huge as well, and of course quiet places to actually be able to escape to and study the material so.

**Yeah.  So am I, maybe I missed it but did you study in a group for the 213 exam?**

Uh for the second exam I had a service learning from I think it was 10 to noon that day um I could be wrong it might have been 9 to 11 but my exam wasn't until 2:50 so I kindof hung out with studying and I started to notice some people trickling into the building that were the later biology group so I ended up you know going into the building and they had set up camp in this little lobby area and so we were able to sit down and start bouncing information off of each other and asking different questions about the questions and what we think the fill in the blanks might be, just kinda sharing information right before the exam and that just gave me so much confidence as to how much I know going into the exam so. Even during service learning we were just on the Edmonds college farm and pulling weeds so we were able to talk about the different questions as well, and try to list off as many different species of plants in our little phylogeny as we can so that was really fun haha

**That's great, that sounds like it kinda just happened, yeah, you didn't really plan it**

Yeah

**Wow that's really cool**

Yeah

**And it sounds like, it sounds like the discussion that was trying to fill in the blanks was like really productive for you**

It was yeah it brought up a whole lot of concepts that I didn't think about and to be able to have just a little bit of information Right before the test in case it came out uh was super beneficial to me because a couple people they mentioned something that I hadn't even studied before, it didn't end up on the exam but at least I had a little bit of knowledge before stepping into the classroom

**Yeah that's really cool, it's like everyone's a little bit different so everyone's going to guess a little bit differently**

Yeah

**If you had the opportunity to take an exam like this again where you had the partially released questions before, now having tried that once, how would you go about studying from it, like knowing what you know now?**

I would probably go a lot more in depth into answering the questions in my notes um just include a part that would have those fill in the blanks um because I really just kindof reviewed the chapters, I didn't write down anything and I learn best when I'm you know typing or engaged in learning um and so I think I would try to do a little bit more of that and studying from like a smaller amount of information as opposed to all of my notes on the whole chapter so

**So you'd like use that prereleased part of the exam to be like what do I need to focus on because what COULD be in the blanks**

Exactly

**Ok um ok and how would you guess on what's gonna be in the blanks, like specifically look through the chapters or**

Yeah I'd probably go back through my chapter notes and try to find out what concepts tied back into what's being asked in the question or how they relate to the question and try to go about it that way um all of the questions, it made sense for everything to kinda follow that pathway when the blank was such a small part of it so it didn't make sense after you know taking the exam for that to have been a concept that didn't tie into the beginning of the question. Um and that was demonstrated on the exams. Every fill in the blank had something to do with what was previously said so.

**That makes sense, so it's kinda like you're building connections with like oh these are the things I was given, and they tie into THESE other things so I study THESE things, like that?**

Yeah haha

**Awesome I wonder how I'm gonna write down [gesture] THESE things.**

Haha

**Yeah sometimes transcribing is hard haha. Thank you that's really useful, I'm conscious of time I don't want to keep you here for too long. Is there anything you feel I should be asking to find out you know how do we make this work**

Um overall I feel like a lot of things that happened over this quarter were already discussed, I don't feel like there's a lot that was really missing, yeah so

**Good to know, well um I guess I'll wrap it up then thank you so much for your time and your effort both for this focus group […] starbucks […] any last ideas?**

No I think that's everything
