## Supplementary material for "Public exams decrease anxiety and facilitate deeper conceptual thinking": Public Exams Supplement 2 Interview Coding

| Final Code: | Definition: | Example Quotes: (*Interview / Page / Line* ) |
|---|---|---|
| Language issues<br><br>(This code was one of the original codes explored in this research.) | Students talking about what they've learned about taking exams, especially in regards to understanding and deciphering the language of what is on an exam. | ● "I would have spent more time on the public exam if they were scenarios because I feel like it would have been more directional with my studying, for example the one really long reading question that [subject] was alluding to was about pretty much like MDMA being cleared out in the liver instead of alcohol being cleared out in the liver, but just that reading was like, yeah, I had to spend a whole minute or two reading and digesting those big words, but then I knew, well, they're going to ask me a question about how the liver cleans out stuff and what might happen if something's wrong with that and so that gave me a direction, so I appreciated that because that did lead to me studying about the liver clearing out toxins and leading to me getting the right answer on the test, so that was nice." *1.7.269*<br>● "Already exams are like, ok, the graph, it's about the graph, I'm not really sure but it's about a graph, it's like ok, if this happens what will happen to the graph? So even though we got questions like this like in our lecture, it's different from this because the questions are structured differently, like, I remember the questions are structured in ways that are like oh, this is testing my English" *3.1.27* |
| Directing to core concepts<br><br>(This code was one of the original codes explored in this research.) | Impact of public exam system on efficiency, completeness or speed of finding and studying information for the exam.<br><br>Comparisons to other courses using study guides is a relevant example here. | ● "When I think of skills I think of it less as knowledge and more as like ability to like get information kinda thing, um, I don't think of it as like I can memorize all the Krebs cycle stuff, or um I can like list all these things off of the top of my head, I think it's more like knowing who to talk to, like at least in my job right now you gotta know who you wanna talk to in terms of getting prescriptions for patients, or like talking to the doctor to get some sort of like change in a patients' medication, um yeah I, I think like the skills are like communication skills, like the collaborative skills, are the skills that I will take away<br>● "I think I do like how he structures it, how some things he shows and then some he doesn't |

| | | |
|---|---|---|
| | | because it really gets you thinking about I do agree with how you kindof have to master that subject in order to answer the question" 1.2.68<br>● "he kinda even like puts the drugs on it like, what was it, cyclobenzaprine, or like some like drug that I've that I've sold at, as my pharmacy assistant job. Um I think it's just like it's valuable because he's going through all these steps, and going through all the ways that you can break it and all the ways that you can save it kinda thing that you can fix it. And he sometimes puts drugs in there and I'm like wow that's up my alley, I'm going to need all that stuff." 6.7.272<br>● "I do really like the way we did this exam because even in the career no boss is gonna ask you, they're not going to surprise you with something I guess in a sense where you have, You haev the opportunity to work on skills to build skills if I wanted to move up within a position or company or whatever like I have an understanding of what's expected of me in that position and the opportunity to learn and to grow and to take classes and get that experience or knowledge that I need to be competitive to make that next step. And so I liked the exam, I mean I have a hard time with tests in general just because no job is necessarily going to- you always have the ability especially in our day to grab your phone and google something, so You have the information accessible to you and yes there is the need in a program like this to show that you have like an understanding and you don't always have to go back to the information to get that, and so I liked that because I felt like this balanced it out. Where I had the information, like the graph that was on the outline is the graph that was going to be on the test, the phylogeny on the outline was the phylogeny on the test, so I had that ability to have that information, See it ahead of time, google, read, whatever I needed to do to have that information and then be able to take the test on it and so I felt like it was a much more realistic view of what life looks like." 9.6.200 |

| | | |
|---|---|---|
| Deepening thought<br><br>(This code was one of the original codes explored in this research.) | Includes references to types of thinking, creativity, enjoyment, and higher-order thinking like metacognition and synthesis.<br><br>As part of metacognition and lack of creativity, discussion of memorization/"regurgitation" in regards to the exam process is here. | ● "When I was starting out I thought it's more like memorization based, or like, you know, it's not really like how applying, like I don't, like, I don't relate [Physiology course] to like having to apply a lot of things, like having to apply what it is; it's an introductory course to our physiology, I was like, ok, so I have to know where the liver is, what is the function and stuff but no, it turns out, like ok, if this happens what will happen to this, it had to like, it was a whole different way, I have to think in a whole different way, I have to make sure that the foundation that I have for the function of this particular organs and the enzymes, I have to really know what they're doing in order to like apply it to the questions that are like the exams." *3.1.20*<br>● "I've never done this before; a professor has never partially given me an exam, so it was a new thing for me, so I was trying to figure out what kind of study habits I need to do in order to do well on the test. So I kindof, I would look at the public exam by myself, go through each question, like what it could be, and then I'd go back to my notes or rewatch a couple lectures or two, and kindof not find the answer but try to figure out what he could be talking about or what kind of topic he is trying to figure out in kindof a way So it's kindof an interesting, like, study kinda session I had over the last two weeks. *1.5.166* |
| Authentic engagement<br><br>(This code was one of the original codes explored in this research.) | Students referencing or demonstrating trust, enfranchisement/involvement with the exam process, empowerment, or negatives/lack of same.<br><br>Students stating that they felt like they were a part of the decision-making teaching team in regards to exams would fit in this code. | ● "He, like, explicitly says exams are horrible at gauging your progress but they're what we have, and he makes an effort to revise exams, he releases it ahead so we can look through it and see if some wording is weird and so he's, like, obviously making an effort so we can't, we can't really—it's like really hard to hate on him as a professor because he's doing all the things that we want him to do, he's, like, hearing us basically, he's like I hear you, like, exams are hard and so I'm gonna, like, you're gonna help me with it." *6.6.234*<br>● "I would hope that in the future when we take more in person exams that the structure will kinda change to sort of match our understanding of the material, or at least the way that we're sorta |

| | | |
|---|---|---|
| | | studying it, because we're kinda just studying it all together to sorta memorize and understand all of it but we're only getting tested on certain aspects of it, so it's kinda like, are we wasting our time with studying everything, could there be a point to where we are being testing on what exactly that we're studying or memorizing?" *7.8.290* |
| Collaboration<br><br>(This code was added to the codebook during iterative analysis of transcripts.) | Students referencing collaboration skills, practice, or real-life relevance of it in regards to the exam.<br><br>This code includes study groups, class-based research groups, teamwork, and/or communication. | ● "We all kinda problem solve and work through it together, which has been really helpful, so there's just a ton of different ideas to flush out one question and the possible answers and then just go through that. So I think a group study for the public exam is best. I think by myself I'd be pretty overwhelmed and there's just-- again, there's just so much so I feel like having other people working on the public exam with you really, really helps me." *4.2.62*<br>● "Everyone worked together to tackle all the questions and it turned out to be really good, um, it was really good teamwork on everyone's part. And then to actually, you know, be able to look at a question and be like, oh no, that answer is only half right, and be able to adjust that and explain why it wasn't correct before, gave us so much more insight. And it gave us, you know, other peoples' opinions and so much just factual data within the chapters that it turned out to be a really good, um, final exam so I felt very prepared when everyone is able to chip in." *11.7.253* |
| Anxiety or Confidence<br><br>(This code was added to the codebook during iterative analysis of transcripts.) | Emotions around exams, especially those that impact feels of ability or inability to do the work and succeed on exam-based outcomes. | ● "For some people I know with test anxiety it was more like being absolutely sure you know all the material, but it can be overwhelming because you never feel like you know all the stuff that you need to know. But that's why I sorta go overboard with my studying and try to, like, start really early because I want to make sure there's like no possibility that I'm not gonna know what's on there but you never know til It's in front of you." *8.5.140*<br>● "I definitely feel how it can be harder with you seeing the questions before, and I see it in, like, a new light and you sorta have to re-flush out all the answers again. I feel like that can sometimes be |

| | | |
|---|---|---|
| | | especially challenging for me, like ok, I feel like I've prepared for this as well as I can and now it's something and I just gotta do the whole thing over again which is sometimes pretty hard. But then on the flip side of that it is nice to be like, ok, I know I recognize this and I can do the same process of what I've done before, so I do kinda feel the love-hate with the public exam but I do think it's helpful in the end." *4.4.145* |
