## Supplementary material for "Public exams decrease anxiety and facilitate deeper conceptual thinking": Public Exams Supplement 3 Survey Item Coding

| Code: | Definition: | Positive Example: | Negative Example: | Non-matching Example: |
|---|---|---|---|---|
| Code #1: Language issues | Understanding what was necessary to do in order to answer the questions | Yes, I really like being able to see the exams ahead of time because they help guide me to what I should study and also make me less stressed because I'm not worrying about whether the test might be written poorly or if there are other issues like that. I can focus more on the actual material. | Yes, I like the style of the exam. Wording sometimes is not the best, but overall it is good! | no, i think the exam is difficult for me to understand. I have a hard time finding a way to study the material |
| Code #2: Directing to core concepts | Understanding what was going to be on the test and what to study | Yes, I really enjoyed the public exam which narrowed down what we had to study for without really limiting it to us only focusing on one small topic, that way we need to know all the chapters for the exam. It made me less nervous and stressed about the exam in this covid situation, and helped me better prep for exams. I wished all the exams were in this format. | No, they were very hard to study for and were not an accurate representation of what we learned in the course. They targeted very specific material which made learning other stuff useless. | Yes, i felt there was a better way to study for this style of exam. |
| Code #3: Deepening thought | Applying knowledge, understanding rather than memorizing, something being | Yes, I enjoy having the questions beforehand so that we know what to focus on with studying. Being able | I have never taken a public exam before, but the style most definitely worked for me. I enjoyed | The first exam wasn't as specific when we had the practice available, but the second exam was much |

| | | | | |
|---|---|---|---|---|
| | “interesting” or prompting curiosity | to analyze the diagrams beforehand is also helpful. | the structure of the exam because I knew what to expect and what kind of questions will be asked. Knowing and seeing the exam beforehand allowed me to adequately prepare academically and mentally to do my best while taking the stress and the 'fear of the unknown' out of my mind (for the most part). Comparing to other exams (non-public), I was able to fully answer each question confidentiality while having fewer questions that I did not know fully how to answer. On the downside, I know some individuals only study the materials explicitly stated on the public exam without studying the whole material covered within the course. Consequently, I personally recommend this type of structured exams for all STEM courses for myself. However, to fully experience the positive advantages of a public exam | better. The exam 2 worked for me better than the first because it was kind of in terms of the question that was asked. |

| | | | | |
|---|---|---|---|---|
| | | | compared to a regular exam, an individual must know how to study for a public exam. But that is the responsibility of the student regardless of the style of the exam. | |
| Code #4: Authentic engagement | Feeling heard, participating in the assessment process, fairness of evaluation | Yes. I appreciate the multiple choice style of the questions. I feel that I was given the materials required to be prepared for those exams, so it was all about my ability to actually put the work in and study those topics. | I'm not sure if I'm overthinking it, but perhaps having more public exam questions have caused instructors to up the difficulty of the exam? If that is the case, I would much rather have less public exam questions and exam questions that focus more on the big picture of the biology we are learning rather than the small details and wordings of the questions. | I have noticed that it only works for me when I work with other people in study sessions. I try to study on my own I have a more difficult time understanding the material, which is something quite new to me since I am used to studying on my own. But overall I like it. |
| Code #5: Collaboratio n | Students working with each other to prepare, or referencing collaborative skills as real-world needs | Yes, it does. It makes me feel less alone in the process and we can travel down many avenues discussing questions in study groups, which makes for a good review session. It is also much less stressful too. | (none) | It worked for me. I can guide myself what the exam will look like. |
| Code #6: Anxiety or Confidence | Emotions, including surprise | Yes. As someone who gets nervous taking tests this test format has made me more confident | Yes, I think it worked well for me because I felt like I was given everything I needed | I liked how I could jump from question to question and double check my answers however a |

| | | | | |
|---|---|---|---|---|
| | | in my knowledge going into tests. I enjoy the public exam questions because it gives me an idea of the topics that I need to spend more time studying which also helps me find gaps in areas of knowledge. | to study successfully, it was up to me how well that was going to be. The unknown part of the question added a lot of stress but I feel like this type of testing style is beneficial to everyone because I can talk to my peers about what is confusing or not. | lot of the questions had too many answers that could be perceived as very similar, or even the same. |
| Unspecified reason | The response given did not explain the student's opinion | Yes they did. I liked how they worked, didn't like some questions though. | I think i do better with blind exams because I pick up on material quicker than most | Yes - I felt more prepared for this exam than the last two, mostly because I knew where to focus my studies and understanding. |
| Code #7: Not relevant to the public exam system | Student's reason given for their opinion was not related to the public exam or similar practice that could be included in the public exam system | Yes，I like that it add some short answer questions I think it will boost my point a little bit more. | Yes, it worked because it is all multiple choice, but it is still too hard. | I think it did because it gave me a sense of what to study for. It was still challenging because we couldn't see most of the problems, but it allowed me to think in different ways I wouldn't have otherwise been able to. |
